# Small molecule SWELL1-LRRC8 complex induction improves glycemic control and nonalcoholic fatty liver disease in murine Type 2 diabetes

**DOI:** 10.1101/2021.02.28.432901

**Authors:** Susheel K. Gunasekar, Litao Xie, Pratik R. Chheda, Chen Kang, David M. Kern, Chau My-Ta, Ashutosh Kumar, Joshua Maurer, Eva E. Gerber, Wojciech J. Grzesik, Macaulay Elliot-Hudson, Yanhui Zhang, Chaitanya A. Kulkarni, Isaac Samuel, Jessica K. Smith, Peter Nau, Yumi Imai, Ryan D. Sheldon, Eric B. Taylor, Daniel J. Lerner, Andrew W. Norris, Stephen G. Brohawn, Robert Kerns, Rajan Sah

## Abstract

Type 2 diabetes (T2D) is associated with insulin resistance, impaired insulin secretion from the pancreatic β-cell, and nonalcoholic fatty liver disease (NAFLD). SWELL1 (LRRC8a) ablation impairs adipose and skeletal muscle insulin-pAKT2 signaling, β-cell insulin secretion and glycemic control - suggesting that SWELL1-LRRC8 complex dysfunction contributes to T2D pathogenesis. Here, we show that I_Cl,SWELL_ and SWELL1 protein are reduced in adipose and β-cells in murine and human T2D. Combining cryo-electron microscopy, molecular docking, medicinal chemistry, and functional studies, we define a structure activity relationship to rationally-designed active derivatives (SN-40X) of a SWELL1 channel inhibitor (DCPIB/SN-401), that bind the SWELL1-LRRC8 hexameric complex, restore SWELL1-LRRC8 protein, plasma membrane trafficking, signaling and islet insulin secretion via SWELL1-dependent mechanisms. *In vivo*, SN-401 and active SN-40X compounds restore glycemic control and prevents NAFLD by improving insulin-sensitivity and insulin secretion in murine T2D. These findings demonstrate that small molecule SWELL1 modulators restore SWELL1-dependent insulin-sensitivity and insulin secretion in T2D and may represent a first-in-class therapeutic approach for T2D and NAFLD.

## Introduction

Type 2 diabetes mellitus (T2D) is a globally ubiquitous metabolic disease characterized by hyperglycemia that is caused by reduced insulin sensitivity in target tissues and impaired insulin secretion from pancreatic β-cells^1–3^. T2D accounts for 90-95% of all diabetes mellitus in the US, or about 24 M people^4^. It is associated with increased risk of cardiovascular disease, renal disease, liver disease, cancer, and infection and a hazard ratio for all-cause mortality of 1.80 compared to patients without T2D ^5, 6^. The cost of medical care for patients with diabetes is 2.3-fold the cost in non-diabetics. In 2017, the direct medical cost of diabetes in the US was $237B^7^.

There are at least ten distinct classes of medications approved to treat T2D: sulfonylureas, meglitinides, amylin mimetics, biguanides, alpha-glucosidase inhibitors, thiazolidinediones, glucagon-like peptide-1 analogs (GLP-1a), dipeptidyl peptidase-4 inhibitors (DPPi), sodium-glucose co-transporter (SGLT)-2 inhibitors (SGLT2i), and insulin. Despite this diverse array of T2D medications, there are several reasons why new medications for T2D are needed. First, cardiovascular disease (CVD) is the leading cause of death in diabetics ^8, 9^, and although newer T2D medications like SGLT2i and GLP-1a effect a reduction in CVD mortality, significant residual CVD mortality remains ^10^, which presents a therapeutic opportunity for T2D medications with novel mechanisms of action. Second, 25-33% of T2D patients have inadequate glycemic control, with HbA1c levels above guideline recommendations ^6, 11–14^. This poor glucose control is associated with increasing risk of death from vascular causes, non-vascular causes and cancer ^8^. Third, T2D medication-induced hypoglycemia remains a significant problem for patients with T2D, especially with patients on multiple T2D medications^4, 15, 16^. For all these reasons, there remains sustained interest in developing new T2D and metabolic syndrome therapeutics, especially with novel mechanisms of action ^17^.

*SWELL1 or LRRC8a* (Leucine-Rich Repeat Containing Protein 8a) encodes a transmembrane protein first described in 2003 as the site of a balanced translocation in an immunodeficient child with agammaglobulinemia and absent B-cells^18, 19^. Subsequent work revealed that this condition was caused by impaired SWELL1-dependent GRB2-PI3K-AKT signaling in lymphocytes, resulting in a developmental block in lymphocyte differentiation^20^. So, for about a decade, SWELL1 was considered a membrane protein that regulates PI3K-AKT mediated lymphocyte function^18, 19^, and it was not until 2014 that SWELL1/LRRC8a was discovered to also form an essential component of the volume-regulated anion channel (VRAC)^21, 22^, forming hetero-hexamers with LRRC8b-e^22, 23^. Therefore, historically, the SWELL1-LRRC8 complex was first described as a membrane protein that participated in non-ion channel mediated protein-protein signaling (non-conductive signaling) and then later found to form an ion channel complex with ion conductive signaling properties. Indeed, prior work highlights each of these modes of SWELL1-LRRC8 channel complex signaling. We showed previously SWELL1 to mediate insulin-PI3K-AKT signaling in adipocytes and skeletal muscle via non-conductive signaling mechanisms, and thereby regulates insulin-sensitivity, by modulating GRB2 signaling^24–26^. Also, we and others showed SWELL1-LRRC8 channel activity (conductive signaling) in the pancreatic β-cell is required for normal insulin secretion^27, 28^. Thus, SWELL1-LRRC8 loss-of-function both down-regulates insulin signaling in target tissues^24, 29^ and insulin secretion from the pancreatic β-cell^27, 28^ inducing a state of glucose intolerance^24, 27, 29^. Since Type 2 diabetes (T2D) is characterized by both a loss of insulin sensitivity of target tissues (fat, skeletal muscle, liver) and ultimately, impaired insulin secretion from the pancreatic β-cell^1–3^, these data raised the question: could impaired SWELL1-mediated signaling contribute to T2D pathogenesis, and if so, could this be corrected pharmacologically to improve systemic glycemia?

In this study, we provide evidence that SWELL1-mediated currents and SWELL1 protein are reduced in murine and human adipocytes and pancreatic β-cells in the setting of T2D and hyperglycemia suggesting that dysfunctional SWELL1-mediated signaling could contribute to T2D pathogenesis by impairing insulin sensitivity and insulin secretion. Next, we identify a small molecule modulator, DCPIB (renamed SN-401), as a tool compound that binds the SWELL1-LRRC8 complex^30^, and potentially functions as a chemical chaperone to augment SWELL1 expression and plasma membrane trafficking at concentrations >90% lower than its IC_50_ of ∼5 µM for I_Cl,SWELL_. *In vivo,* SN-401 normalizes glucose tolerance by increasing insulin sensitivity and secretion in insulin-resistant T2D mouse models, while augmenting tissue glucose uptake, suppressing hepatic glucose production, and greatly reducing hepatic steatosis and hepatocyte damage (ballooning) in obese T2D mice. Importantly, while SN-401 normalizes glycemia in diabetic mice, it has very mild glucose-lowering effects on non-obese euglycemic mice – indicating a low risk of hypoglycemic events associated with other commonly used anti-diabetic therapies, including sulfonylureas and insulin. Combining cryo-EM structure data of SN-401 bound to its target SWELL1/LRRC8a^30^ with molecular docking simulations, and novel cryo-EM structure data of an active SN-40X congener bound to SWELL1 hexameric channels in lipid nanodiscs, we validate a structure-activity relationship (SAR) based approach to generate novel SN-401 congeners with subtle molecular changes to either enhance or delete on-target activity, both *in vitro* and *in vivo.* This approach allows us to attribute the cellular and systemic SN-40X effects to drug-target binding, while controlling for off-target effects. We propose small molecule SWELL1 modulators may represent a first-in-class therapeutic approach to treat metabolic syndrome and associated diseases by restoring SWELL1 signaling across multiple organ systems that are dysfunctional in T2D.

## Results

### I_Cl,SWELL_ and SWELL1 protein are reduced in T2D β-cells and adipocytes

SWELL1/LRRC8a ablation impairs insulin signaling in target tissues ^24, 29^ and insulin secretion from the pancreatic β-cell ^27, 28^, inducing a pre-diabetic state of glucose intolerance ^24, 27, 29^. These recent findings suggest that reductions in SWELL1 may contribute to Type 2 diabetes (T2D). To determine if SWELL1-mediated currents are altered in T2D we measured I_Cl,SWELL_ in pancreatic β-cells freshly isolated from T2D mice raised on HFD for 5-7 months (**Fig. 1a&c**) and from T2D patients (**Fig. 1b&d, Supplementary Table S1**) compared to non-T2D controls. In both mouse and human T2D β-cells, the maximum I_Cl,SWELL_ current density (measured at +100 mV) upon stimulation with hypotonic swelling is significantly reduced (90% in murine; 63% in human, **Fig. 1c&d**) compared to non-T2D controls, similar to reductions observed in SWELL1 knock-out (KO) and knock-down (KD) murine and human β-cells^27^, respectively. As SWELL1/LRRC8a is a critical component of I_Cl,SWELL_/VRAC ^21, 22^ in both adipose tissue ^24, 29^ and β-cells ^27, 28^, we asked whether these reductions in I_Cl,SWELL_ in the setting of T2D^32^ are associated with reductions in SWELL1 protein expression. Total SWELL1 protein in diabetic human cadaveric islets (representing numerous islet cell types) also shows a trend toward being reduced 50% compared to islets from non-diabetics (**Fig. 1e, Supplementary Table S2**), suggesting that reduced SWELL1 protein may underlie these reductions in I_Cl,SWELL_ currents.

**Figure 1.**
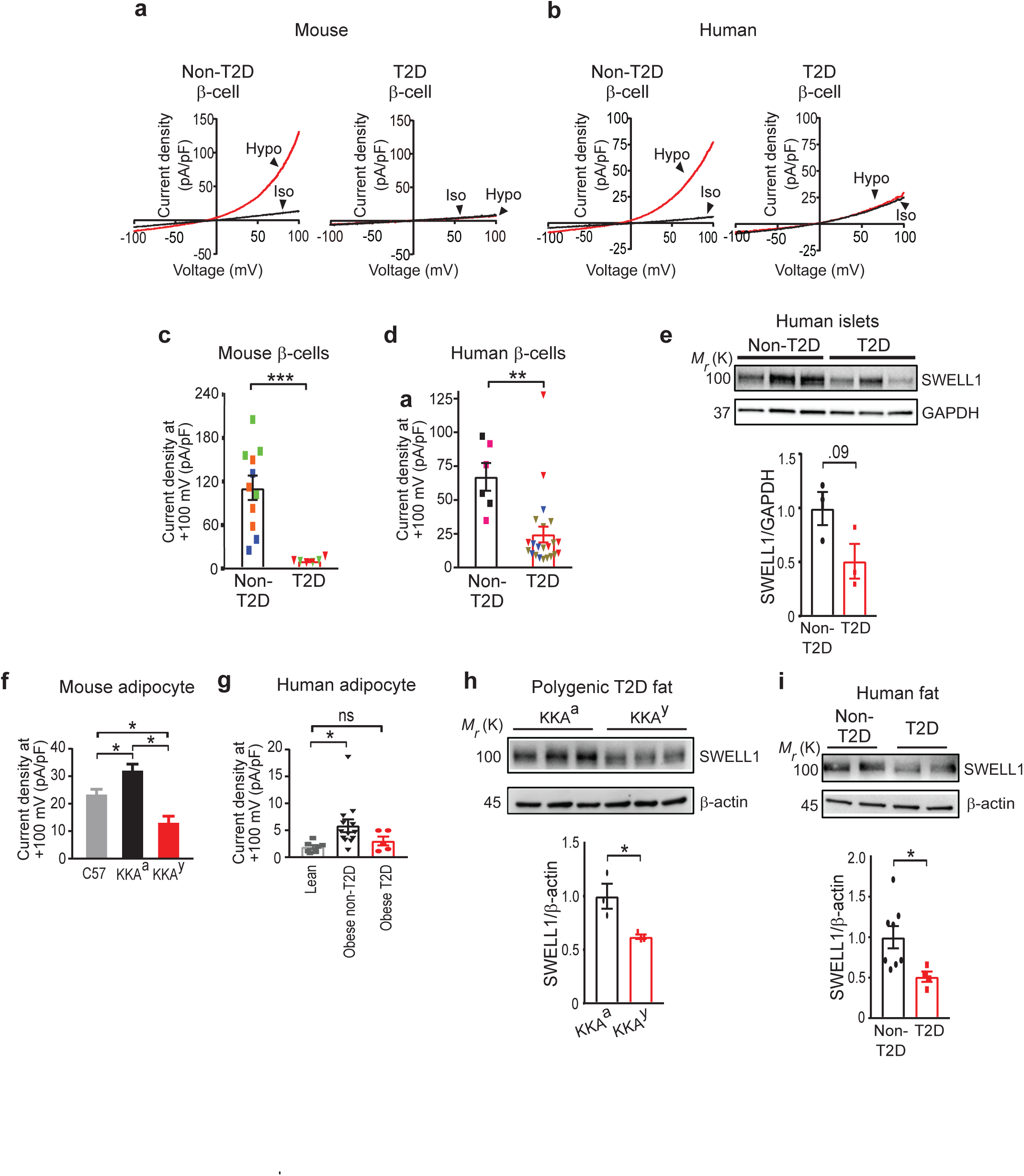
I_Cl,SWELL_ and SWELL1 protein are reduced in T2D β-cells and adipocytes. **a-b.** Current-voltage plots of I_Cl,SWELL_ measured in non-T2D and T2D mouse (**a**) and human (**b**) β-cells at baseline (iso, black trace) and; with hypotonic (210 mOsm) stimulation (hypo, red trace). **c-d.** Mean outward I_Cl,SWELL_ current densities at +100 mV from non-T2D (n = 11 cells) and T2D (n = 6 cells) mouse (**c**) and non-T2D (n = 6 cells) and T2D (n = 22 cells) human (**d**) β-cells. Symbol color represent cells recorded from different individual mice or humans. **e.** Western blot for SWELL1 protein isolated from cadaveric islets of non-T2D and T2D donors (n = 3 each), and densitometry quantification below. **f.** Mean outward I_Cl,SWELL_ current densities at +100 mV from adipocytes isolated from epididymal fat of C57^$^, control strain KKA^a$^ and polygenic-T2D KKA^y$^ mice (n = 14-34 cells). **g.** Mean outward I_Cl,SWELL_ current densities recorded at +100 mV from adipocytes isolated from visceral fat of lean^#^ (n = 7 cells), obese non-T2D^#^ (n = 13 cells) and T2D patients (n = 5 cells). **h.** Western blot for SWELL1 protein expression in epididymal adipose tissue isolated from polygenic-T2D KKA^y^ mice compared to the parental control strain KKA^a^ (n=3 each). **i.** Western blot comparing SWELL1 protein expression in visceral adipose tissue isolated from non-T2D and T2D patients. ^$^Data for C57, KKA ^a^ and KKA^y^ mouse adipocytes are replotted from previously reported data in Inoue, H et al. (2010) and the ^#^lean and obese non-T2D adipocytes are replotted from our previously reported data in Zhang, Y et al. (2017) for purposes of comparison in **f** and **g**. Data are represented as Mean ±SEM. Two-tailed unpaired t-test were used in **c**-**e, h and i**. Two-tailed permutation t-test group comparison for mouse adipocytes and one-way ANOVA for human adipocytes were used in **f** and **g** respectively. *, ** and *** represent *p-values of <0.05, <0.01* and *<0.001,* respectively.

Curiously, reductions in β-cell I_Cl,SWELL_ observed in the setting of T2D (**Fig. 1a-d**) are consistent with previous measurements of VRAC/I_Cl,SWELL_ in the adipocytes of the murine KKA^y^ T2D model^32^, which are reduced by 60% in T2D KKA^y^ mice compared to KKA^a^ controls^32^ (**Fig. 1f**, data plotted from Inoue et al, 2010^32^). Similarly, SWELL1-mediated I_Cl,SWELL_ measured in isolated human adipocytes from an obese T2D patient (BMI =52.3, HgbA1c = 6.9%; Fasting Glucose = 148-151 mg/dl) show a trend toward being reduced 48% compared to obese, non-T2D patients that we reported previously^24^, and not different from I_Cl,SWELL_ in adipocytes from lean patients (**Fig. 1g, Supplementary Table S3**). Consistent with reductions observed in I_Cl,SWELL_, SWELL1 protein is also reduced (38%) in adipose tissue of T2D KKA^y^ mice as compared to parental control KKA^a^ mice (**Fig. 1h**). Similarly, SWELL1 protein is 50% lower in adipose tissue from obese T2D patients (HgbA1c > 6.0%) compared to adipose tissue from normoglycemic obese patients (HgbA1c < 6.0% **Fig. 1i, Supplementary Table S4**). Taken together, these findings suggest reduced SWELL1 activity in adipocytes and β-cells (and possibly other tissues) may underlie insulin-resistance and impaired insulin secretion associated with T2D. Moreover, SWELL1 protein expression increases in both adipose tissue and liver in the setting of early euglycemic obesity^29^ and shRNA-mediated suppression of this SWELL1 induction exacerbates insulin-resistance and glucose intolerance^29^. Therefore, we speculate that maintenance or induction of SWELL1 expression/signaling in peripheral tissues may support insulin sensitivity and secretion to preserve systemic glycemia in the setting of T2D.

### SWELL1 protein expression regulates insulin-stimulated PI3K-AKT2-AS160 signaling

To test whether SWELL1 regulates insulin signaling, we re-expressed Flag-tagged SWELL1 (SWELL1 O/E) in SWELL1 KO 3T3-F442A adipocytes and measured insulin-stimulated phosphorylated AKT2 (pAKT2) and phosphorylated AS160 (pAS160) as a readout of insulin-signaling (**Fig. 2a&b**). SWELL1 KO 3T3-F442A adipocytes exhibit significantly blunted insulin-mediated pAKT2 and pAS160 signaling compared to WT adipocytes, similar to described previously^24, 26^, and this is fully rescued by re-expression of SWELL1 in SWELL1 KO adipocytes (KO+SWELL1 O/E, **Fig. 2a&b**)^26^. SWELL1 re-expression also recapitulates SWELL1-mediated I_Cl,SWELL_ in in SWELL1 KO cells in response to hypotonic stimulation (**Fig. 2c and Supplementary Fig. S1a-c**), which is consistent with restoration of SWELL1-LRRC8a signaling complexes at the plasma membrane. Notably, the reductions in total AKT2 protein expression observed in SWELL1 KO adipocytes is not rescued by SWELL1 re-expression, indicating that transient changes in SWELL1 protein expression in adipocytes regulates pAKT2 signaling, as opposed to total AKT2 protein expression. We confirmed FLAG-tagged SWELL1 traffics normally to the plasma membrane when expressed in both WT and SWELL1 KO adipocytes visualized by immunofluorescence (IF) using anti-FLAG and SWELL1 KO-validated anti-SWELL1 antibodies, respectively (**Supplementary Fig. S1d&e**). FLAG-tagged SWELL1 overexpressed in WT and SWELL1 KO adipocytes assumes a punctate pattern at the cell periphery, similar to endogenous SWELL1 in WT adipocytes. Overall, these data indicate that SWELL1 expression levels regulate insulin-PI3K-AKT2-AS160 signaling in adipocytes - potentially by modulating GRB2 signaling^20, 24–26^. Furthermore, these data imply that pharmacological SWELL1 induction in peripheral tissues in the setting of T2D may enhance insulin signaling and improve systemic insulin-sensitivity and glycemic control.

**Figure 2.**
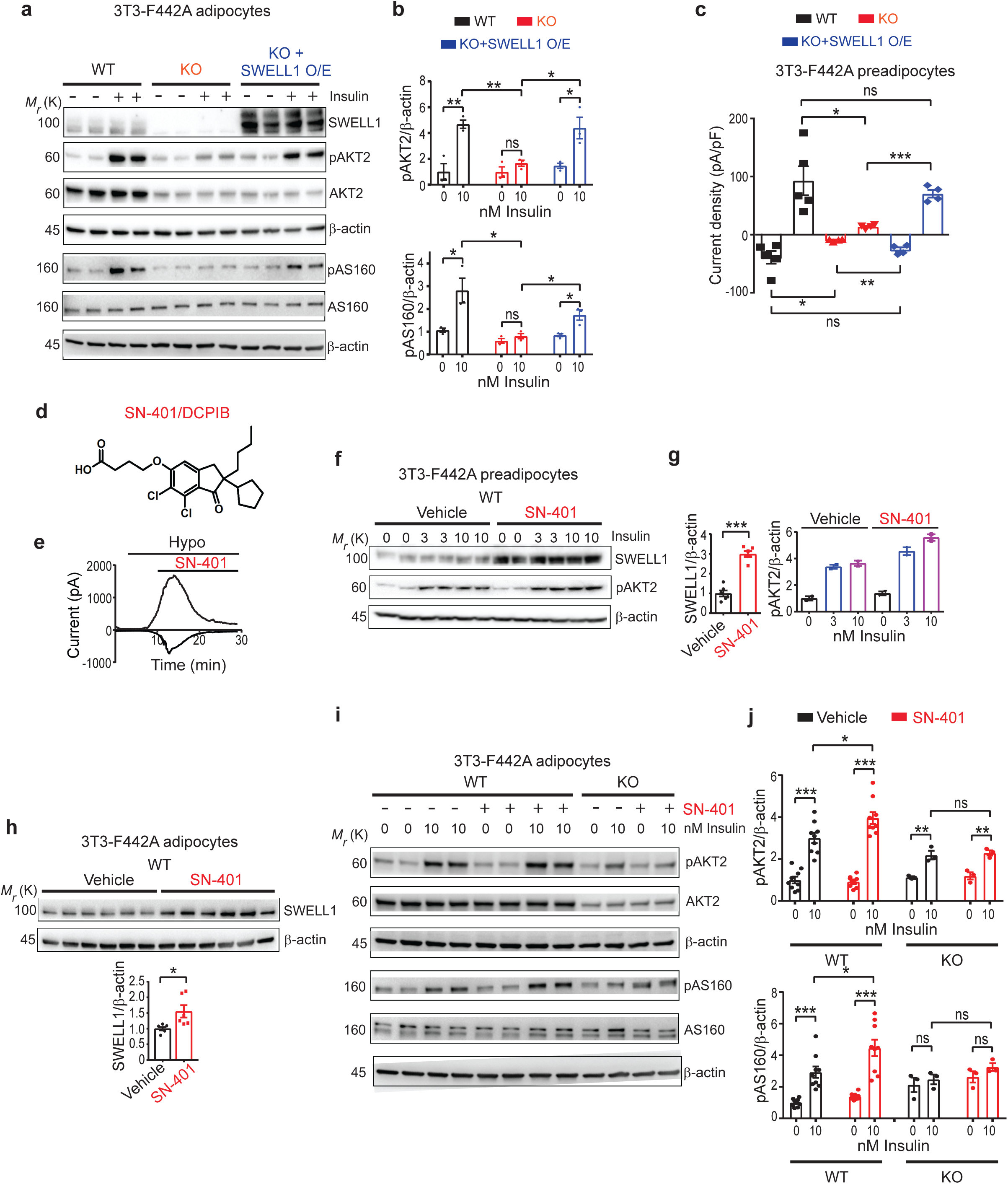
SWELL1 protein expression regulates insulin-stimulated PI3K-AKT2-AS160 signaling. **a-b.** Western blots detecting SWELL1, pAKT2, AKT2, pAS160, AS160 and β-actin in wildtype (WT, black), SWELL1 knockout (KO, red) and adenoviral re-expression of SWELL1 in KO (KO+SWELL1 O/E, blue) 3T3-F442A adipocytes stimulated with 0 and 10 nM insulin for 15 min (**a**) and the corresponding densitometric ratio for pAKT2/β-actin and pAS160/β-actin are shown on the right (**b**) (n = 3 independent experiments for each condition). **c.** Mean inward and outward current densities recorded at -100 and +100 mV from WT (black, n = 5 cells), KO (red, n = 4 cells) and KO+SWELL1 O/E (blue, n = 4 cells) 3T3-F442A preadipocytes. **d.** SN-401/DCPIB chemical structure. **e.** I_Cl,SWELL_ inward and outward current over time upon hypotonic (210 mOsm) stimulation and subsequent inhibition by 10 μM SN-401 in a HEK-293 cell. **f-g.** Western blots detecting SWELL1, pAKT2 and β-actin in WT 3T3-F442A preadipocytes treated with either vehicle or SN-401 (10 µM) for 96 hours (**f**) (n=2 each) and the corresponding densitometric ratio (**g**). **h.** Western blots detecting SWELL1 and β-actin in WT 3T3-F442A adipocytes treated with either vehicle or SN-401 (10 µM) for 96 hours (n=6 each) and the corresponding densitometric ratio below. **i-j.** Western blots detecting pAKT2, AKT2, pAS160, AS160 and β-actin in WT (n=9 each) and SWELL1 KO (n=3 each) 3T3-F442A adipocytes treated with either vehicle or SN-401 (500 nM) for 96 hours and stimulated with 0 and 10 nM insulin for 15 min (**i**) and the corresponding densitometric ratio for pAKT2/β-actin and pAS160/ β-actin respectively (**j**). Data are represented as Mean ±SEM. Two-tailed unpaired t-test was used in **b, c, g** and **j** where *, ** and *** represent *p-values of <0.05*, *<0.01* and *<0.001*, respectively. ns, difference did not exceed the threshold for significance.

### A small molecule binds SWELL1-LRRC8 channel complexes, increases adipocyte SWELL1 protein expression and SWELL1-dependent insulin signaling

The small molecule 4-[(2-Butyl-6,7-dichloro-2-cyclopentyl-2,3-dihydro-1-oxo-1*H*-inden-5-yl)oxy]butanoic acid *(DCPIB***, Fig. 2d**) is among a series of structurally diverse (acylaryloxy)acetic acid derivatives, that were synthesized and studied for diuretic properties in the late 1970s^33, 34^ and evaluated in the 1980s as potential treatments for brain edema ^35, 36^. While DCPIB was derived from the FDA-approved diuretic ethacrynic acid, it has minimal diuretic activity^37^, and has instead been used as a selective VRAC/I_Cl,SWELL_ inhibitor^24, 27, 31^ (**Fig. 2e**), binding at a constriction point within the SWELL1-LRRC8 hexamer^30, 38–40^. Having demonstrated that SWELL1 is regulates insulin-AKT2 signaling in multiple cell types, including adipocytes^24, 25, 29^, skeletal muscle^26^, and endothelium^41^, we anticipated pharmacological inhibition of VRAC/I_Cl,SWELL_ with DCPIB, which we here re-name SN-401, would decrease insulin signaling. Unexpectedly, SN-401 increased SWELL1 protein expression in 3T3-F442A preadipocytes (3-fold control expression; **Fig. 2f&g**) and adipocytes (1.5-fold control expression; **Fig. 2h**) when applied for 96 hours, and was associated with enhanced insulin-stimulated levels of pAKT2 (**Fig. 2f-g&i-j**), and insulin-stimulated levels of pAS160 (**Fig. 2i&j**). These SN-401-mediated effects on insulin-AKT2-AS160 signaling are absent in SWELL1 KO 3T3-F442A adipocytes, consistent with an on-target SWELL1-mediated mechanism of action for SN-401 (**Fig. 2i&j**). The SN-401-mediated increases in SWELL1 protein expression are not associated with increases in SWELL1 mRNA, nor in the mRNA for other LRRC8 subunits: LRRC8b, LRRC8c, LRRC8d or LRRC8e that form the SWELL1 channel complex (**Supplementary Fig. S2a-c**), implicating post-transcriptional mechanisms for increased SWELL1 expression and SWELL1-LRRC8 associated signaling.

### SN-401 increases SWELL1 and improves systemic glucose homeostasis in murine T2D models by enhancing insulin sensitivity and secretion

To determine if SN-401 improves insulin signaling and glucose homeostasis *in vivo* we treated two T2D mouse models: obese, HFD-fed mice and the polygenic T2D KKA^y^ mouse model with SN-401 (5 mg/kg i.p. for 4-10 days). *In vivo,* SN-401 augments SWELL1 expression 2.3-fold in adipose tissue of HFD-fed T2D mice (**Fig. 3a**). Similarly, SN-401 increases SWELL1 expression in adipose tissue of T2D KKA^y^ mice to levels comparable to both non-T2D C57/B6 mice and to the parental KKA^a^ parental strain (**Fig. 3b**). This restoration of SWELL1 expression is associated with normalized fasting blood glucose (FG), glucose tolerance (GTT), and markedly improved insulin-tolerance (ITT) in both HFD-induced T2D mice (**Fig. 3c**) and in the polygenic T2D KKAy model (**Fig. 3d-f)**, without significant reductions in body weight (**Supplementary Table S5**). Remarkably, treating the control KKA^a^ parental strain with SN-401 at the same treatment dose (5 mg/kg x 4-10 days) does not cause hypoglycemia, nor does it alter glucose and insulin tolerance (**Fig. 3d-f**). Similarly, lean, non-T2D, glucose-tolerant mice treated with SN-401 have similar FG, GTT and ITT compared to vehicle-treated mice (**Fig. 3g&h and Supplementary Fig. S3a-c**). However, when made insulin-resistant and diabetic after 16 weeks of HFD feeding, these same mice (from **Fig. 3g&h**) treated with SN-401 show marked improvements in FG (**Fig. 3i**), GTT and ITT (**Fig. 3j**) as compared to vehicle. These data show that SN-401 restores glucose homeostasis in the setting of T2D, but has little effect on glucose homeostasis in non-T2D mice. Importantly, this portends a low risk for inducing hypoglycemia. SN-401 was well-tolerated during chronic i.p. injection protocols, with no overt signs of toxicity with daily i.p. injections for up to 8 weeks, despite striking effects on glucose tolerance (**Supplementary Fig. S3d**).

**Figure 3.**
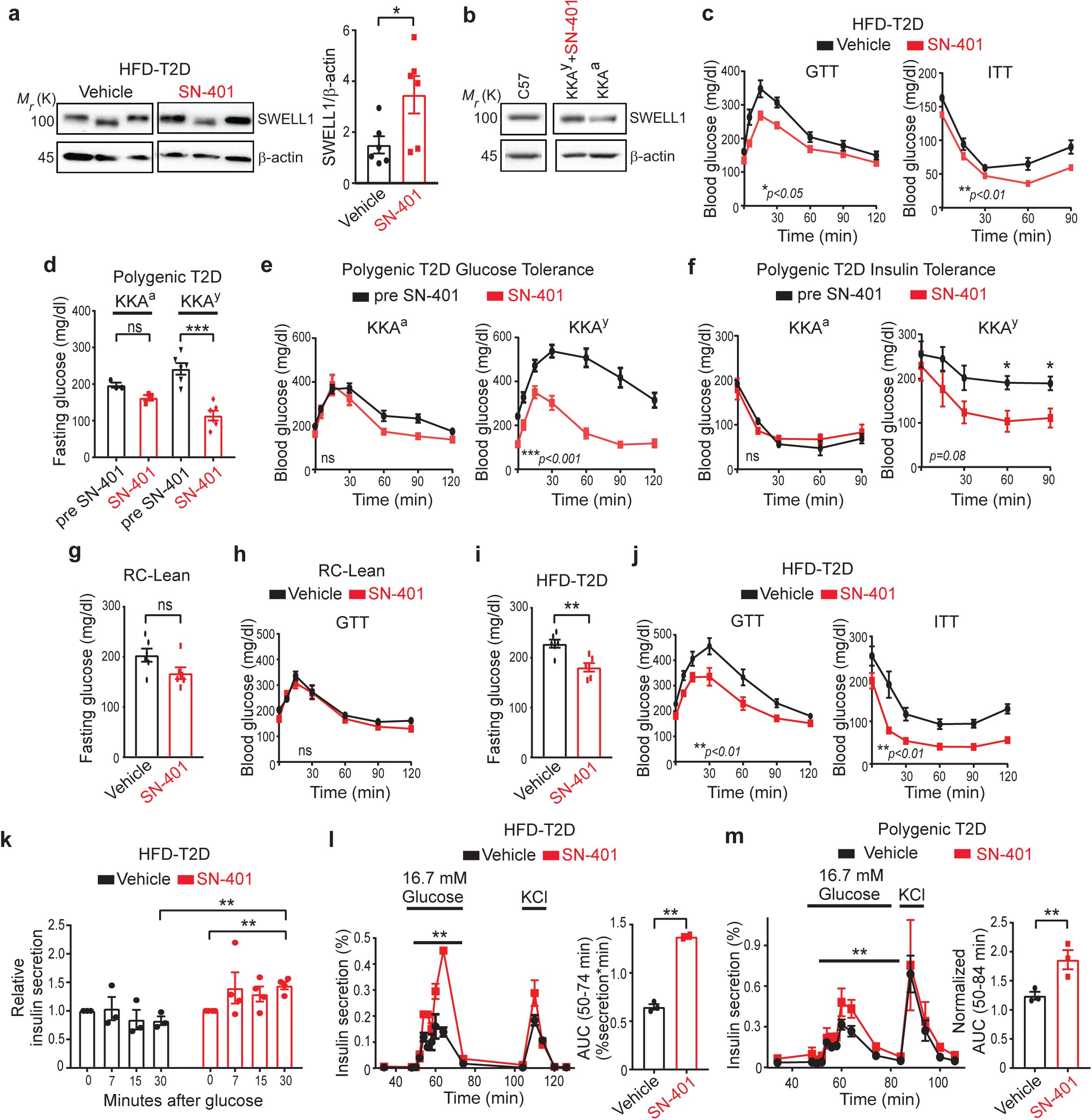
SN-401 increases SWELL1 and improves glycemic control in murine T2D models by enhancing insulin sensitivity and secretion. **a.** Western blots detecting SWELL1 protein in visceral fat of C57BL/6 mice on high-fat diet (HFD) for 21 weeks and treated with either vehicle or SN-401 (5 mg/kg i.p., as in scheme Fig. 4e) and the corresponding densitometric ratios for SWELL1/β-actin (right) (n = 6 mice in each group). **b.** Western blots comparing SWELL1 protein expression in inguinal adipose tissue of a polygenic-T2D KKA^y^ mouse treated with SN-401 (5 mg/kg i.p daily x 14 days) compared to untreated control KKA^a^ and wild-type C57BL/6 mice. **c.** Glucose tolerance test (GTT) and insulin tolerance test (ITT) of C57BL/6 mice on HFD for 8 weeks treated with either vehicle or SN-401 (5 mg/kg i.p) for 10 days (n = 7 mice in each group). **d-f.** Fasting glucose levels **(d)**, GTT **(e)** and ITT **(f)** of T2D KKA^y^ mice (n = 6) and its control strain KKA^a^ (n = 3) compared pre- and post-SN-401 (5 mg/kg i.p) treatment for 4 days, respectively. **g-h.** Fasting glucose levels **(g)** of regular chow-diet fed (RC), lean mice treated with either vehicle or SN-401 (5 mg/kg i.p) for 6 days and the corresponding GTT **(h)** (n = 6 in each group). **i-j.** Fasting glucose levels **(i)** and GTT (16 weeks HFD, 4 days treatment) and ITT (18 weeks HFD, 4 days treatment) **(j)** of HFD-T2D mice treated with either vehicle or SN-401 (5 mg/kg i.p). **k.** Relative insulin secretion in plasma of HFD-T2D mice (18 weeks HFD, 4 days treatment) after i.p. glucose (0.75 g/kg BW) treated with either vehicle (n = 3) or SN-401 (n = 4; 5 mg/kg i.p). **l-m.** Glucose stimulated insulin secretion (GSIS) perifusion assay of islets isolated from HFD-T2D mouse (21 week time point) treated with either vehicle (n = 3 mice, and 3 experimental replicates) or SN-401 (n = 3 mice, and 2 experimental replicates, 5 mg/kg i.p) **(l)** and from polygenic-T2D KKA^y^ mouse treated with either vehicle or SN-401 (5 mg/kg i.p for 6 days, n = 3 mice in each group, 3 experimental replicates), **(m)**; and their corresponding area under the curve (AUC) comparisons, respectively, shown on the right. Mean presented ±SEM. Two-tailed unpaired t-test was used in in **a**,**g**, **i**, **l** and **m**. Paired t-test was used in **d**. Paired (in group) and unpaired (between group) t-tests were performed in **k**. Two-way ANOVA was used for **c**, **e**, **f**, **h** and **j** (p-value in bottom corner of graph). *, ** and *** represent *p-values of <0.05*, *<0.01* and *<0.001,* respectively. ns, difference did not exceed the threshold for significance.

To examine the possible contribution of SN-401-mediated enhancements in insulin secretion from pancreatic β-cells, we next measured glucose-stimulated insulin secretion (GSIS) in SN-401 treated mice subjected to 21 weeks of HFD. We found that the impairments in GSIS commonly observed with long-term HFD (21 weeks HFD) are significantly improved in SN-401-treated HFD mice based on serum insulin measurements (**Fig. 3k**) and perifusion GSIS from isolated islets (**Fig. 3l**), consistent with the predicted effect of SWELL1 induction in pancreatic β-cells ^27, 28^. Similar results are obtained in perfusion assays performed in SN-401 compared to vehicle treated T2D KKA^y^ mice (**Fig. 3m**). Collectively, these data suggest that SN-401-mediated improvements in systemic glycemia in T2D occur via augmentation of both peripheral insulin sensitivity and β-cell insulin secretion – the inverse phenotype to *in vivo* loss-of-function studies^24,27–29^.

### SN-401 improves systemic insulin sensitivity, tissue glucose uptake, and nonalcoholic fatty liver disease in murine T2D models

To more rigorously evaluate SN-401 effects on insulin sensitization and glucose metabolism in T2D mice we performed euglycemic hyperinsulinemic clamps traced with ^3^H-glucose and ^14^C-deoxyglucose in T2D KKA^y^ mice treated with SN-401 or vehicle. SN-401 treated T2D KKA^y^ mice require a higher glucose-infusion rate (GIR) to maintain euglycemia compared to vehicle, consistent with enhanced systemic insulin-sensitivity (**Fig. 4a**). The rate of glucose appearance (R_a_), which reflects hepatic glucose production from gluconeogenesis and/or glycogenolysis, was reduced 40% in SN-401-treated T2D KKA^y^ mice at baseline (Basal, **Fig. 4b**), and further suppressed 75% during glucose/insulin infusion (Clamp, **Fig. 4b**), revealing SN-401 increases hepatic insulin sensitivity – similar to thiazolidinediones (TZD)^42^.

**Figure. 4.**
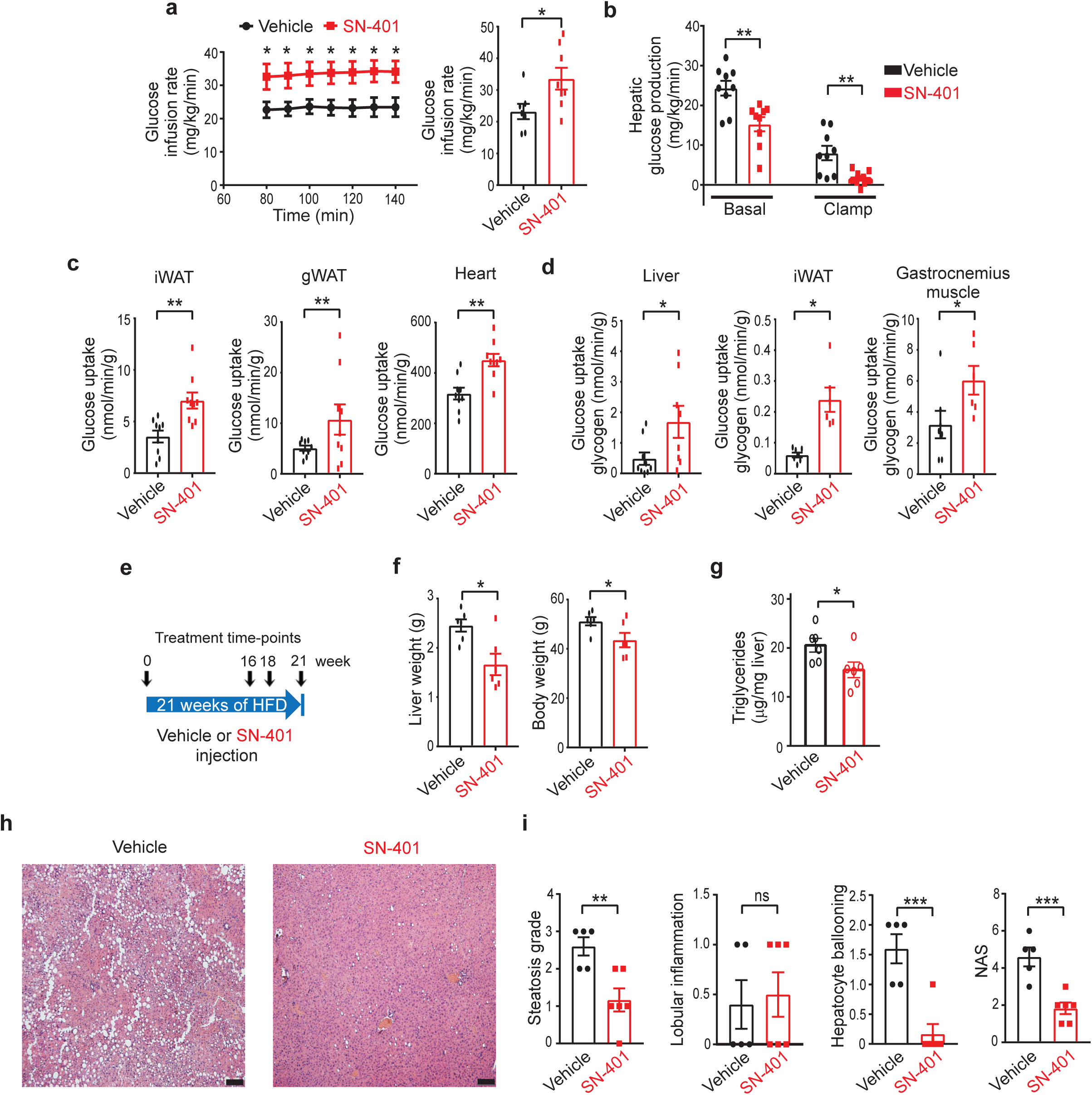
SN-401 improves systemic insulin sensitivity, tissue glucose uptake and nonalcoholic fatty liver disease in murine T2D models. **a.** Mean glucose-infusion rate during euglycemic hyperinsulinemic clamps of polygenic T2D KKA^y^ mice treated with vehicle (n = 7) or SN-401 (n = 8) for 4 days (5 mg/kg i.p). **b.** Hepatic glucose production at baseline and during euglycemic hyperinsulinemic clamp of T2D KKA^y^ mice treated with vehicle or SN-401 (n = 9 in each group) for 4 days (5 mg/kg i.p). **c.** Glucose uptake determined from 2-deoxyglucose (2-DG) uptake in inguinal while adipose tissue (iWAT) and gonadal white adipose trissue (gWAT) and heart during traced clamp of T2D KKA^y^ mice treated with vehicle or SN-401 (n = 9 in each group) for 4 days (5 mg/kg i.p). **d.** Glucose uptake into glycogen determined from 2-DG uptake in liver (n = 9 for vehicle and n = 8 for SN-401), adipose (iWAT, n = 7 vehicle and n = 6 SN-401) and gastrocnemius muscle (n = 7 vehicle and n = 6 SN-401) during clamp of T2D KKA^y^ mice treated with either Vehicle or SN-401 for 4 days (5 mg/kg i.p). **e.** Schematic representation of treatment protocol of C57BL/6 mice injected with either vehicle or SN-401 (n = 6 in each group) during HFD-feeding. **f.** Liver mass (left) and body weight (right) of HFD-T2D mice following treatment with either vehicle or SN-401 (5 mg/kg i.p.) in **e**. **g-i**. Liver triglycerides (**g**; 6 mice in each group), corresponding representative hematoxylin- and eosin-stained liver sections **(h)**, histologic scoring for steatosis, lobular inflammation, hepatocyte damage (ballooning) and NAFLD-activity score (NAS; vehicle=5 and SN-401=6) with integrated scores for steatosis, inflammation, and ballooning **(i**) of liver samples from **e**. Mean presented ±SEM. Two-tailed unpaired t-test in **a**-**d**, **f**, **g** and **i**. Statistical significance is denoted by *, ** and *** representing *p<0.05*, *p<0.01* and *p<0.001,* respectively. Scale bar - 100 µm.

As the SN-401-mediated increase in SWELL1 is expected to enhance insulin-pAKT2-pAS160 signaling, GLUT4 plasma membrane translocation, and tissue glucose uptake ^24^, we next measured the effect of SN-401 on glucose uptake in adipose, myocardium and skeletal muscle using 2-deoxyglucose (2-DG). SN-401 enhanced insulin-stimulated 2-DG uptake into inguinal white adipose tissue (iWAT), gonadal white adipose tissue (gWAT), and myocardium (**Fig. 4c**). As SWELL1 ablation markedly reduces insulin-pAKT2-pGSK3β signaling^24, 26, 41^ and cellular glycogen content^24^, we asked whether the SN-401-mediated increase in SWELL1 would increase glucose incorporation into tissue glycogen in the setting of T2D. Indeed, liver, adipose, and skeletal muscle glucose incorporation into glycogen is markedly increased in SN-401-treated mice (**Fig. 4d**), consistent with a SWELL1-mediated insulin-pAKT2-pGSK3β-glycogen synthase gain-of-function.

Nonalcoholic fatty liver disease (NAFLD), like T2D, is associated with insulin resistance ^43^. NASH is an advanced form of nonalcoholic liver disease defined by three histological features: hepatic steatosis, hepatic lobular inflammation, hepatocyte damage (ballooning) and can be present with or without fibrosis. NAFLD and T2D likely share at least some pathophysiologic mechanisms because more than one-third of patients (37%) with T2D have NASH ^44^ and almost one-half of patients with NASH (44%) have T2D ^45^. To evaluate the effect of SN-401 on the genesis of NAFLD, mice were raised on HFD for 16 weeks followed by intermittent dosing with SN-401 over the course of 5 weeks (**Fig. 4e**). These mice had grossly smaller livers (**Fig. 4f**), lower hepatic triglyceride concentrations (**Fig. 4g)** and experienced a mild 14% reduction in body weight compared to vehicle-treated mice (**Fig. 4f).** Histologic evaluation revealed significant reductions in hepatic steatosis and hepatocyte damage compared to vehicle-treated mice (**Fig. 4h&i, Supplemental Fig. S4**). The NAFLD activity score (NAS), which integrates histologic scoring of hepatic steatosis, lobular inflammation, and hepatocyte ballooning^46^ (**Fig. 4i**), also improved >2 points in SN-401-treated mice compared to vehicle-treated mice. These SN-401 mediated reductions in hepatic steatosis and hepatocyte damage are consistent with the observed increases in hepatic insulin sensitivity and consequent reductions in hepatic glucose production via gluconeogenesis available for hepatic *de novo* lipogenesis, as observed with other insulin sensitizers, such as metformin and TZDs^47^. Taken together, these data reveal that SN-401 augments SWELL1 protein and SWELL1-mediated signaling to concomitantly enhance both systemic insulin sensitivity and pancreatic β-cell insulin secretion, thereby normalizing glycemic control in T2D mouse models. This improved metabolic state can reduce ectopic lipid deposition, hepatocyte damage, and NAFLD that is associated with obesity and T2D.

### Chemical synthesis, molecular docking and cryo-EM reveal specific SN-401-SWELL1 interactions required for on-target activity

To confirm that SN-401-induced increases in SWELL1 protein and signaling are mediated by direct binding to the SWELL1-LRRC8 channel complex, as opposed to off-target effects, we designed and synthesized novel SN-401 congeners (**Fig. 5a**) with subtle structural changes that either enhanced (SN-403, SN-406, SN-407), or entirely eliminated (Inactive1, Inactive2) SN-401 inhibition of I_Cl,SWELL_ (**Fig. 5b&c; Supplementary Fig. S5a-c**). The cryo-EM structure of SN-401/DCPIB bound within the SWELL1 homo-hexamer revealed that SN-401 binds at a constriction point in the pore wherein the electronegative SN-401 carboxylate group interacts electrostatically with the R103 residue in one or more of the SWELL1 monomers^30^. Moreover, SN-401 appeared to stabilize the pore region of the SWELL1 hexamer in lipid-nanodiscs^30^. To characterize the structural features of SN-401 responsible for binding to SWELL1-LRRC8, we performed molecular docking simulations of SN-401 and its analogs into the SWELL1 homo-hexamer (PDB: 6NZZ), and identified two molecular determinants predicted to be critical for SN-401-SWELL1-LRRC8 binding (**Fig. 5d**): (1) the length of the carbon chain leading to the anionic carboxylate group predicted to electrostatically interact with one or more R103 guanidine groups (found in SWELL1/LRRC8a and LRRC8b; **Fig. 5d-solid circles**); and (2) proper orientation of the hydrophobic cyclopentyl group that slides into a hydrophobic cleft at the interface of LRRC8 monomers (conserved among all LRRC8 subunit interfaces; **Fig. 5d-broken circles**). Docking simulations predicted shortening the carbon chain leading to the carboxylate by 2 carbons would yield a molecule, Inactive 1, that could *either* interact with R103 through the carboxylate group (**Supplementary Fig. S5d*(i)***), *or* have the cyclopentyl ring occupy the hydrophobic cleft (**Supplementary Fig. S5d*(ii)***), but unable to simultaneously participate in both interactions (**Supplementary Fig. S5d**). Similarly, docking simulations predicted removing the butyl group of SN-401 would yield a molecule, Inactive 2, unable to orient the cyclopentyl group into a position favorable for interaction with the hydrophobic cleft without introducing structural strain in the molecule (**Supplementary Fig. S5e**, **black arrow**). Both of these structural modifications, predicted to abrogate either carboxylate-R103 electrostatic binding (Inactive 1) *or* cyclopentyl-hydrophobic pocket binding (Inactive 2) were sufficient to eliminate I_Cl,SWELL_ inhibitory activity *in vitro* (**Fig. 5b&c**). Conversely, lengthening the carbon chain attached to the carboxylate group of SN-401 by 1-3 carbons (SN-403; 1 carbon, SN-406: 2 carbons, and SN407: 3 carbons) was predicted to enhance the R103 electrostatic interactions (**Fig. 5e; Supplementary Fig. S5f&g, black solid circle**), and better orient the cyclopentyl group to bind within the hydrophobic cleft (**Fig. 5e, Supplementary Fig. S5f&g, broken circle**). Additional binding interactions for SN-403, SN-406 and SN-407 are also predicted along the channel, due to the longer carbon chains affording additional hydrophobic interactions with side chain carbons of the R103 residues (**Fig. 5e; Supplementary Fig. S5f&g, purple dashes).** As anticipated, SN-403, SN-406, and SN-407 had increased I_Cl,SWELL_ inhibitory activity as compared to SN-401 (**Fig. 5b&c; Supplementary Fig. S5a-c**).

**Figure 5.**
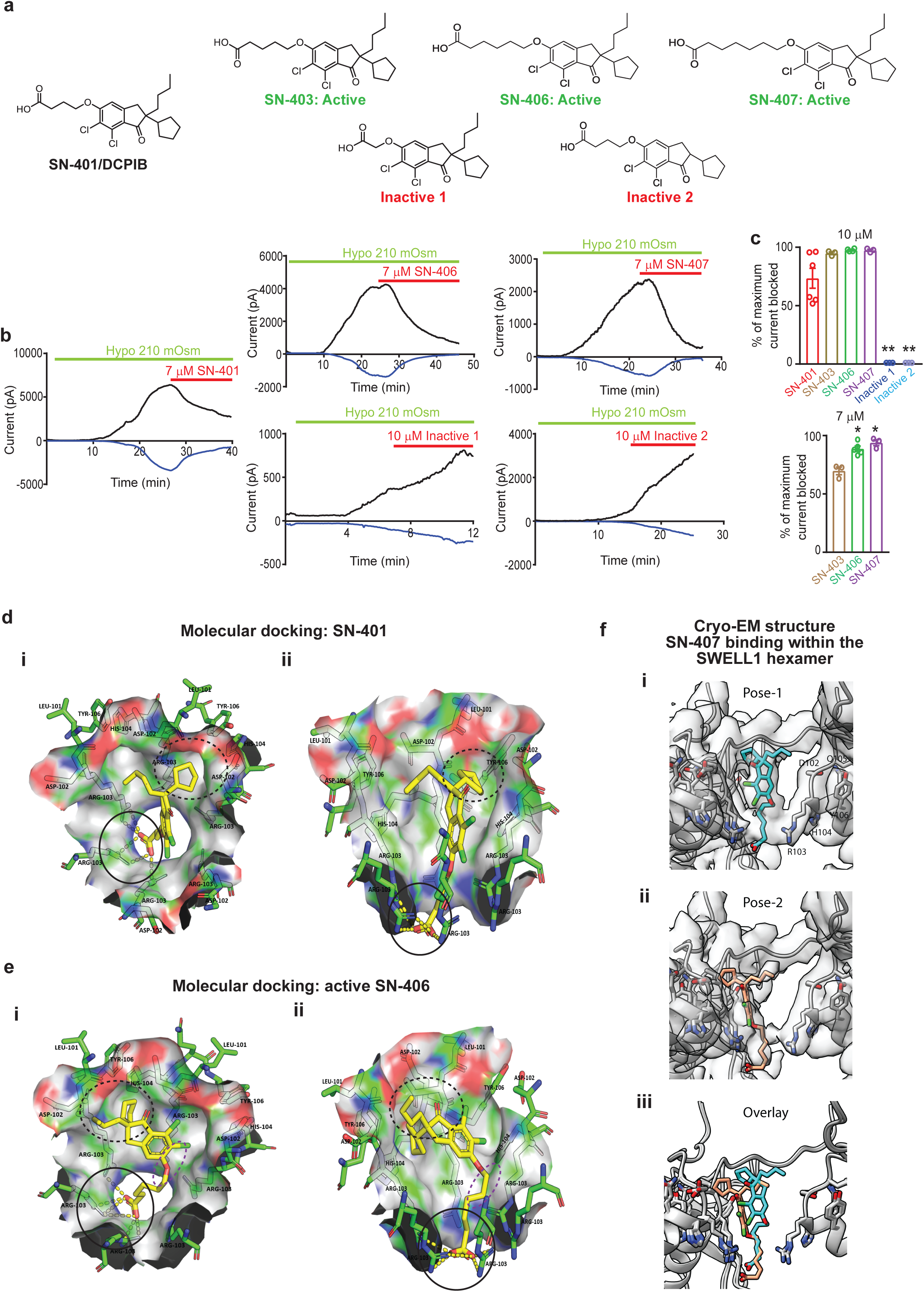
Structure Activity Relationship, molecular docking, and cryo-EM reveal specific SN-401-SWELL1 interactions required for on-target activity. **a.** Chemical structures of SN-401, SN-403, SN-406, SN-407, Inactive 1 and Inactive 2. **b.** I_Cl,SWELL_ inward and outward current over time upon hypotonic (210 mOsm) stimulation and subsequent inhibition with 7 μM SN-401/SN-406 or 10 μM Inactive 1&2 in HEK-293 cells. **c.** Mean of percentage of maximum outward current blocked by SN-401 (n=6), SN-403 (n=3), SN-406 (n=4), Inactive 1 (n=3) and Inactive 2 (n=3) at 10 µM (top) and by SN-403 (n=3), SN-406 (n=5) and SN-407 (n=3) at 7 µM (bottom) in HEK-293 cells, respectively. **d.** top (i) and side (ii) view of binding poses of SN-401; SN-401 carboxylate groups interacts with R103 residue guanidine groups (solid black circle), the SN-401 cyclopentyl group occupies a shallow hydrophobic cleft at the interface of two monomers formed by SWELL1 D102 and L101 (black broken circle)^#^. **e.** top (i) and side view (ii) of best binding pose of SN-406; the carboxylate group interacts with R103 (black circle), cyclopentyl group occupies the hydrophobic cleft (black broken circle) and the alkyl side chain of SN-406 interacts with the alkyl side chain of R103 (purple broken circle) ^#^. **f.** Cryo-EM images revealing: Pose-1 (i), Pose-2 (ii) and overlay of Pose 1&2 (iii) views of selectivity filter with SN-407 bound from the membrane plane (side view). The atomic model is represented as ribbons and sticks within the cryo-EM density with three subunits removed in the side view for clarity. Cryo-EM density is represented in transparent gray, nitrogens are colored blue, oxygens red, chlorines green, protein carbons gray, and SN-407 carbons teal (Pose1) or orange (Pose-2). ^#^Poses generated for respective compounds by docking into PDB 6NZZ using Molecular Operating Environment 2016 (MOE) software package. SN-401 and SN-406 are depicted as yellow sticks and R103, D102 and L101 are depicted as green sticks in **d** and **e**. Mean presented ±SEM. Two-tailed unpaired t-test was used in **c.** *, **, and *** represent *p-values of <0.05*, *<0.01* and *<0.001,* respectively.

To test the predictions of our molecular docking simulations, we used cryo-EM to determine the structure of the SWELL1 homohexamer in lipid nanodiscs in the presence of SN-407 (**Fig. 5f**, **Supplementary Figs. S6-8**). In initial maps using six-fold symmetry, SN-407 density was less apparent than for DCPIB/SN-401^30^, potentially due to a reduction in SN-407 occupancy that is a consequence of lower compound solubility or the presence of multiple drug poses in different particles (**Supplementary Fig. S7**). Therefore, we utilized six-fold symmetry expansion and symmetry relaxation^48^ and were able to resolve two distinct poses for SN-407 with similar occupancy. Pose-1 shows the drug oriented vertically in the channel’s selectivity filter in a manner that is similar to that observed for DCPIB/SN-401, but with the lengthened carboxylate chain coiling to maintain its interaction with R103 (**Fig. 5f(i)**). In Pose-2, SN-407 is tilted off the SWELL1 central axis, positioning its cyclopentyl group closer to the hydrophobic cleft between SWELL1 subunits (**Fig. 5f(ii)**). These data confirm lengthening the carboxylate chain in SN-407 preserves electrostatic interactions with R103 and enables additional contacts between the carboxylate chain and upper hydrophobic moieties in distinct binding poses, consistent with the results of our docking simulations. Collectively, these molecular docking, cryo-EM and functional experiments indicate that SN-401 and SWELL1-active congeners (SN-403/406/407) bind to SWELL1-LRRC8 hexamers at both R103 (via carboxylate end) and at the interface between adjacent LRRC8 monomers (via hydrophobic end), to potentially stabilize the closed state of the channel, and thereby inhibit I_Cl,SWELL_ activity. Indeed, SN-401/SN-407 cryo-EM data, and molecular docking simulations likely reflects the closed state of the SWELL1 homomer. This raises the possibility that SN-401 and active SN-40X compounds bind with higher affinity to SWELL1-LRRC8 complexes in the closed state most commonly encountered under native, physiological conditions, as compared to the open state. Moreover, we hypothesize these SN-40X compounds may function as molecular tethers to stabilize assembly of the SWELL1-LRRC8 hexamer by binding between R103 and adjacent LRRC8 monomers, potentially reducing SWELL1-LRRC8 complex disassembly, subsequent proteasomal degradation, and thereby augment translocation from ER to plasma membrane signaling domains, akin to pharmacological chaperones^49,50,51,52,53^

### SN-401 and SWELL1-active congener SN-406 inhibit I_Cl,SWELL_ and promote SWELL1 dependent signaling at sub-micromolar concentrations

To test this hypothesis, we compared the potency of SN-401/SN-406 to block I_Cl,SWELL_ when applied to closed as compared to open SWELL1-LRRC8 channels. SN-401/SN-406 concentrations of 7-10 μM are required to effectively block channels when first opened by hypotonic activation (**Fig. 5b&c**)^31^. In contrast, application of 1 μM of SN-401 or SN-406 to SWELL1-LRRC8 channels in the closed state (i.e. for 30 min *prior* to hypotonic activation) markedly suppressed and delayed subsequent hypotonic activation of I_Cl,SWELL_, compared to either vehicle alone, or Inactive compounds (**Fig. 6a&b**). SN-401 and SN-406 at concentrations as low as 250 nM had a similar effect (**Fig. 6c&d**). These data support the notion that SN-40X compounds bind with higher affinity to SWELL1-LRRC8 channels in the closed/resting state than the open/activated state, and stabilize the closed conformation at less than one-tenth the concentration required to inhibit activated SWELL1-LRRC8 channels.

**Figure 6.**
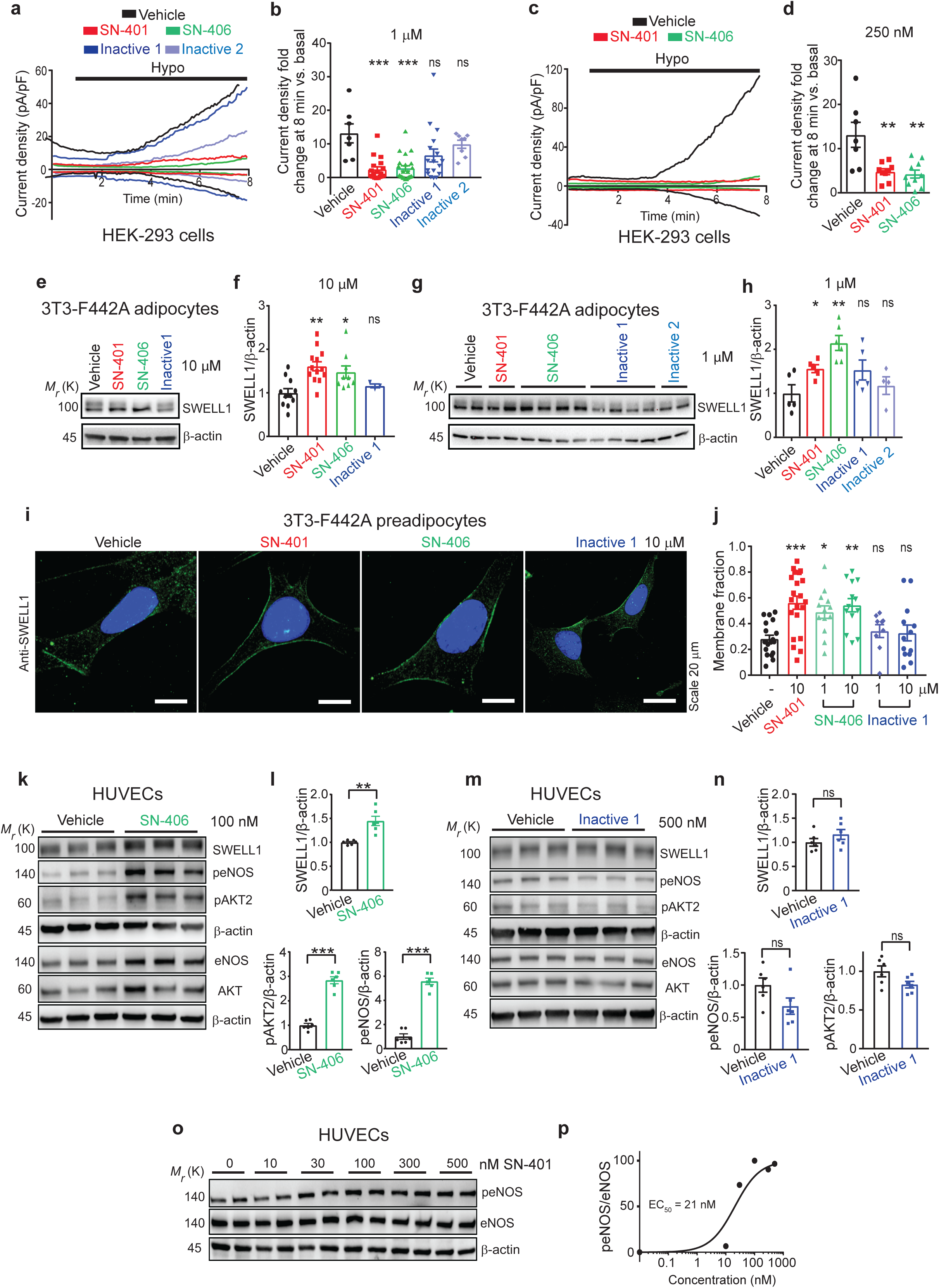
SN-401 and SWELL1-active congener SN-406 inhibit I_Cl,SWELL_ and promote SWELL1 dependent signaling at sub-micromolar concentrations. **a-d.** Representative I_Cl,SWELL_ inward and outward current traces over time recorded from HEK-293 cells preincubated for 30 min with vehicle, SN-401, SN-406, Inactive 1 or Inactive 2 at concentrations of 1 µM **(a)** and vehicle, SN-401 and SN-406 at concentrations of 250 nM **(c),** and subsequently stimulated with hypotonic solution in the presence of the compound. Corresponding fold change in mean outward I_Cl,SWELL_ current densities at +100 mV measured at the 7 minute time point after hypotonic stimulation are shown in **b** and **d** respectively. **e-f.** Representative western blots detecting SWELL1 and β-actin in 3T3-F442A adipocytes (**e**) and the corresponding mean densitometric data (**f**) obtained from treatment with vehicle (n=8), SN-401 (n=10), SN-406 (n=6) or Inactive 1 (n=3) at 10 µM for 96 hours. **g-h.** Representative western blots detecting SWELL1 and β-actin in 3T3-F442A adipocytes (**g**) and the corresponding mean densitometric data (**h**) obtained from treatment with vehicle (n=5), SN-401 (n=5), SN-406 (n=6), Inactive 1 (n=5) or Inactive 2 (n=4) at 1 µM for 96 hours. **i-j.** Representative immunostaining images demonstrating localization of endogenous SWELL1 in 3T3-F442A preadipocytes treated with vehicle or SN-401/SN-406/Inactive 1 at 10 µM for 48 hours (Scale bar – 20 µm) (**i**) and the corresponding quantification of mean SWELL1 membrane-versus cytoplasm-localized fraction obtained from vehicle (n=19), SN-401 (n=21), SN-406 (n=13 for 1 µM and 10 µM), or Inactive 1 (n=9 for 1 µM and n=13 for 10 µM) treated cells (**j**). **k-l.** Representative western blots detecting SWELL1, peNOS, eNOS, pAKT2, AKT2 and β-actin in HUVEC cells treated with either vehicle or 100 nM SN-406 for 96 hours (**k**) and their corresponding densitometric ratios (**l,** n= 6 each) respectively. **m-n.** Representative western blots detecting SWELL1, peNOS, eNOS, pAKT2, AKT2 and β-actin in HUVEC cells treated with either vehicle or 500 nM Inactive 1 for 96 hours (**m**) and their corresponding densitometric ratios (**n**, n= 6). **o-p.** Representative western blots detecting peNOS, eNOS and β-actin in HUVEC cells treated with either vehicle or 10, 30,100, 300 or 500 nM SN-401 for 96 hours (**o**, n=2 each) and the corresponding curve for dose dependent stimulation of peNOS/eNOS expression (**p**). Data are represented as mean ±SEM. Two-tailed unpaired t-test was used in **f** and **h** (compared to vehicle)**, l and n.** One-way ANOVA was used for **b, d and j**. Non-linear least square method was used to fit the dose response curve in **p**. *, ** and *** represents *p<0.05*, *p<0.01* and *p<0.001* respectively.

Next, we applied SWELL1-active (SN-401, SN-406), and Inactive (Inactive1, Inactive 2) compounds to differentiated 3T3-F442A adipocytes under basal culture conditions for 4 days and then measured SWELL1 protein after 6 h of serum starving. At both 10 and 1 μM, SN-401 and SN-406 markedly augment SWELL1 protein to levels 1.5-2.1-fold greater than vehicle-treated controls, while SWELL1-inactive congeners Inactive 1 and Inactive 2 do not significantly increase SWELL1 protein levels (**Fig. 6e-h**). SN-401 and SN-406 also enhanced plasma membrane (PM) localization of endogenous SWELL1 in preadipocytes compared to vehicle- or Inactive1 (**Fig. 6i&j; Supplementary Fig. S9a**), consistent with increased endoplasmic reticulum (ER) to plasma membrane trafficking of SWELL1. Notably, SN-401 and SN-406 are capable of augmenting both SWELL1 protein and trafficking at concentrations as low as 1 μM (**Fig. 6g-j; Supplementary Fig. S9a**), or an order of magnitude below the ∼10 μM concentration required for inhibiting activated SWELL1-LRRC8 (upon hypotonic stimulation). These findings are consistent with the results of SN-401/SN-406 I_Cl,SWELL_ inhibition when pre-applied to closed SWELL1-LRRC8 channels (**Fig. 6a-d**) and also with our observations that 500 nM SN-401 is sufficient to augment SWELL1 dependent insulin-AKT2-AS160 signaling in 3T3-F442A adipocytes (**Fig. 2i&j**). Similarly, SN-401 applied at 500 nM to human umbilical vein endothelial cells (HUVECs) increases eNOS phosphorylation, a downstream AKT target, and this effect is abrogated by small interfering/short hairpin mediated *SWELL1* knockdown in HUVECs, supporting a SWELL1 dependent mechanism (**Supplementary Fig. S9b-d**). Furthermore, SN-406 applied to HUVECs at 100 nM is sufficient to induce SWELL1 1.5-fold, basal pAKT2 2.8-fold, and downstream p-eNOS 5.5-fold as compared to vehicle (**Fig. 6k&l)**, while Inactive 1 has no effect at 500 nM (**Fig. 6m&n**). Indeed, SN-401 exhibits dose-dependent induction p-eNOS in HUVECs with an EC_50_ of 21 nM (**Fig. 6o&p**).

### SWELL1-active compounds prevent reductions in SWELL1 protein and rescue SWELL1 dependent islet insulin secretion under glucolipotoxic conditions

We next asked whether endoplasmic reticulum (ER) stress associated with glucolipotoxicity in metabolic syndrome may promote SWELL1 protein degradation, and thereby reduce I_Cl,SWELL_ and SWELL1 protein in T2D (**Fig. 1**). In this context, we hypothesized that SN-401 and SN-406 might assist with SWELL1-LRRC8 assembly and rescue SWELL1-LRRC8 from degradation. To test this concept *in vitro*, we first treated 3T3-F442A adipocytes with either vehicle, SN-401, SN-406 or Inactive 2, and then subjected these cells to 1 mM palmitate + 25 mM glucose to induce to glucolipotoxic stress (**Fig. 7a&b**). We found that SWELL1 protein was reduced by 50% upon palmitate/glucose treatment, consistent with ER stress-mediated SWELL1 degradation, and this reduction was entirely prevented by both SWELL1-active SN-401 and SN-406, but not by SWELL1-inactive 2 (**Fig. 7a&b**). These data are consistent with the notion that SN-401 and SWELL1-active congeners are functioning to stabilize SWELL1-LRRC8 assembly and signaling under glucolipotoxic conditions associated with T2D and metabolic syndrome.

**Figure 7.**
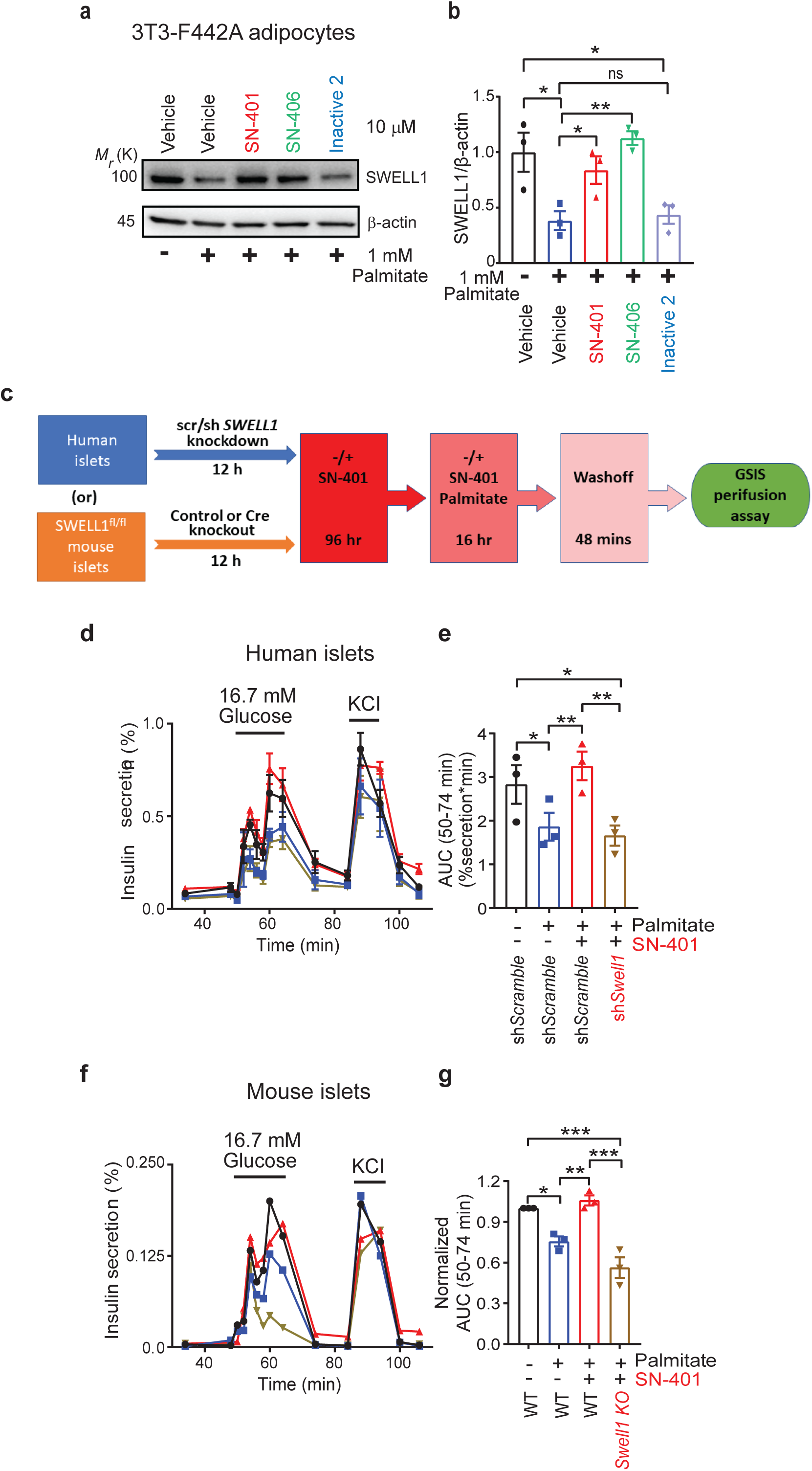
SWELL1-active compounds prevent reductions in SWELL1 protein and rescue SWELL1 dependent islet insulin secretion under glucolipotoxic conditions. **a-b.** Representative western blots detecting SWELL1 and β-actin in 3T3-F442A adipocytes pre-treated with either vehicle or SN-401/SN-406/Inactive 2 (10 µM) for 96 hours and subsequently treated with -/+ palmitate in the absence or presence of compounds for 16 hours (**a**, n=3 in each condition) and corresponding densitometric ratio for SWELL1/β-actin (**b**). **c.** Schematic for glucose stimulated insulin secretion (GSIS) perifusion assay in human or mouse islets. **d-e.** GSIS perifusion assay of islets obtained from cadaveric human islets transduced with either adenoviral short hairpin control (shScramble) or SWELL1 (shSWELL1) for 12 hours, then pre-treated with either vehicle or SN-401 (10 µM) for 96 hours and subsequently treated with -/+ palmitate or palmitate with -/+ SN-401 (n = 3 each) for 16 hours (**d**) and the corresponding area under the curve (AUC) (**e**). **f-g.** GSIS perifusion assay of WT and β-cell SWELL1 KO islets obtained by isolation of islets from floxed-SWELL1^fl/fl^ mouse and transducing either with adenoviral control (WT) or Cre-recombinase (SWELL1 KO) for 12 hours, then pre-treated with either vehicle or SN-401 (10 µM) for 96 hours and subsequently treated with -/+ palmitate or palmitate with -/+ SN-401 (n = 3 each) for 16 hours (**f**) and the corresponding area under the curve (AUC) (**g**). Data are represented as mean ±SEM except for **f** where a representative trace was shown for illustrative purposes. Two-tailed unpaired t-test was used in **b.** One-way ANOVA was used for **e** and **g.** *, ** and *** represents *p<0.05*, *p<0.01* and *p<0.001* respectively and ‘ns’ indicates the difference was not significant.

To examine whether these protective effects of SN-401 under glucolipotoxic conditions is also capable of rescuing islet insulin secretion, and whether this effect occurs via target engagement, we measured dynamic glucose-stimulated insulin secretion (GSIS) by perifusion in human and murine islets +/- palmitate +/- SN-401 and +/- SWELL1 as outlined in **Fig. 7c**. In both human and murine islets, we found that 16 hours of 1mM palmitate treatment reduces GSIS compared to baseline (**Fig. 7d-g**, **Supplementary Table. S2**). However, when islets are treated with SN-401 (10 µM) for 4 days prior to palmitate treatment, then maintained during palmitate treatment, and subsequently SN-401/palmitate washed off during GSIS, insulin secretion is normalized (**Fig. 7d-g**, **Supplementary Table. S2**). Importantly, this SN-401 mediated GSIS normalization under glucolipotoxic conditions is SWELL1 dependent, since this rescue is completely abrogated in SWELL1 KD human islets (**Fig. 7d&e**, **Supplementary Table. S2**) and β-cell targeted SWELL1 KO murine islets (**Fig. 7f&g**).

### SWELL1-active SN-401 congeners improve glycemic control in murine T2D

To determine if the effects of SN-401 observed *in vivo* in T2D mice are attributable to SWELL1-LRRC8 binding, as opposed to off-target effects, we next measured fasting blood glucose and glucose tolerance in HFD T2D mice treated with either SWELL1-active SN-403 or SN-406 as compared to SWELL1-inactive Inactive 1 (all at 5 mg/kg/day x 4 days). In mice treated with HFD for 8 weeks, SN-403 significantly reduced fasting blood glucose and improved glucose tolerance compared to Inactive 1 (**Fig. 8a**). In cohorts of mice raised on HFD for 12-18 weeks, with more severe obesity-induced T2D, SN-406 also markedly reduced fasting blood glucose and improved glucose tolerance (**Fig. 8b**). Similarly, in a separate experiment, SN-406 significantly improved glucose tolerance in HFD T2D mice, compared to Inactive 1 (**Fig. 8c**), and this is associated with a trend toward improved insulin sensitivity based on the Homeostatic Model Assessment of Insulin Resistance (HOMA-IR) ^54^ (**Fig. 8d**), and significantly augmented insulin secretion in perifusion GSIS (**Fig. 8e**). Finally, based on the GTT AUC, SN-407 also improved glucose tolerance in T2D KKA^y^ mice, compared to Inactive 1 (**Fig. 8f**) and increased GSIS (**Fig. 8g**). These data reveal the *in vivo* anti-hyperglycemic action of SN-401 and its bioactive congeners require SWELL1-LRRC8 binding and thus supports the notion of SWELL1 on-target activity *in vivo*.

**Figure. 8.**
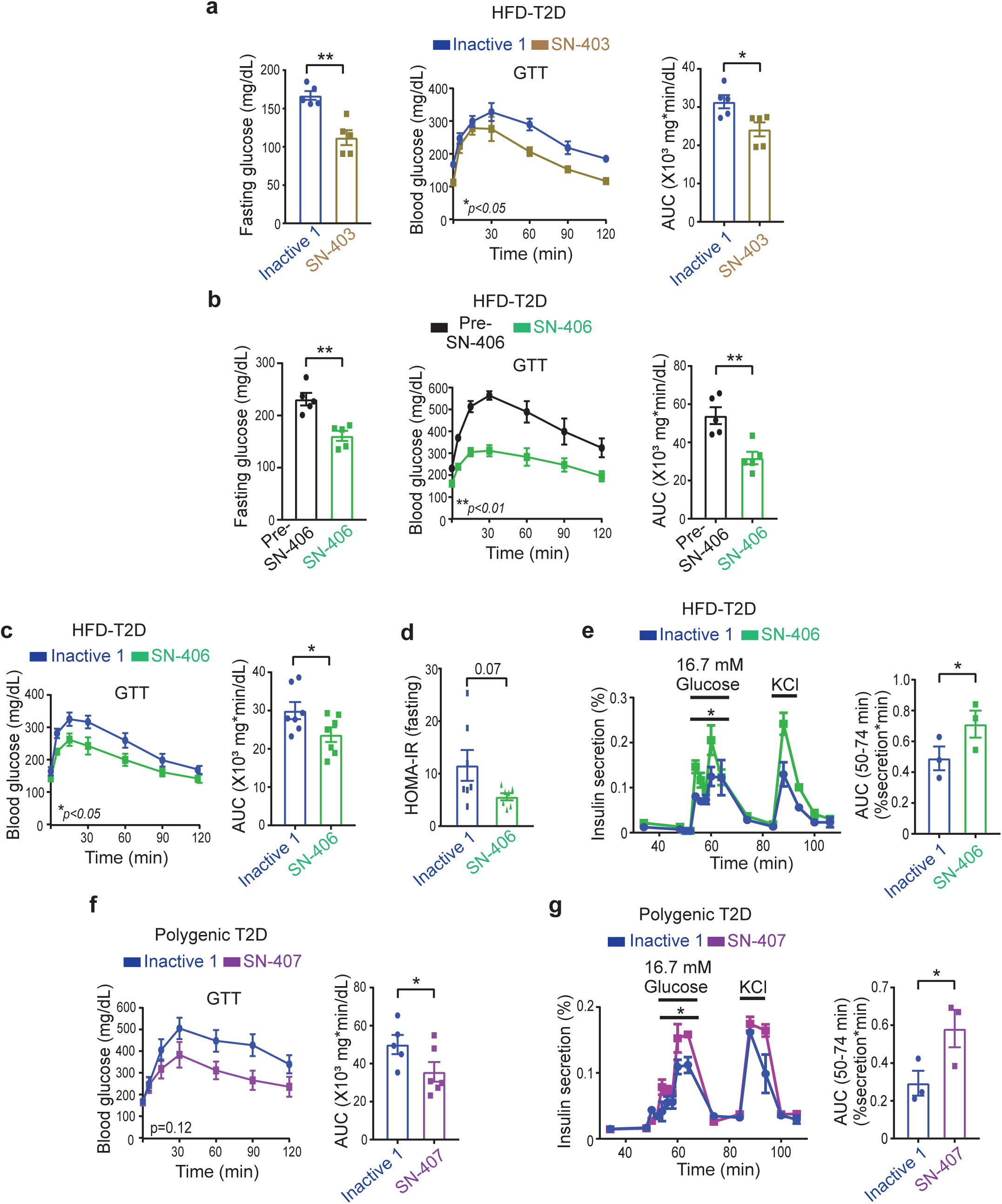
SWELL1-active SN-401 congeners improve glycemic control in murine T2D models. **a.** Fasting glucose levels, GTT and its corresponding area under the curve (AUC) of 8 week HFD-fed mice treated with either SWELL1-inactive 1 or SWELL1-active SN-403 (5 mg/kg i.p) for 4 days (n = 5 in each group). **b.** Fasting glucose levels, GTT and its corresponding AUC of 12 weeks HFD-fed mice pre- and post-treatment with SN-406 (5 mg/kg i.p) for 4 days (n = 5 in each group). **c.** GTT and corresponding AUC of 12 weeks HFD-fed mice treated with either SWELL1-inactive 1 or SWELL1-active SN-406 (5 mg/kg i.p) for 4 days (n = 7 in each group) and **(d)** the corresponding HOMA-IR index. **e.** Glucose-stimulated insulin secretion (GSIS) perifusion assay of islets isolated from mice in **c** (left) and the corresponding area under the curve (right). **f.** GTT and corresponding AUC of polygenic-T2D KKA^y^ mice treated with either SWELL1-inactive 1 (n=5) or SWELL1-active SN-407 (n=6) (5 mg/kg i.p) for 4 days. **g.** Glucose-stimulated insulin secretion (GSIS) perifusion assay from islets isolated from mice in **f** (left) and the corresponding area under the curve (right). Data are represented as mean ±SEM. Two-tailed unpaired t-test was used in **a**,**c-d, e-g** for FG, GTT AUC, GSIS AUC and HOMA-IR. **d**. Paired t-test was used in **b** for FG and GTT AUC. Two-way ANOVA was used in **a-c** and **f** for GTTs. Statistical significance is denoted by *, ** and *** representing *p<0.05*, *p<0.01* and *p<0.001* respectively.

An important feature of the hypothesized mechanism of action of SN-40X is that these active compounds bind to SWELL1-LRRC8 channel complexes *in vivo* in the ∼100-500 nM range to augment SWELL1-dependent signaling (**Fig 2i&j**; **Fig. 6k-p; Supplementary Fig. S9b-d**) without achieving the serum concentrations necessary for open channel SWELL1 current inhibition (∼5-10 uM)^31^, followed by unbinding. As such, SWELL1 function may be rescued without significant SWELL1-LRRC8 or VRAC inhibition. Consistent with this hypothesis, *in vivo* pharmacokinetics (PK) of SN-401 and SN-406 in mice following i.p. or p.o. administration of 5 mg/kg of SN-401 or SN-406 reveal plasma concentrations that either transiently approach (**Supplementary Fig. S10a**, i.p. dosing), or remain well below I_Cl,SWELL_ inhibitory concentrations (**Supplementary Fig. S10b**, p.o. dosing) while exceeding concentrations sufficient for induction of SWELL1 signaling activity (> ∼100 nM) for 8-12 hours. Importantly, SN-401 PK in a target tissue, adipose, also closely tracks serum concentrations via both i.p. and p.o. administration (**Supplementary Fig. S10c)**. Finally, these *in vivo* PK studies demonstrate that SN-401 has high oral bioavailability (AUC_p.o._/AUC_i.v._ = 79%, **Supplementary Table S7**), and when administered via oral gavage to HFD-fed T2D C57 mice at 5 mg/kg/day SN-401 fully retains *in vivo* therapeutic efficacy (**Supplementary Fig. S10d).** Collectively, these PK data reveal that appropriate SN-401 concentrations are attained to achieve the observed therapeutic effect, while remaining insufficient to inhibit activated VRAC.

## Discussion

Our current working model is that the transition from compensated obesity (pre-diabetes, normoglycemia) to decompensated obesity (T2D, hyperglycemia) reflects, among other things, a relative reduction in SWELL1 protein expression and signaling in peripheral insulin-sensitive tissues^29, 32^ and in pancreatic β-cells^55, 56^ – metabolically phenocopying SWELL1-loss-of-function models^24, 27–29^. This contributes to the combined insulin resistance and impaired insulin-secretion associated with poorly-controlled T2D and hyperglycemia. SWELL1 forms a macromolecular signaling complex that includes heterohexamers of SWELL1 and LRRC8b-e^22, 23^, with stoichiometries that likely vary from tissue to tissue. We propose SWELL1-LRRC8 signaling complexes are inherently unstable, and thus a proportion of complexes succumb to disassembly and degradation. Glucolipotoxicity and ensuing ER stress associated with T2D states^57–59^ provide an unfavorable environment for SWELL1-LRRC8 complex assembly, contributing to SWELL1 degradation and reductions in SWELL1 protein and SWELL1-mediated I_Cl,SWELL_ observed in T2D. We speculate that small molecules SN-401 and active congeners (SN-40X) serve as pharmacological chaperones^49^ to stabilize formation of the SWELL1-LRRC8 complex. This reduces SWELL1 degradation, and enhances the passage of SWELL1-LRRC8 heteromers through the ER and Golgi apparatus to the plasma membrane – thereby rectifying the SWELL1-deficient state in multiple metabolically important tissues in the setting of metabolic syndrome to improve glycemic control via both insulin sensitization^24, 25, 29^ and secretion^27, 28^ mechanisms. Indeed, the concept of small molecule inhibitors acting as therapeutic molecular chaperones^49^ to support the folding, assembly and trafficking of proteins (including ion channels) has been demonstrated for Niemann-Pick C disease^50, 51^ and congenital hyperinsulinism (SUR1-K_ATP_ channel mutants)^60–62^. Similarly, in the case of congenital hyperinsulinism, the SUR1-K_ATP_ chemical chaperones are also themselves, paradoxically, K_ATP_ channel inhibitors^60–62^. Also, this therapeutic mechanism is analogous to small molecule correctors for another chloride channel, CFTR (VX-659/VX-445, Vertex Pharmaceuticals)^52, 53^, which is proving to be a breakthrough therapeutic approach^63, 64^ for cystic fibrosis. *In vivo*, we hypothesize SN-40X compounds bind to SWELL1-LRRC8 complexes in the closed state, within the concentration range between C_max_ and ∼100 nM. This shifts the balance toward maintaining stable SWELL1-LRRC8 complexes to preserve normal levels and localization (trafficking) within the T2D glucolipotoxic milieu. SN-40X may then unbind from the SWELL1-LRRC8 complex, thereby restoring insulin signaling in target tissues, and permitting SWELL1-mediated β-cell insulin secretion. This seemingly paradoxical mechanism may rely on the *phasic* SN-40X concentrations observed *in vivo* (see *in vivo* PK data) to allow for SN-401 binding, resultant chaperone-mediated rescue, followed by unbinding; as opposed to *tonic* SN-40X-SWELL1 binding. Another prediction of this model is that lower-affinity SN-40X compounds may be preferable to very high-affinity congeners, to provide the appropriate pharmacodynamics required for unbinding, and optimal therapeutic efficacy. Indeed, this mechanism is reminiscent of the paradoxical use of insulin secretagogue sulfonylurea receptor inhibitors as pharmacological chaperones to rescue KATP mutants in congenital hyperinsulinism by binding (and inhibiting) these mutant KATP channels^60–62^, and then unbinding, thereby favoring lower affinity inhibitors: tolbutamide and carbamazepine, over glibenclamide.

Through structure activity relationship (SAR), *in silico* molecular docking studies and cryo-EM studies, we identified hotspots on opposing ends of the SN-401 molecule that interact with separate regions of the SWELL1-LRRC8 complex: the carboxylate group with R103 in multiple LRRC8 subunits at a constriction in the pore, and the cyclopentyl group within the hydrophobic cleft formed by adjacent LRRC8 monomers; functioning like a molecular staple or tether to bind loosely associated SWELL1-LRRC8 monomers (especially in the setting of T2D) into a more stable hexameric structure. Indeed, the cryo-EM structure obtained in lipid nanodiscs of SN-401^30^ and novel derivative SN-407 supports hypothesized binding models of SN-40X with SWELL1 homomer.

Another advantage provided by SAR studies was identification and synthesis of SN-401 congeners that removed (Inactive 1 and 2) or enhanced (SN-403/406/407) SWELL1-binding, as these provided powerful tools to query SWELL1-on target activity directly *in vitro* and *in vivo*, and also validated the proof-of-concept for developing novel SN-401 congeners with enhanced efficacy. Indeed, this approach was necessary to prove SWELL1-LRRC8 on-target activity of the SN-40X series *in vivo*, because SWELL1-LRRC8 is expressed broadly in numerous insulin-sensitive tissues and in islet cells. As the global SWELL1/LRRC8a KO mouse is essentially embryonically lethal ^20^ testing SN-40X compounds in global SWELL1^-/-^ mice is not possible, and generating multi-tissue (adipose, liver, skeletal muscle, β-cell) SWELL1 KO mice is outside the scope of the current study. Therefore, using the SAR to generate SWELL1-LRRC8 inactive compounds (Inactive 1 and 2) as a negative control provided an alternative approach to prove *in vivo* on-target activity on a broadly expressed signaling molecule. In addition to this medicinal chemistry approach employed both *in vitro* and *in vivo* to test on target activity, we found that SN-40X mediated induction of AKT-AS160 and AKT-eNOS signaling requires SWELL1 in cultured adipocytes and HUVECs, respectively. Moreover, SN-401 mediated rescue of islet insulin secretion under gluco-lipotoxic conditions *in vitro* also requires SWELL1. Finally, it is important to note that the studies demonstrating promiscuity of SN-401/DCPIB with other ion channel targets all applied DCPIB at ∼10-200 µM^65–70^. This is 100-200-fold higher than the concentrations required to potentiate SWELL1-dependent signaling *in vitro* (**Fig. 2i&j**; **Fig. 6k-p; Supplementary Fig. S9b-d**), and similarly higher than SN-401 and SN-406 concentrations predominantly attained *in vivo* (**Supplementary Fig. S10a-c**) to achieve a therapeutic effect (**Fig. 3&8**). Accordingly, these studies are not applicable with respect to putative off-target mechanisms for the therapeutic effects observed from SN-40X compounds.

SWELL1-LRRC8 complexes are broadly expressed in multiple tissues, and consist of unknown combinations of SWELL1, LRRC8b, LRRC8c, LRRC8d and LRRC8e, indicating SWELL1 complexes may be enormously heterogenous. However, SWELL1-LRRC8 stabilizers like SN-401 may be designed to target many, if not all, possible channel complexes since all will contain the elements necessary for SN-401 binding: at least one R103 (from the requisite SWELL1 monomer: carboxyl group binding site), and the nature of the hydrophobic cleft (cyclopentyl binding site), which is conserved among all LRRC8 monomers. Indeed, traced glucose clamps did reveal insulin sensitization effects in multiple tissues, including adipose, skeletal muscle, liver and heart. The increased glucose-uptake in heart is particularly interesting, since this may provide salutary effects on cardiac energetics that could favorably impact both systolic (HFrEF) and diastolic (HFpEF) function in diabetic cardiomyopathy, and thereby potentially improve cardiac outcomes in T2D, as observed with SGLT2 inhibitors^71–76^.

The current study provides an initial proof-of-concept for pharmacological induction of SWELL1 signaling using SWELL1 modulators (SN-40X) to treat metabolic diseases at multiple homeostatic nodes, including adipose, skeletal muscle, liver, and pancreatic β-cell, whereby SN-40X compounds function to restore both insulin-sensitivity and insulin secretion. Hence, SN-401 may represent a tool compound from which a novel drug class may be derived to treat T2D, NASH, and other metabolic diseases.

## Methods

### Patients

Human islets (Integrated Islet Distribution Program, Prodo Laboratories and Alberta Diabetes Institute Islet Core) and adipocytes were obtained and cultured as described previously^24, 27^. The patients involved in the study were anonymous and information such as gender, age, HbA1c, glucose levels and BMI only were available to the research team. The study was approved and carried out as per the guidelines of the University of Iowa and Washington University Institutional Review Board (IRB).

### Animals

All experimental procedures involving mice were approved by the Institutional Animal Care and Use Committee of the University of Iowa and Washington University at St. Louis. All C57BL/6 mice involved in this study were purchased from Charles River Labs. Both KK.Cg-Ay/J (KKA^y^) and KK.Cg-Aa/J (KKA^a^) mice involved in study were gender and age-matched mice obtained from Jackson Labs (Stock No: 002468) and bred up for experiments. The mice were fed *ad libitum* with either regular chow (RC) or high-fat diet (Research Diets, Inc., 60 kcal% fat) with free access to water and housed in a light-, temperature- and humidity-controlled room. For high-fat diet (HFD) studies, only male mice were used and were started on HFD regimen at the age of 6-9 weeks. For all experiments involving KKA^y^ and KKA^a^ mice, both males and females were used at approximately 50/50 ratio. In all experiments involving mice, investigators were kept blinded both during the experiments and subsequent analysis.

### Small molecule treatment

All compounds were dissolved in Kolliphor® EL (Sigma, #C5135). Either vehicle (Kolliphor® EL), SN-401 (DCPIB, 5 mg/kg of body weight/day, Tocris, D1540), SN-403, SN-406, SN-407 or Inactive 1 were administered i.p. as indicated using 1cc syringe/26G X 1/2 inch needle daily for 4-10 days, and in one experiment, SN-401 was administered daily for 8 weeks. SN-401, formulated as above, was also administered by oral gavage at 5 mg/kg/day for 5 days using a 20G x 1.5 inch reusable metal gavage needle.

### Adenovirus

Human adenoviruses type 5 with hLRRC8A/SWELL1-shRNA (Ad5-mCherry-U6-hLRRC8A/SWELL1-shRNA, 2.2 × 10^10^ PFU/ml), a scrambled non-targeting control (Ad5-U6-scramble-mCherry, 1 × 10^10^ PFU/ml), Ad5-CAG-LoxP-stop-LoxP-3XFlag-SWELL1 (1X 10^10^ PFU/ml), β-cell-targeted adenovirus type 5 with Ad5-RIP2-GFP (4.1 × 10^10^ PFU/ml), GCaMP6s (Ad5-RIP1-GCaMP6s, 4.9 × 10^10^ PFU/ml), and GCaMP6s-2A-iCre (Ad5-GCaMP6s-RFP-2A-Cre, 5.8 × 10^10^ PFU/ml) were obtained from Vector Biolabs. Adenovirus type 5 with Ad5-CMV-Cre-eGFP (8 × 10^10^ PFU/ml) and Ad5-CMV-Cre-mCherry (3 × 10^10^ PFU/ml) were obtained from the University of Iowa Viral Vector Core.

### Cell culture

Wildtype (WT) and SWELL1 knockout (KO) 3T3-F442A (Sigma-Aldrich) cells were cultured and differentiated as described previously^24^. Preadipocytes were maintained in 90% DMEM (25 mM D-Glucose and 4 mM L-Glutamine) containing 10% fetal bovine serum (FBS) and 100 IU penicillin and 100 µg/ml streptomycin on collagen-coated (rat tail type-I collagen, Corning) plates at 37°C and 5% CO_2_. Upon reaching confluency, the cells were differentiated in the above-mentioned media supplemented with 5 µg/ml insulin (Cell Applications) and replenished every other day with the differentiation media. For insulin signaling studies on WT and KO adipocytes with or without SWELL1 overexpression (O/E), the cells were differentiated for 10 days and transduced with Ad5-CAG-LoxP-stop-LoxP-SWELL1-3XFlag virus (MOI 12) on day 11 in 2% FBS containing differentiation medium. To induce the overexpression, Ad5-CMV-Cre-eGFP (or mcherry) (MOI 12) was added on day 13 in 2% FBS containing differentiation medium. The cells were then switched to 10% FBS containing differentiation medium from day 15 to 17. On day 18, the cells were starved in serum free media for 6 hours and stimulated with 0 and 10 nM insulin for 15 min. Either Ad5-CAG-LoxP-stop-LoxP-SWELL1-3XFlag or Ad5-CMV-Cre-eGFP (or mcherry) virus transduced cells alone were used as controls. Based on GFP/mcherry fluorescence, viral transduction efficiency was ∼90%.

For SN-401 treatment and insulin signaling studies in 3T3-F442A preadipocytes, the cells were incubated with either vehicle (DMSO) or 10 µM SN-401 for 96 hours. The cells were serum starved for 6 hours with vehicle (DMSO) or SN-401 and washed with PBS three times and stimulated with 0, 3 and 10 nM insulin containing media for 15 minutes prior to collecting lysates. In the case of 3T3-F442A adipocytes, the WT and KO cells were treated with either vehicle (DMSO), 1 or 10 µM SN-40X (after 7-11 days of differentiation) for 96 hours and then stimulated with 0 and 10 nM insulin/serum containing media with vehicle (DMSO) or SN-40X for 15 minutes for SWELL1 detection. For AKT and AS160 signaling, the WT and KO cells were treated with either vehicle (DMSO) or 500 nM SN-401 for 96 hours and serum starved in the presence of vehicle or SN-401 (500 nM) for 6 hours. The cells were washed twice in hypotonic buffer (240 mOsm; 90 mM NaCl, 5 mM NaHCO_3_, 4.8 mM KCl, 1.2 mM KH_2_PO_4_, 2.5 mM CaCl_2_, 2.4 mM MgSO_4_, 10 mM HEPES and 25 mM Glucose, pH 7.4) and then incubated at 37 °C in hypotonic buffer for 10 minutes followed by a serum free media wash and subsequent stimulation with insulin/serum containing media for 30 minutes at 37°C without SN-401 (or vehicle). To simulate gluco-lipotoxicity, 8 mM sodium palmitate was dissolved in 18.4% fatty-acid free BSA at 37 °C in DMEM medium with 25 mM glucose to obtain a conjugation ratio of 1:3 palmitate:BSA^77^. As described above, the 3T3-F442A adipocytes were incubated with vehicle or SN-401, SN-406, Inactive 2 at 10 µM for 96 hours and treated with 1 mM palmitate for additional 16 hours in the presence of compounds and lysates were collected and further processed.

HUVECs were purchased from ATCC and were grown in M199 growth media supplemented with 20% FBS, 0.05g Heparin Sodium Salt (Alfa Aesar) and 15 mg ECGS (Millipore Sigma). Cells were cultured on 1% of gelatin coated plates at 37°C and 5% CO_2_. For SN-40X stimulated eNOS and AKT signaling assays, HUVECs were treated for 96 hours with the respective compounds and serum starved overnight (+DMSO or +SN-40X) for 16 hours in M199 media plus 1% FBS (Atlanta Biological). After the serum starve, HUVECs were returned to normal growth media for 30 minutes (+DMSO or +SN-40X) prior to lysate collection. Small interfering RNA (siRNA) mediated knockdown was adapted from^78^. Briefly, HUVECs were transfected with either a silencer select siRNA with si-SWELL1 (Cat#4392420, sense: GCAACUUCUGGUUCAAAUUTT antisense: AAUUUGAACCAGAAGUUGCTG, Invitrogen) or a non-targeting control silencer select siRNA (Cat# 4390846, Invitrogen) upon reaching 90-95% confluency. siRNA was transfected twice, 24 and 72 hours after initial seeding of HUVECs. Each siRNA was combined with Opti-MEM (285.25 µl, Cat#11058-021, Invitrogen) siPORT™ amine (8.75 µl, Cat#AM4503, Invitrogen) and the silencer select siRNA (6 µl) in a final volume of 300 µl. HUVECs were transfected for 4 hours at 37°C in 1% FBS containing DMEM media. For short hairpin RNA (shRNA) mediated knockdown approach, HUVECs were transduced with either human adenovirus type 5 targeting SWELL1 (Ad5-shSWELL1) or a scrambled non-targeting control (Ad5-shSCR) at a multiplicity of infection (MOI) of 50 for 24 hours at 37°C upon reaching 70% confluency. The cells were then washed with DMEM media and transduced a second time, with fresh virus, with a MOI of 25 for 12 hours at 37°C. The SN-40X compounds were present in the culture media throughout the transduction for both the si/sh RNA mediated knockdown approaches and after the final transduction step, the cells were serum starved as described above for HUVECs and lysates were collected. HEK-293 (ATCC® CRL-1573™) cells were maintained in 90% DMEM (25 mM D-Glucose and 4 mM L-Glutamine) containing 10% fetal bovine serum (FBS) and 100 IU penicillin and 100 µg/ml streptomycin.

### Molecular docking

SN-401 and its analogs were docked into the expanded state structure of a LRRC8A-SN-401 homo-hexamer in MSP1E3D1 nanodisc (PDB ID: 6NZZ) using Molecular Operating Environment (MOE) 2016.08 software package [Chemical Computing Group (Montreal, Canada)]. The 3D structure obtained from PDB (PDB ID: 6NZZ) was prepared for docking by first generating the missing loops using the loop generation functionality in Yasara software package followed by sequentially adding hydrogens, adjusting the 3D protonation state and performing energy minimization using Amber10 force-field in MOE. The ligand structures to be docked were prepared by adjusting partial charges followed by energy minimization using Amber10 force-field. The site for docking was defined by selecting the protein residues within 5Å from co-crystallized ligand (SN-401). Docking parameters were set as Placement: Triangle matcher; Scoring function: London dG; Retain Poses: 30; Refinement: Rigid Receptor; Re-scoring function: GBVI/WSA dG; Retain poses: 5. Binding poses for the compounds were predicted using the above validated docking algorithm.

### Chemical Synthesis

#### General Information

All commercially available reagents and solvents were used directly without further purification unless otherwise noted. Reactions were monitored either by thin-layer chromatography (carried out on silica plates, silica gel 60 F_254_, Merck) and visualized under UV light. Flash chromatography was performed using silica gel 60 as stationary phase performed under positive air pressure. ^1^H NMR spectra were recorded in CDCl_3_ on a Bruker Avance spectrometer operating at 300 MHz at ambient temperature unless otherwise noted. All peaks are reported in ppm on a scale downfield from TMS and using the residual solvent peak in CDCl_3_ (H δ = 7.26) or TMS (δ = 0.0) as an internal standard. Data for ^1^H NMR are reported as follows: chemical shift (ppm, scale), multiplicity (s =singlet, d = doublet, t = triplet, q = quartet, m = multiplet and/or multiplet resonances, dd = double of doublets, dt = double of triplets, br = broad), coupling constant (Hz), and integration. All high-resolution mass spectra (HRMS) were measured on Waters Q-Tof Premier mass spectrometer using electrospray ionization (ESI) time-of-flight (TOF).

**Scheme 1:**
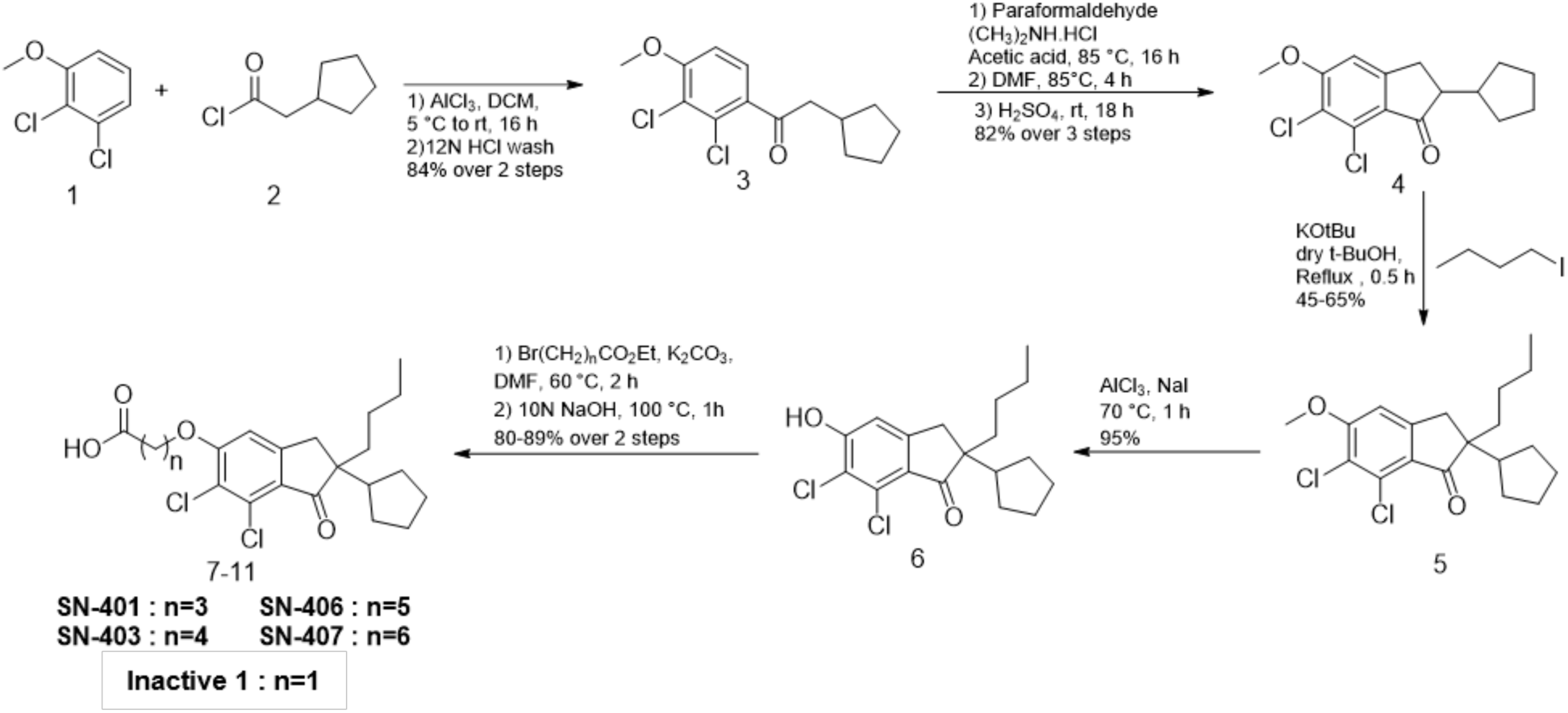

*2-cyclopentyl-1-(2,3-dichloro-4-methoxyphenyl)ethan-1-one* (**3**): To a stirring solution of aluminum chloride (13.64 g, 102 mmol, 1.1 equiv.) in dichloromethane (250 ml) at 0 ℃ was added cyclopentyl acetyl chloride (15 g, 102 mmol, 1.1 equiv.) and the resulting solution was allowed to stir at 0 ℃ under nitrogen atmosphere for 10 minutes. To this was added a solution of 2, 3-dicholoro anisole (16.46 g, 92.9 mmol, 1 equiv.) in dichloromethane (50 ml) at 0 ℃ and the resulting solution was allowed to warm to room temperature and stirred for 16 hours. Once complete, the reaction was added to cold concentrated hydrochloric acid (100 ml) followed by extraction in dichloromethane (150 ml × 3). The organic fractions were pooled, concentrated and purified by silica gel chromatography using 0-15% ethyl acetate in hexanes as eluent to furnish compound **3** as white solid (22.41 g, 84%). ^1^H NMR (300 MHz, CDCl_3_) δ 7.39 (d, *J* = 8.7 Hz, 1H), 6.89 (d, *J* = 8.7 Hz, 1H), 3.96 (s, 3H), 2.96 (d, *J* = 7.2 Hz, 2H), 2.38 – 2.21 (m, 1H), 1.92 – 1.75 (m, 2H), 1.69 – 1.46 (m, 4H), 1.28 – 1.05 (m, 2H). HRMS (ESI), *m*/*z* calcd for C_14_H_17_Cl_2_O_2_ [M + H]^+^ 287.0605, found 287.0603.

*6,7-dichloro-2-cyclopentyl-5-methoxy-2,3-dihydro-1H-inden-1-one* (**4**): To 2-cyclopentyl-1-(2,3-dichloro-4-methoxyphenyl)ethan-1-one (**3**) (21.5 g, 74.8 mmol, 1 equiv.) in a round bottom flask was added paraformaldehyde (6.74 g, 224.5 mmol, 3 equiv.), dimethylamine hydrochloride (30.52 g, 374 mmol, 5 equiv.) and acetic acid (2.15 ml) and the resulting mixture was allowed to stir at 85 ℃ for 16 hours. To the reaction was then added dimethylformamide (92 ml) and the resulting solution was allowed to stir at 85 ℃ for 4 hours. Once complete, the reaction was diluted with ethyl acetate and then washed with 1N hydrochloric acid. The organic fractions were collected and concentrated under vacuum and used for next step without purification. To the concentrated product in a round bottom flask was added cold concentrated sulfuric acid (120 ml) at 0 ℃ and the resulting solution was allowed to stir at room temperature for 18 hours. Once complete, the reaction was diluted with cold water and extracted thrice with ethyl acetate (100 ml). The organic fractions were pooled, concentrated and purified by silica gel chromatography using 0-15% ethyl acetate in hexanes as eluent to furnish compound **4** as beige solid (18.36 g, 82%). ^1^H NMR (300 MHz, CDCl_3_) δ 6.88 (s, 1H), 4.00 (s, 3H), 3.16 (dd, *J* = 18.1, 8.7 Hz, 1H), 2.80 (d, *J* = 14.4 Hz, 2H), 2.43 – 2.22 (m, 1H), 1.96 (s, 1H), 1.73 – 1.48 (m, 5H), 1.46 – 1.33 (m, 1H), 1.17 – 1.00 (m, 1H). LRMS (ESI), *m*/*z* calcd for C_15_H_17_Cl_2_O_2_ [M + H]^+^ 299.0605, found 299.0614.

*2-butyl-6,7-dichloro-2-cyclopentyl-5-methoxy-2,3-dihydro-1H-inden-1-one* (**5**): A stirring suspension of **4** (23 gm, 76.8 mmol, 1 equiv.) in anhydrous tert-butanol (220 ml) was allowed to reflux at 95 ℃ for 30 minutes. To the resulting solution was added potassium tert-butanol (1M in tert-butanol) (84 ml, 84.5 mmol, 1.1 equiv.) and the resulting solution was refluxed for 30 minutes. The reaction was then cooled to room temperature followed by addition of iodobutane (44.2 ml, 384 mmol, 5 equiv.) and the reaction was then allowed to reflux for additional 60 minutes. The reaction was allowed to cool, concentrated and purified by silica gel chromatography using 0-10% ethyl acetate in hexanes as eluent to furnish compound **5** as clear oil (17.75 g, 65%). ^1^H NMR (300 MHz, CDCl_3_) δ 6.89 (s, 1H), 4.09 – 3.90 (m, 3H), 2.98 – 2.70 (m, 2H), 2.36 – 2.18 (m, 1H), 1.89 – 1.71 (m, 2H), 1.58 – 1.42 (m, 5H), 1.33 – 1.09 (m, 4H), 1.09 – 0.94 (m, 2H), 0.93 – 0.73 (m, 4H). HRMS (ESI), *m*/*z* calcd for C_19_H_25_Cl_2_O_2_ [M + H]^+^ 355.1231, found 355.1231.

2-butyl-6,7-dichloro-2-cyclopentyl-5-hydroxy-2,3-dihydro-1H-inden-1-one (**6**): To **5** (3.14 g, 8.87 mmol, 1 equiv.) was added aluminum chloride (2.36g, 17 mmol, 2 equiv.) and sodium iodide (2.7 g, 17 mmol, 2 equiv.) and the resulting solid mixture was triturated and allowed to stir at 70 ℃ for 60 minutes. Once complete, the reaction was diluted with dichloromethane and washed with aqueous saturated sodium thiosulfate solution. The organic fractions were collected and concentrated to give a beige solid which was then washed multiple times with hexanes to provide compound **6** as white solid (2.87 g, 95%). ^1^H NMR (300 MHz, CDCl_3_) δ 7.03 (s, 1H), 6.32 (s, 1H), 2.97 – 2.73 (m, 2H), 2.36 – 2.17 (m, 1H), 1.88 – 1.68 (m, 2H), 1.62 – 1.39 (m, 6H), 1.31 – 1.11 (m, 3H), 1.08 – 0.97 (m, 2H), 0.97 – 0.87 (m, 1H), 0.83 (t, *J* = 7.3 Hz, 3H). HRMS (ESI), *m*/*z* calcd for C_18_H_23_Cl_2_O_2_ [M + H]^+^ 341.1075, found 341.1089.

*2-((2-butyl-6,7-dichloro-2-cyclopentyl-1-oxo-2,3-dihydro-1H-inden-5-yl)oxy)acetic acid* (**7**) (Inactive 1): To a stirring solution of **6** (170 mg, 0.50 mmol, 1 equiv.) in anhydrous dimethylformamide (1 ml) was added potassium carbonate (76 mg, 0.56 mmol, 1.1 equiv.) and ethyl 2-bromoacetate (61 ml, 0.56 mmol, 1.1 equiv.) and the reaction was allowed to stir at 60 ℃ for 2 hours. Once complete, to the reaction was added 4 N NaOH (1 ml) and the reaction was allowed to stir at 100 ℃ for 60 minutes. Once complete, reaction was concentrated and purified by column chromatography using 0-10% methanol in dichloromethane as eluent to provide **Inactive 1** as a clear solid (173 mg, 87%). ^1^H NMR (300 MHz, CDCl_3_) δ 6.80 (s, 1H), 5.88 (s, 1H), 4.88 (s, 2H), 2.87 (q, *J* = 17.9 Hz, 2H), 2.34 – 2.20 (m, 1H), 1.91 – 1.69 (m, 2H), 1.66 – 1.39 (m, 6H), 1.32 – 1.13 (m, 3H), 1.10 – 0.95 (m, 2H), 0.94 – 0.86 (m, 1H), 0.83 (t, *J* = 7.3 Hz, 3H). HRMS (ESI), *m*/*z* calcd for C_20_H_25_Cl_2_O_4_ [M + H]^+^ 399.1130, found 399.1132.

*4-((2-butyl-6,7-dichloro-2-cyclopentyl-1-oxo-2,3-dihydro-1H-inden-5-yl)oxy)butanoic acid* (**8**) (**SN-401**): To a stirring solution of **6** (100 mg, 0.29 mmol, 1 equiv.) in anhydrous dimethylformamide (1 ml) was added potassium carbonate (45 mg, 0.32 mmol, 1.1 equiv.) and ethyl 4-bromobutyrate (46 ml, 0.32 mmol, 1.1 equiv.) and the reaction was allowed to stir at 60 ℃ for 2 hours. Once complete, to the reaction was added 4 N NaOH (1 ml) and the reaction was allowed to stir at 100 ℃ for 60 minutes. Once complete, reaction was concentrated and purified by column chromatography using 0-10% methanol in dichloromethane as eluent to provide **SN-401** as a clear solid (111 mg, 89%). ^1^H NMR (300 MHz, CDCl_3_) δ 10.77 (s, 1H), 6.86 (s, 1H), 4.21 (t, *J* = 5.9 Hz, 2H), 2.88 (t, *J* = 14.4 Hz, 2H), 2.69 (t, *J* = 7.0 Hz, 2H), 2.26 (dd, *J* = 12.6, 6.1 Hz, 3H), 1.87 – 1.73 (m, 2H), 1.64 – 1.44 (m, 6H), 1.35 – 1.10 (m, 4H), 1.08 – 0.95 (m, *J* = 15.0, 7.7 Hz, 2H), 0.82 (t, *J* = 7.3 Hz, 3H). HRMS (ESI), *m*/*z* calcd for C_22_H_29_Cl_2_O_4_ [M + H]^+^ 427.1443, found 427.1446.

*5-((2-butyl-6,7-dichloro-2-cyclopentyl-1-oxo-2,3-dihydro-1H-inden-5-yl)oxy)pentanoic acid* (**9**) (**SN-403**): To a stirring solution of **6** (100 mg, 0.29 mmol, 1 equiv.) in anhydrous dimethylformamide (1 ml) was added potassium carbonate (45 mg, 0.32 mmol, 1.1 equiv.) and ethyl 6-bromovalerate (51 ml, 0.32 mmol, 1.1 equiv.) and the reaction was allowed to stir at 60 ℃ for 2 hours. Once complete, to the reaction was added 4 N NaOH (1 ml) and the reaction was allowed to stir at 100 ℃ for 60 minutes. Once complete, reaction was concentrated and purified by column chromatography using 0-10% methanol in dichloromethane as eluent to provide **SN-403** as a clear solid (114 mg, 88%). ^1^H NMR (300 MHz, CDCl_3_) δ 10.95 (s, 1H), 6.85 (brs, 1H), 4.16 (t, *J* = 5.7 Hz, 2H), 2.96 – 2.75 (m, 2H), 2.61 – 2.44 (m, 2H), 2.35 – 2.17 (m, 1H), 2.10 – 1.87 (m, 4H), 1.86 – 1.70 (m, 2H), 1.66 – 1.38 (m, 6H), 1.32 – 1.13 (m, 3H), 1.08 – 0.96 (m, 2H), 0.94 – 0.86 (m, 1H), 0.86 – 0.73 (m, 3H). HRMS (ESI), *m*/*z* calcd for C_23_H_31_Cl_2_O_4_ [M + H]^+^ 441.1599, found 441.1601.

*6-((2-butyl-6,7-dichloro-2-cyclopentyl-1-oxo-2,3-dihydro-1H-inden-5-yl)oxy)hexanoic acid* (**10**) (**SN-406**): To a stirring solution of **6** (100 mg, 0.29 mmol, 1 equiv.) in anhydrous dimethylformamide (1 ml) was added potassium carbonate (45 mg, 0.32 mmol, 1.1 equiv.) and ethyl 6-bromohexanoate (58 ml, 0.32 mmol, 1.1 equiv.) and the reaction was allowed to stir at 60 ℃ for 2 hours. Once complete, to the reaction was added 4 N NaOH (1 ml) and the reaction was allowed to stir at 100 ℃ for 60 minutes. Once complete, reaction was concentrated and purified by column chromatography using 0-10% methanol in dichloromethane as eluent to provide **SN-406** as a clear solid (115 mg, 86%). ^1^H NMR (300 MHz, CDCl_3_) δ 11.70 (s, 1H), 6.85 (s, 1H), 4.13 (t, *J* = 6.2 Hz, 2H), 2.93 – 2.74 (m, 2H), 2.43 (t, *J* = 7.3 Hz, 2H), 2.32 – 2.17 (m, 1H), 1.98 – 1.87 (m, 2H), 1.85 – 1.68 (m, 4H), 1.66 – 1.40 (m, 8H), 1.28 – 1.12 (m, 3H), 1.07 – 0.93 (m, 2H), 0.91 – 0.70 (m, 4H). HRMS (ESI), *m*/*z* calcd for C_24_H_33_Cl_2_O_4_ [M + H]^+^ 455.1756, found 455.1756.

*7-((2-butyl-6,7-dichloro-2-cyclopentyl-1-oxo-2,3-dihydro-1H-inden-5-yl)oxy)heptanoic acid* (**11**) (**SN-407**): To a stirring solution of **6** (100 mg, 0.29 mmol, 1 equiv.) in anhydrous dimethylformamide (1 ml) was added potassium carbonate (45 mg, 0.32 mmol, 1.1 equiv.) and ethyl 7-bromoheptanoate (63 ml, 0.32 mmol, 1.1 equiv.) and the reaction was allowed to stir at 60 ℃ for 2 hours. Once complete, to the reaction was added 4 N NaOH (1 ml) and the reaction was allowed to stir at 100 ℃ for 60 minutes. Once complete, reaction was concentrated and purified by column chromatography using 0-10% methanol in dichloromethane as eluent to provide **SN-407** as a clear solid (122 mg, 89%). ^1^H NMR (300 MHz, CDCl_3_) δ 11.52 (s, 1H), 6.85 (s, 1H), 4.12 (t, *J* = 6.3 Hz, 2H), 2.84 (q, *J* = 18.2 Hz, 2H), 2.47 – 2.32 (m, 2H), 2.32 – 2.18 (m, 1H), 1.96 – 1.84 (m, 2H), 1.83 – 1.64 (m, 4H), 1.62 – 1.39 (m, 10H), 1.28 – 1.14 (m, 3H), 1.08 – 0.94 (m, 2H), 0.91 (d, *J* = 8.5 Hz, 1H), 0.81 (t, *J* = 7.3 Hz, 3H). HRMS (ESI), *m*/*z* calcd for C_25_H_35_Cl_2_O_4_ [M + H]^+^ 469.1912, found 469.1896.

**Scheme 2:**
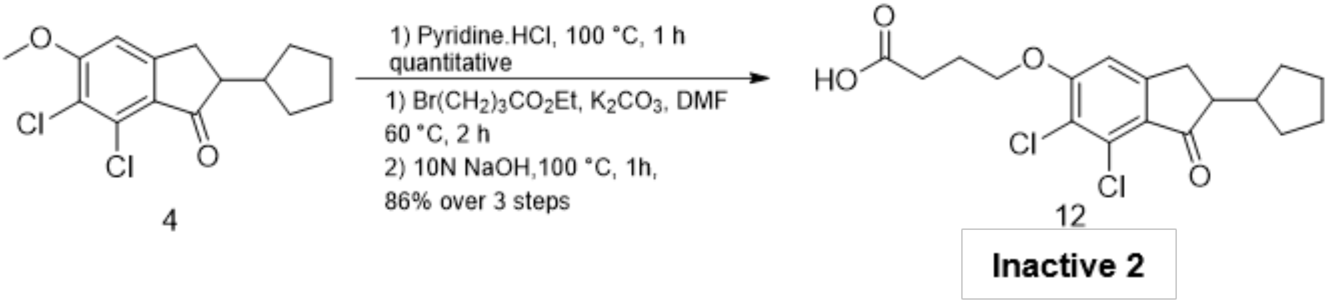

*4-((6,7-dichloro-2-cyclopentyl-1-oxo-2,3-dihydro-1H-inden-5-yl)oxy)butanoic acid* (**12**) (Inactive 2): To **4** (100 mg, 0.36 mmol, 1 equiv.) was added aluminum chloride (89 mg, 0.67 mmol, 2 equiv.) and sodium iodide (101 mg, 0.67 mmol, 2 equiv.) and the resulting solid mixture was triturated and allowed to stir at 70 ℃ for 60 minutes. Once complete, the reaction was diluted with dichloromethane and washed with aqueous saturated sodium thiosulfate solution. The organic fractions were collected and concentrated to give a beige solid which was then washed multiple times with hexanes to provide phenol intermediate as white solid which was used for the next step. To a stirring solution of the product form the first step in anhydrous dimethylformamide (1 ml) was added potassium carbonate (53 mg, 0.39 mmol, 1.1 equiv.) and ethyl 4-bromobutyrate (55 ml, 0.39 mmol, 1.1 equiv.) and the reaction was allowed to stir at 60 ℃ for 2 hours. Once complete, to the reaction was added 4 N NaOH (1 ml) and the reaction was allowed to stir at 100 ℃ for 60 minutes. Once complete, reaction was concentrated and purified by column chromatography using 0-10% methanol in dichloromethane as eluent to provide Inactive 2 as a clear solid (107 mg, 86%). ^1^H NMR (300 MHz, CDCl_3_) δ 6.87 (s, 1H), 4.21 (t, *J* = 5.9 Hz, 2H), 3.26 – 3.02 (m, 1H), 2.94 – 2.56 (m, 4H), 2.40 – 2.19 (m, 3H), 2.03 – 1.90 (m, 1H), 1.74 – 1.50 (m, 5H), 1.47 – 1.32 (m, 1H), 1.19 – 1.00 (m, 1H). HRMS (ESI), *m*/*z* calcd for C_18_H_21_Cl_2_O_4_ [M + H]^+^ 371.0817, found 371.0808.

**Scheme 3:**
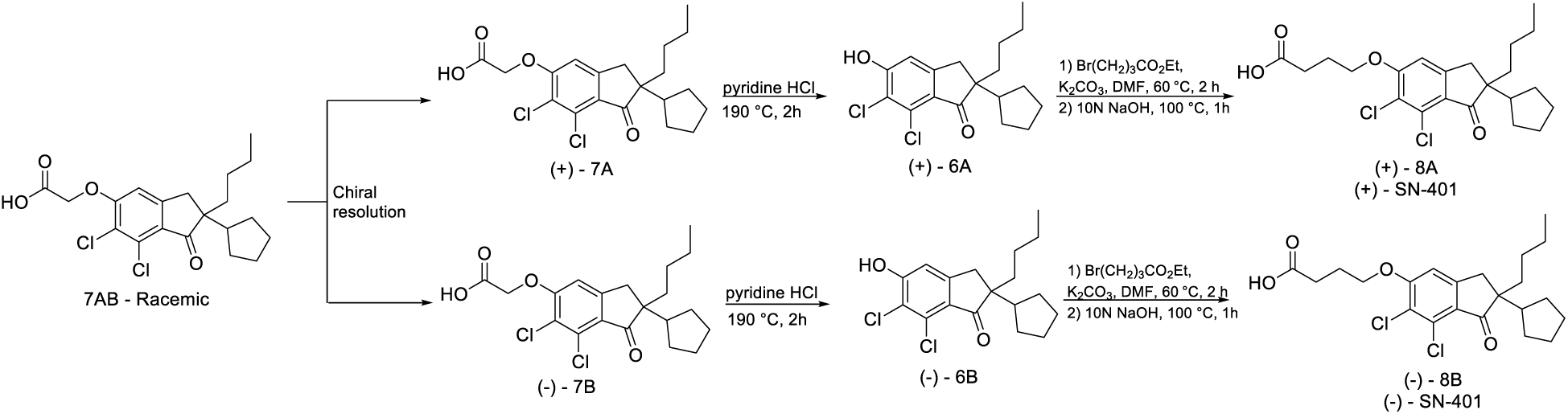

Enantiomerically enriched **SN-401** isomers were synthesized following literature reported procedure and as depicted in scheme 3^1^. In brief, racemic compound **7** (1 equiv.) was dissolved along with cinchonine (1 equiv.) in minimum amount of hot DMF and the allowed to cool. The precipitated salt was separated (filtrate used to obtain opposite enantiomer) and recrystallized 5 additional times from DMF, followed by acidification of salt with aqueous HCl and extraction into ether. The ether was evaporated under vacuum to furnish the enantiomerically enriched **(+) – 7A** in 23% yield; [a]^25^_D_ +16.8° (c 5, EtOH). The DMF filtrate from the first step now enriched in **(-) – 7B** was concentrated and acidified with aqueous HCl and extracted in ether and concentrated to give solid. This resulting solid (1 equiv.) was dissolved with cinchonidine (1 equiv.) in minimum amount of hot ethanol and then allowed to cool. The precipitated salt was separated and recrystallized 5 additional times from DMF, followed by acidification of salt with aqueous HCl and extraction into ether. The ether was evaporated under vacuum to furnish enantiomerically enriched **(-) – 7A** in 19% yield; [a]^25^_D_ -15.6° (c 5, EtOH). The enantiomerically enriched **7A** and **7B** were then subjected to same two step reaction sequence involving transformation to respective phenols **(+) - 6A** and **(-) – 6B** followed by conversion to desired enantiomerically enriched oxybutyric acids **(+) - 8A** [a]^25^_D_ +15.9° (c 5, EtOH) and **(-) – 8B** [a]^25^_D_ -14.5° (c 5, EtOH). The ^1^H NMR and HRMS for enantiomerically enriched products are same as racemic compounds and thus not reported.

### Electrophysiology

Patch-clamp recordings of β-cells, 3T3-F442A adipocytes and mature human adipocytes were performed as described previously^24, 27^. 3T3-F442A WT and KO preadipocytes were prepared as described in the Cell culture section above. For SWELL1 overexpression recordings, preadipocytes were first transduced with Ad5-CAG-LoxP-stop-LoxP-3XFlag-SWELL1 (MOI 12) in 2% FBS culture medium for two days and then overexpression induced by adding Ad5-CMV-Cre-eGFP (MOI 10-12) in 2% FBS culture medium for two more days and changed to 10% FBS containing culture media and were selected based on GFP expression (∼2-3 days). For β-cell recordings, islets were transduced with Ad-RIP2-GFP and then dispersed after 48-72 hours for patch-clamp experiments. GFP+ cells marked β-cells selected for patch-clamp recordings. Non-T2D islets were isolated from mice on regular chow diet between 8-13 weeks of age. Of these mice, 4 had an average body weight of 28.6 ± 0.51 g and blood glucose level of 148 ± 6.49 mg/dl respectively. T2D islets were obtained from mice fed with HFD for 4-5 months and their average body weight and glucose levels were 52.7 ± 3.0 g and 229 ± 21.4 mg/dl, respectively. For measuring I_Cl,SWELL_ inhibition by SN-401 congeners after activation of I_Cl,SWELL_, HEK-293 cells were perfused with hypotonic solution (Hypo, 210 mOsm) described below and then SN-401 congeners + Hypo applied at 10 and 7 µM to assess for % I_Cl,SWELL_ inhibition. To assess for I_Cl,SWELL_ inhibition upon application of SN-401 congeners to the closed SWELL1-LRRC8 channel, HEK-293 cells were pre-incubated with vehicle (or SN-401, SN-406, Inactive 1 and Inactive 2) for 30 mins prior to hypotonic stimulation and then stimulated with hypotonic solution + SN-401 congeners. Recordings were measured using Axopatch 200B amplifier paired to a Digidata 1550 digitizer using pClamp 10.4 software. The extracellular buffer composition for hypotonic stimulation contains 90 mM NaCl, 2 mM CsCl, 1 mM MgCl_2_, 1 mM CaCl_2_, 10 mM HEPES, 10 mM Mannitol, pH 7.4 with NaOH (210 mOsm/kg). The extracellular isotonic buffer composition is same as above, but with mannitol concentration at 110 mM to raise the osmolarity to 300 mOsm/kg. The composition of intracellular buffer is 120 mM L-aspartic acid, 20 mM CsCl, 1 mM MgCl_2_, 5 mM EGTA, 10 mM HEPES, 5 mM MgATP, 120 mM CsOH, 0.1 mM GTP, pH 7.2 with CsOH. All recordings were carried out at room temperature (RT) with HEK-293 cells, β-cells and 3T3-F442A cells performed in whole-cell configuration and human adipocytes in perforated-patch configuration, as previously ^24, 27^.

### Western blot

Adipocytes were washed twice in ice-cold phosphate buffer saline and lysed in RIPA buffer (150 mM Nacl, 20 mM HEPES, 1% NP-40, 5 mM EDTA, pH 7.4) with proteinase/phosphatase inhibitors (Roche). The cell lysate was further sonicated in 10 sec cycle intervals for 2-3 times and centrifuged at 14000 rpm for 30 min at 4°C and repeated one more time to remove the excess fat. The supernatant was collected and further estimated for protein concentration using DC protein assay kit (Bio-Rad). Fat tissues were homogenized and suspended in RIPA buffer with inhibitors and further processed in similar fashion as described above. HUVECs were also prepared in a similarly except the lysates were spun only once to remove the cells debris and obtain the clear supernatant. Human islets were washed twice with phosphate buffer saline (PBS) and lysed using RIPA buffer containing protease and phosphatase inhibitors. The lysate was further clarified by freezing in liquid nitrogen for about 10 s for three cycles. The supernatant was collected from the whole lysate centrifuged at 12,000 rpm for 20 min at 4 °C. Protein samples were further prepared by boiling in 4X laemmli buffer. Approximately 10-50 µg of total protein was loaded in 4-15% gradient gel (Bio-Rad) for separation and protein transfer was carried out onto the PVDF membranes (Bio-Rad). Membranes were blocked in 5% BSA (or 5% milk for SWELL1) in TBST buffer (0.2 M Tris, 1.37 M NaCl, 0.2% Tween-20, pH 7.4) for 1 h and incubated with appropriate primary antibodies (5% BSA or milk) overnight at 4°C. The membranes were further washed in TBST buffer before adding secondary antibody (Bio-Rad, Goat-anti-rabbit, #170-6515) in 1% BSA (or 1% milk for SWELL1) in TBST buffer for 1 h at RT. The signals were developed by chemiluminescence (Pierce) and visualized using a Chemidoc imaging system (Biorad). The images were further analyzed for band intensities using ImageJ software. Following primary antibodies were used: anti-phospho-AKT2 (#8599s), anti-AKT2 (#3063s), anti-AKT (#4685), anti-phospho-AS160 (#4288s), anti-AS160 (#2670s), anti-GAPDH (#D16H11), p-eNOS (#9571), Total eNOS (#32027) and anti-β-actin (#8457s) from Cell Signaling; anti-SWELL1 as previously described^2^.

### Immunofluorescence

3T3-F442A preadipocytes (WT, KO) and differentiated adipocytes without or with SWELL1 overexpression (WT+SWELL1 O/E, KO+SWELL1 O/E) were prepared as described in the Cell culture section on collagen coated coverslips. In the case of SWELL1 membrane trafficking, the 3T3-F442A preadipocytes were incubated in the presence of vehicle (or SN-401, SN-406 and Inactive 1) at either 1 or 10 µM for 48h and further processed. The cells were fixed in ice-cold acetone for 15 min at -20°C and then washed four times with 1X PBS and permeabilized with 0.1% Triton X-100 in 1X PBS for 5 min at RT and subsequently blocked with 5% normal goat serum for 1 h at RT. Either anti-SWELL1 (1:400) or anti-Flag (1:1500, Sigma #F3165) antibody were added to the cells and incubated overnight at 4°C. The cells were then washed three times (1X PBS) prior and after addition of 1:1000 Alexa Fluor 488/568 secondary antibody (anti-rabbit, #A11034 or anti-mouse, #A11004) for 1 h at RT. Cells were counterstained with nuclear TO-PRO-3 (Life Technologies, #T3605) or DAPI (Invitrogen, #D1306) staining (1 µM) for 20 min followed by three washes with 1X PBS. Coverslips were further mounted on slides with ProLong Diamond anti-fading media. All images were captured using Zeiss LSM700/LSM510 confocal microscope with 63X objective (NA 1.4). SWELL1 membrane localization was quantified by stacking all the z-images and converting it into a binary image where the cytoplasmic intensity per unit area was subtracted from the total cell intensity per unit area using ImageJ software.

### Metabolic phenotyping

Mice were fasted for 6 h prior to glucose tolerance tests (GTT). Baseline glucose levels at 0 min timepoint (fasting glucose, FG) were measured from blood sample collected from tail snipping using glucometer (Bayer Healthcare LLC). Either 1 g or 0.75 g D-Glucose/kg body weight were injected (i.p.) for lean or HFD mice, respectively and glucose levels were measured at 7, 15, 30, 60, 90 and 120 min timepoints after injection. For insulin tolerance tests (ITTs), the mice were fasted for 4 h. Similar to GTTs, the baseline blood glucose levels were measured at 0 min timepoint and 15, 30, 60, 90 and 120 min timepoints post-injection (i.p.) of insulin (HumulinR, 1U/kg body weight for lean mice or 1.25 U/kg body weight for HFD mice). GTTs or ITTs with vehicle (or SN-401, SN-403, SN-406, SN-407 and Inactive 1) treated groups were performed approximately 24 hours after the last injection. For measuring serum insulin levels, the vehicle (or SN-401, SN-406 and Inactive 1) treated HFD mice were fasted for 6 h and injected (i.p.) with 0.75 g D-Glucose/kg body weight and blood samples collected at 0, 7, 15 and 30 min time points in microvette capillary tubes (SARSTEDT, #16.444) and centrifuged at 2000Xg for 20 min at 4°C. The collected plasma was then measured for insulin content by using Ultra-Sensitive Mouse Insulin ELISA Kit (Crystal Chem, #90080). All mouse studies were performed in a blinded fashion. Body weights for all the mice are listed in **Supplementary Table. S5**.

### Murine islet isolation and perifusion assay

For patch-clamp studies involving primary mouse β-cells, mice were anesthesized by injecting Avertin (0.0125 g/ml in H_2_O) followed by cervical dislocation. HFD or polygenic KKAy mice treated with either vehicle or SN-401, SN-406, SN-407 and Inactive 1 were anesthesized with 1-4% isoflurane followed by cervical dislocation. Islets were further isolated as described previously^27^. Human islets were cultured in RPMI media with 2% FBS overnight. The next day either scramble or shSWELL1 adenoviral transduction was carried out (final concentration of 5 × 10^7 PFU/ml) and the islets were incubated for 12 h. The islets were then washed with PBS three times and cultured in RPMI medium with 10% FBS for 4–5 days. For SN-401/Palmitate experiments, human islets were either transduced with adenoviral short hairpin for control (shScramble) or SWELL1 knockdown (shSWELL1) and murine islets isolated from floxed-SWELL1 mouse (SWELL1^fl/fl^) were either transduced with adenoviral control (Ad-RIP1-GCaMP6s) or Cre-recombinase (Ad-RIP1-GCaMP6s-2A-Cre) virus for 12 hours respectively in 2% FBS containing RPMI media. The islets were then washed in 1XPBS for three times and treated with either vehicle or SN-401 (10 µM) for 96 hours followed by treatment with 1:3 palmitate:BSA with or without SN-401 in 10% FBS containing RPMI media for 16 h (**Fig. 7c**). The GSIS perifusion assay for islets were performed using a PERI4-02 from Biorep Technologies. For each experiment, around 50 freshly isolated islets (all from the same isolation batch) were handpicked to match size of islets across the samples and loaded into the polycarbonate perifusion chamber between two layers of polyacrylamide-microbead slurry (Bio-Gel P-4, BioRad) by the same experienced operator. Perifusion buffer contained (in mM): 120 NaCl, 24 NaHCO_3_, 4.8 KCl, 2.5 CaCl_2_, 1.2 MgSO_4_, 10 HEPES, 2.8 glucose, 27.2 mannitol, 0.25% w/v bovine serum albumin, pH 7.4 with NaOH (300 mOsm/kg). Perifusion buffer kept at 37°C was circulated at 120 μl/min. After 48 min of washing with 2.8 mM glucose solution for stabilization, islets were stimulated with the following sequence: 16 min of 16.7 mM glucose, 40 min of 2.8 mM glucose, 10 min of 30 mM KCl, and 12 min of 2.8 mM glucose. Osmolarity was matched by adjusting mannitol concentration when preparing solution containing 16.7 mM glucose. Serial samples were collected either every 1 or 2 min into 96 wells kept at 4°C. Insulin concentrations were further determined using commercially available ELISA kit (Mercodia). The area under the curve (AUC) for the high-glucose induced insulin release was calculated for time points between 50 to 74/84 min. At the completion of the experiments, islets were further lysed by addition of RIPA buffer and the amount of insulin was detected by ELISA.

### Drug pharmacokinetics

The pharmacokinetic studies for SN-401 and SN-406 were performed at Charles River Laboratory as outlined below. Male C57/BL6 mice were used in the study and assessed for a single dose (5 mg/kg) administration. The compounds were prepared in Cremaphor for i.p. and p.o dose routes and in 5% ethanol, 10% Tween-20 and water mix for i.v. route at a final concentration of 1 mg/mL. Terminal blood samples were collected via cardiac venipuncture under anesthesia at time points 0.08, 0.5, 2, 8 h post dose for i.v and at timepoints 0.25, 2, 8, 24 h post dose for i.p. and p.o. groups respectively with a sample size of 3 mice per timepoint. The blood samples were collected in tubes with K2 EDTA anticoagulant and further processed to collect plasma by centrifugation at 3500 rpm at 5 °C for 10 min. Samples were further processed in LC/MS to determine the concentration of the compounds. Non-compartmental analysis was performed to obtain the PK parameters using the PKPlus software package (Simulation Plus). The area under the plasma concentration-time curve (AUC_inf_) is calculated from time 0 to infinity where the C_max_ is the maximal concentration achieved in plasma and t_1/2_ is the terminal elimination half-life. Oral bioavailability was calculated as AUC_Oral_/AUC_IV_*100.

### Hyperinsulinemic euglycemic glucose clamps

Sterile silicone catheters (Dow-Corning) were placed into the jugular vein of mice under isoflurane anesthesia. Placed catheter was flushed with 200 U/mL heparin in saline and the free end of the catheter was directed subcutaneously via a blunted 14-gauge sterile needle and connected to a small tubing device that exited through the back of the animal. Mice were allowed to recover from surgery for 3 days, then received IP injections of vehicle or SN-401 (5 mg/kg) for 4 days. Hyperinsulinemic euglycemic clamps were performed on day 8 post-surgery on unrestrained, conscious mice as described elsewhere^79, 80^, with some modifications. Mice were fasted for 6 h at which time insulin and glucose infusion were initiated (time 0). At 80 min prior to time 0 basal sampling was conducted, where whole-body glucose flux was traced by infusion of 0.05 μCi/min D-[3-^3^H]-glucose (Perkin Elmer), after a priming 5 μCi bolus for 1 minute. After the basal period, starting at time 0 D-[3-^3^H]-glucose was continuously infused at the 0.2 μCi/min rate and the infusion of insulin (Humulin, Eli Lilly) was initiated with a bolus of 80 mU/kg/min (10 µl volume for 1 min) then followed by continuous infusion of insulin at the dose of 8 mU/kg/min throughout the assay. Fifty percent dextrose (Hospira) was infused at a variable rates (GIR) starting at the same time as the initiation of insulin infusion to maintain euglycemia at the targeted level of 150 mg/dL (8.3 mM). Blood glucose (BG) measurements were taken every ten minutes via tail vein sampling using Contour glucometer (Bayer). After mouse reached stable BG and GIR (typically, after 75 minutes since starting the insulin infusion; for some mice, a longer time was required to achieve steady state) a single bolus of 12 μCi of [1-^14^C]-2-deoxy-D-glucose (Perkin Elmer) in 96 μl of saline was administered. Plasma samples (collected from centrifuged blood) for determination of tracers enrichment, glucose level and insulin concentration were obtained at times -80, -20, -10, 0, and every 10 min starting at 80 min post-insulin (5 min. after [1-^14^C]-2-deoxy-D-glucose bolus was administered) until the conclusion of the assay at 140 min. Tissue samples were then collected from mice under isofluorane anesthesia from organs of interest (e.g., liver, heart, kidney, white adipose tissue, brown adipose tissue, gastrocnemius, soleus etc.) for determination of 1-^14^C]-2-deoxy-D-glucose tracer uptake. Plasma and tissue samples were processed as described previously^79^. Briefly, plasma samples were deproteinized with Ba(OH)_2_ and ZnSO_4_ and dried to eliminate tritiated water. The glucose turnover rate (mg/kg-min) was calculated as the rate of tracer infusion (dpm/min) divided by the corrected plasma glucose specific activity (dpm/mg) per kg body weight of the mouse. Fluctuations from steady state were accounted for by use of Steele’s model. Plasma glucose was measured using Analox GMD9 system (Analox Technologies).

Tissue samples (∼30 mg each) were homogenized in 750 µl of 0.5% perchloric acid, neutralized with 10 M KOH and centrifuged. The supernatant was then used for first measuring the abundance of total [1-^14^C] signal (derived from both 1-^14^C -2-deoxy-D-glucose, 1-^14^C -2-deoxy-D-glucose 6 phosphate) and, following a precipitation step with 0.3 N Ba(OH)_3_ and 0.3 N ZnSO_4_, for the measuring of non-phosphorylated 1-^14^C -2-deoxy-D-glucose. Glycogen was isolated by ethanol precipitation from 30% KOH tissue lysates, as described^81^. Insulin level in plasma at T0 and T140 were measured using a Stellux ELISA rodent insulin kit (Alpco).

### Protein purification and nanodisc formation

SWELL1 was purified as described previously^30^. Freshly purified SWELL1 from gel filtration in Buffer 1 (20 mM HEPES, 150 mM KCl, 1 mM EDTA, 0.025% DDM, pH 7.4) was reconstituted into MSP1E3D1 with a lipid mix (2:1:1 weight ratio of DOPE:POPC:POPS lipids (Avanti, Alabaster, Alabama)) at a final molar ratio of 1:2.5:200 (Monomer Ratio: SWELL1, MSP1E3D1, Lipid Mix). First, solubilized lipid in Column Buffer (20 mM HEPES, 150 mM KCl, 1 mM EDTA pH 7.4) was mixed with additional DDM detergent, Column Buffer, and SWELL1. This solution was mixed at 4°C for 30 min before addition of purified MSP1E3D1. This addition brought the final concentrations to approximately 10 µM SWELL1, 25 µM MSP1E3D1, 2 mM lipid mix, and 3.3 mM DDM in Column Buffer (1 mL reaction). The solution with MSP1E3D1 was mixed at 4°C for 10 min before addition of 130 mg of Biobeads SM2 (Bio-Rad, Hercules, CA). Biobeads (washed into methanol, water, and then Column Buffer) were weighed with liquid removed by P1000 tip (Damp weight). This mix was incubated at 4°C for 30 min before addition of another 130 mg of Biobeads (final 260 mg of Biobeads per mL). This final mixture was then mixed at 4°C overnight (∼14 hr). Supernatant was cleared of beads by letting large beads settle and carefully removing supernatant with a pipet. Sample was spun for 5 min at 21,000 × g before loading onto a Superose 6 column in Column Buffer without EDTA. Peak fractions corresponding to SWELL1 in MSP1E3D1 were collected, 100 kDa cutoff spin concentrated, and then re-run on the Superose 6. The fractions corresponding to the center of the peak were then pooled and concentrated prior to grid preparation.

### Grid preparation

SN-407 in DMSO (Stock 10 mM) was added to SWELL1-Msp1E3D1 sample to give a final concentration of 1 mg/mL SWELL1-MSP1E3D1 and 100 µM SN-407. The drug was allowed to equilibrate and bind complex on ice for ∼1 hour prior to freezing grids. Sample with drug was cleared by a 5 min 21,000 × g spin prior to grid making. For freezing grids, a 2 µl drop of protein was applied to freshly glow discharged Holey Carbon, 300 mesh R 1.2/1.3 gold grids (Quantifoil, Großlöbichau, Germany). A Vitrobot Mark IV (Thermo Fisher Scientific, Waltham, MA) was utilized with 22°C, 100% humidity, one blot force, and a 3 s blot time, before plunge freezing in liquid ethane. Grids were then clipped in autoloader cartridges for collection.

### Data collection

SWELL1-MSP1E3D1 with SN-407 grids were transferred to a Talos Arctica cryo-electron microscope (Thermo Fisher Scientific) operated at an acceleration voltage of 200 kV. Images were recorded in an automated fashion with SerialEM^82^ using image shift with a target defocus range of −0.7 ∼ −2.2 µm over 5.493 s as 50 subframes with a K3 direct electron detector (Gatan, Pleasanton, CA) in super-resolution mode with a super-resolution pixel size of 0.5685 Å^83^. The electron dose was 9.392 e^-^ / Å / s (1.0318 e^-^/ Å2/frame) at the detector level and total accumulated dose was 51.59 e-/Å2.

### Cryo-EM data processing

A total of 3576 movie stacks were collected, motion-corrected and binned to 1.137 Å/pixel using MotionCor2 in RELION3.1^84^, and CTF-corrected using Ctffind 4.1.13^85^ (See **Supplemental Fig. 7**). Micrographs with a Ctffind reported resolution estimate worse than 5 Å were discarded. A particle set generated from manual picking and template-based autopicking in RELION3.1 was cleaned and processed to 60,803 particles representing diverse views of the SWELL1 particle. These particles were then used to train Topaz^86^ to pick a set of 936,282 particles. This set was cleaned with 2D classification and heterogeneous refinement in cryoSPARCv2. We then generated a refinement in cryoSPARCv2 and then RELION3.1 using C6 symmetry and utilized this map to perform Bayesian Polishing. Polished particles were then refined in RELION3.1 with C6 symmetry with a mask for the extracellular domain (ECD), transmembrane, and linker domains of SWELL1 to 3.05 Å. This map did not show clear evidence of drug density in the ECD, which we hypothesized could be due to a combination of partial drug occupancy and asymmetric drug density (off the symmetry axis of SWELL1). To test this hypothesis we performed symmetry expansion (in C6) followed by sequential 3D classification with C1 symmetry in RELION3.1 using an increasingly tightened mask on the ECD. We noted and selected classes with putative drug density in the ECD. We then used C1 refinement with C6 symmetry relaxation^48^ in RELION3.1 to further refine the density. These refinement angles were used for one additional 3D classification job to generate 2 classes: one with vertical density (85,831 particles) and one with tilted density (78,324 particles). The particles in each of these classes were then refined an additional time with symmetry relaxation and local angular sampling. Finally, these angles were used in a refinement in C1 with masking for the ECD, transmembrane, and linker domain to generate the final maps.

### Modeling and refinement

Maps from refinement for the vertical and tilted drug density were used for modeling in Coot ^87^. First, the model from ^30^ (PDB: 6NZW) was docked in the map density. As the lipid mix used in this study contained a majority of DOPE lipid, the lipid acyl chains between LRRC8A subunits were modeled as DOPE. The model and restraints for Smod7 were generated using Phenix.elbow^88^ and then SN-407 was placed and refined in the putative drug density for each map. Real space refinement of each model was carried out using Phenix.real_space_refine. Molprobity^89^ was used to evaluate the stereochemistry and geometry of the structure for subsequent rounds of manual adjustment in Coot and refinement in Phenix. Phenix.mtriage was then used map and model validation. Figures were prepared using Chimera^90^, ChimeraX^91^, Prism, and Adobe Photoshop and Illustrator software.

### Quantitative RT-PCR

3T3-F442A preadipocytes cells treated with either vehicle (DMSO) or 10 µM SN-401 for 96 h were solubilized in TRIzol and the total RNA was isolated using PureLink RNA kit (Life Technologies). For differentiated adipocytes, vehicle or 10 µM SN-401 were added during 7-11 days of differentiation for 96 h and then serum starved (+ DMSO/SN-401) for 6 h and stimulated with 0 and 10 nM insulin/serum containing media (+ DMSO or SN-40X) for 15 min and processed for RNA as described above. The cDNA synthesis, qRT-PCR reaction and quantification were carried out as described previously^24^.

### Liver isolation, triglycerides and histology

HFD mice treated with either vehicle or SN-401 were anesthetized with 1-4% isoflurane followed by cervical dislocation. Gross liver weights were measured and identical sections from right medial lobe of liver were dissected for further examinations. Total triglyceride content was determined by homogenizing 10-50 mg of tissue in 1.5 ml of chloroform:methanol (2:1 v/v) and centrifuged at 12000 rpm for 10 mins at 4°C. An aliquot, 20 ul, was evaporated in a 1.5 ml microcentrifuge tube for 30 mins. Triglyceride content was determined by adding 100 µl of Infinity Triglyceride Reagent (Fisher Scientific) to the dried sample followed by 30 min incubation at RT. The samples were then transferred to a 96 well plate along with standards (0-2000 mg/dl) and absorbance was measured at 540 nm and the final concentration was determined by normalizing to tissue weight. For histological examination, liver sections were fixed in 10% zinc formalin and paraffin embedded for sectioning. Hematoxylin and eosin (H&E) stained sections were then assessed for steatosis grade, lobular inflammation and hepatocyte ballooning for non-alcoholic fatty liver disease (NAFLD) scoring as described ^46, 92, 93^.

### Quantification and statistical analysis

Standard unpaired or paired two-tailed Student’s t-test were performed while comparing two groups. One-way Anova was used for multiple group comparison. For GTTs and ITTs, 2-way analysis of variance (Anova) was used. The threshold for significance was 0.05 for all statistical comparisons. *, ** and *** represent *p-*values of <0.05, <0.01 and <0.001, respectively. All data are presented as mean ±SEM. Details of statistical analyses are presented in the figure legends.

## Acknowledgements

We thank R. Sigmund, J. Galbraith and M. Knudson of the University of Iowa Tissue procurement Core facility (TPC) for services provided related to acquisition of human adipose specimens (NCI award number P30CA086862). We thank the Diabetes Research Core at the Washington University in St. Louis for triglyceride estimation DRC (NIH P30 DK 020579). We also thank the Fraternal Order of Eagles Diabetes Research Center at the University of Iowa in performing euglycemic hyperinsulinemic clamps, Aloysius Klingelhutz (University of Iowa) for human adipose tissue transport and Michael Wright (University of Iowa) for gifting HEK-293 cells. P.R.C. and C.A.K. acknowledge support of predoctoral fellowships from the University of Iowa Center for Biocatalysis and Bioprocessing affiliated with the NIH-sponsored Predoctoral Training Program in Biotechnology (T32 GM008365). This work was supported by the John L. & Carol E. Lach Chair in Drug Delivery Technology (RJK), and by grants from the NIH NIDDK 1R01DK106009 (R.S.), R01DK115791 (A.W.N.) and the Roy J. Carver Trust (R.S.).

## Author Contributions

Conceptualization, R.S.; methodology, S.K.G., L.X., C.K, A.K., P.C., D.M.K., E.E.G., C.A.K., W.J.G., R.D.S, E.B.T., C.M.T, S.G.B., Y.Z., R.S.; formal analysis, R.S., S.K.G., L.X., C.K., D.M.K, E.E.G., R.D.S., S.G.B., W.J.G., C.M.T., D.J.L.; investigation, R.S., S.K.G., L.X., C.K., P.C., R.K., W.J.G., C.M.T.; resources, J.K.S., I.S., P.N., Y.I., R.K. R. S.; writing (original draft), R.S., writing (review and editing), R.S., S.K.G., A.W.N., Y.I., D.M.K., S.G.B., W.J.G., L.X., P.C., R.K., D.J.L.; visualization, R.S., S.K.G., L.X., D.M.K., P.C., C.K.; supervision, R.S., A.W.N., S.G.B., Y.I., R.K.; funding acquisition, R.K., R.S.

## Data availability

All requests for resources and reagents should be addressed to and will be fulfilled upon reasonable request.

## Competing financial interests

R.S. is co-founder of Senseion Therapeutics, Inc., a start-up company developing SWELL1 modulators for human disease. D.J.L. is co-Founder and CEO of Senseion Therapeutics, Inc.

## Supplementary Figure Legends

**Supplementary Fig. S1.**
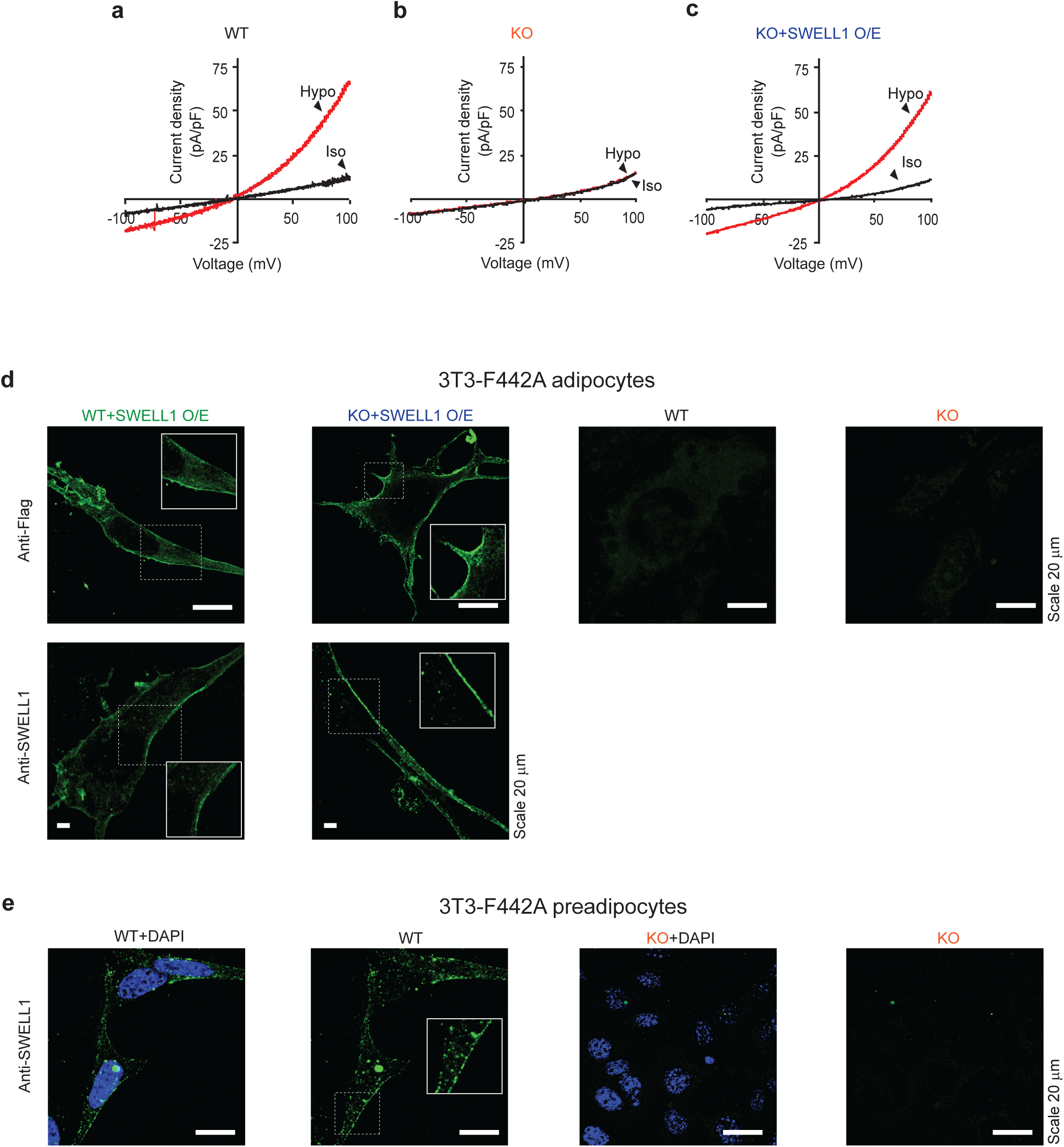
Transient expression of full-length SWELL1 with C-terminal 3XFlag tag rescues I_Cl,SWELL_ and traffics to the plasma membrane. **a-c.** Current-voltage plots of I_Cl,SWELL_ measured in 3T3-F442A preadipocytes WT (**a**), KO (**b**) and adenoviral overexpression of SWELL1 in KO (KO+SWELL1 O/E) (**c**) at baseline (iso, black trace) and hypotonic (hypo, red trace) stimulation respectively. **d**. Immunostaining images demonstrating localization of endogenous SWELL1 or overexpressed SWELL1 with anti-Flag or anti-SWELL1 antibody (Scale bar: 20 µm). **e**. Validation of SWELL1 antibody in WT 3T3-F442A compared to SWELL1 KO pre-adipocytes (Scale bar: 20 µm), revealing a punctate pattern of endogenous SWELL1 localization (inset).

**Supplementary Fig. S2.**
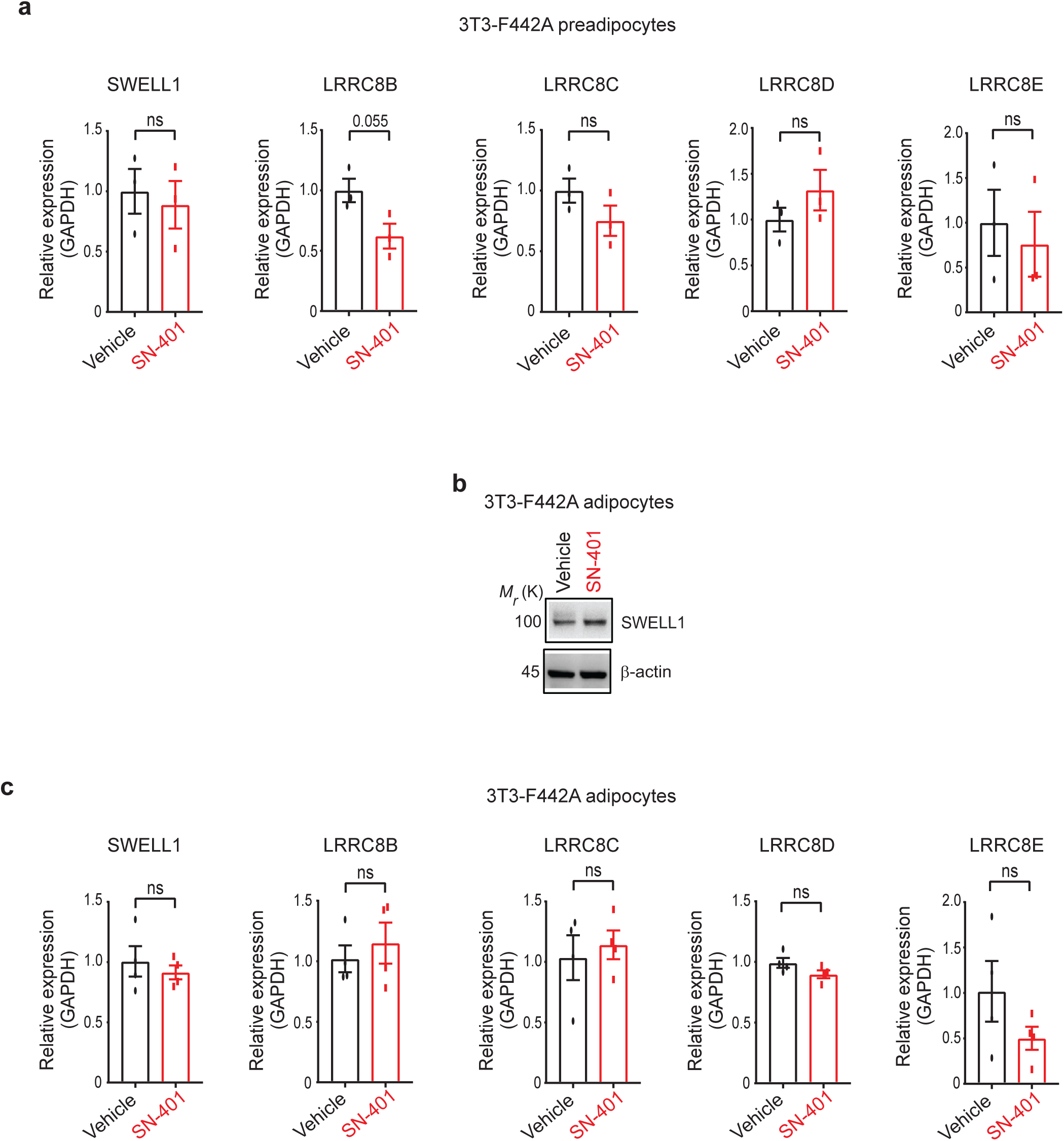
Increases in SWELL1 protein are not associated with increases in mRNA expression of SWELL1/LRRC8a or LRRC8b-e. **a.** Relative mRNA expression of LRRC8 family members relative to GAPDH assessed by qPCR (n = 3 each) for 3T3 F-442A preadipocytes treated with either vehicle or 10 µM SN-401 for 96 hours. **b**. Western blots showing increase in SWELL1 protein expression relative to β-actin in 3T3-F442A adipocytes treated with either vehicle or 10 µM SN-401 for 96 hours. **c**. Relative mRNA expression of LRRC8 family members relative to GAPDH assessed by qPCR (n = 4 each) from the same 3T3-F442A adipocytes treated with either vehicle or 10 µM SN-401 for 96 hours obtained in b. Data are represented as mean ±SEM. Two-tailed unpaired t-test was used in a and c where *, ** and *** represents *p<0.05*, *p<0.01* and *p<0.001* respectively.

**Supplementary Fig. S3.**
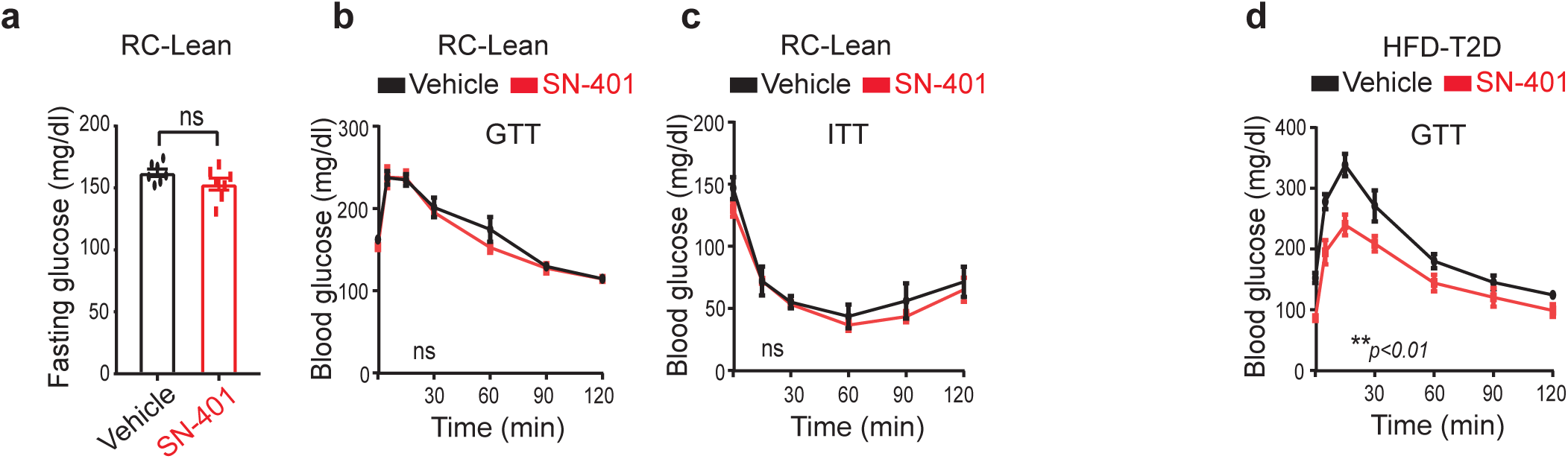
SN-401 activity in lean non-T2D mice and effects of chronic dosing in HFD-fed mice. Fasting glucose levels **(a)**, GTT **(b)** and ITT **(c)** of C57BL/6 lean mice on regular-chow diet treated with either vehicle or SN-401 (5 mg/kg i.p) for 10 days (n = 7 males in each group). d. GTT of HFD-T2D mice (8 weeks HFD) treated with either vehicle (n = 5 males) or SN-401 (5 mg/kg i.p, n = 4 males) for 8 weeks. Data are represented as mean ±SEM. Two-tailed unpaired t-test was used in a. ‘ns’ indicates the difference was not significant.

**Supplementary Fig. S4.**
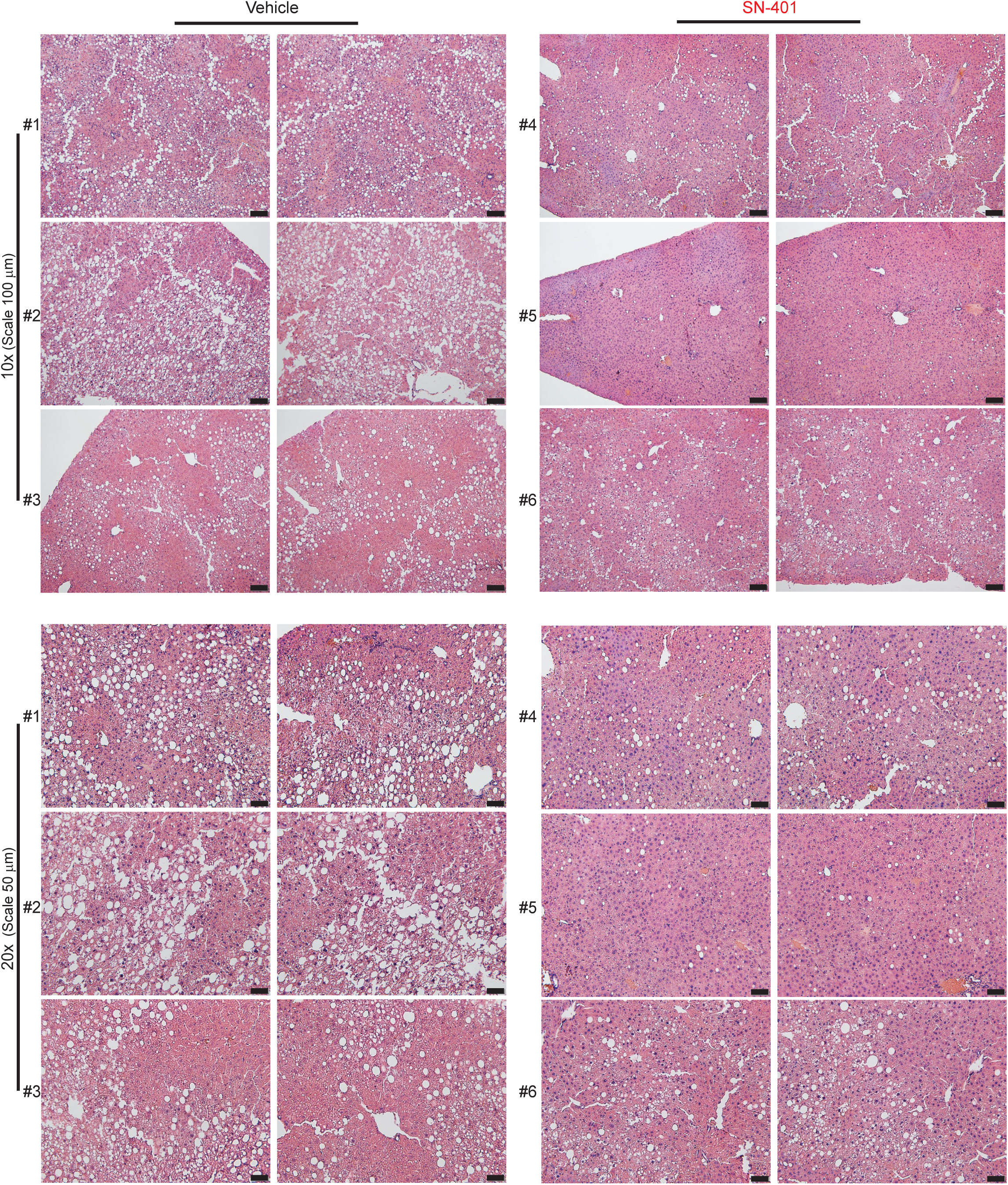
SN-401 improves non-alcoholic fatty liver disease in murine T2D models. Images of hematoxylin and eosin stained liver histology sections of HFD-T2D mice treated with either vehicle or SN-401 (5 mg/kg i.p) as in Fig. 4e. Scale: 10X, 100 µm and 20X, 50 µm).

**Supplementary Fig. S5.**
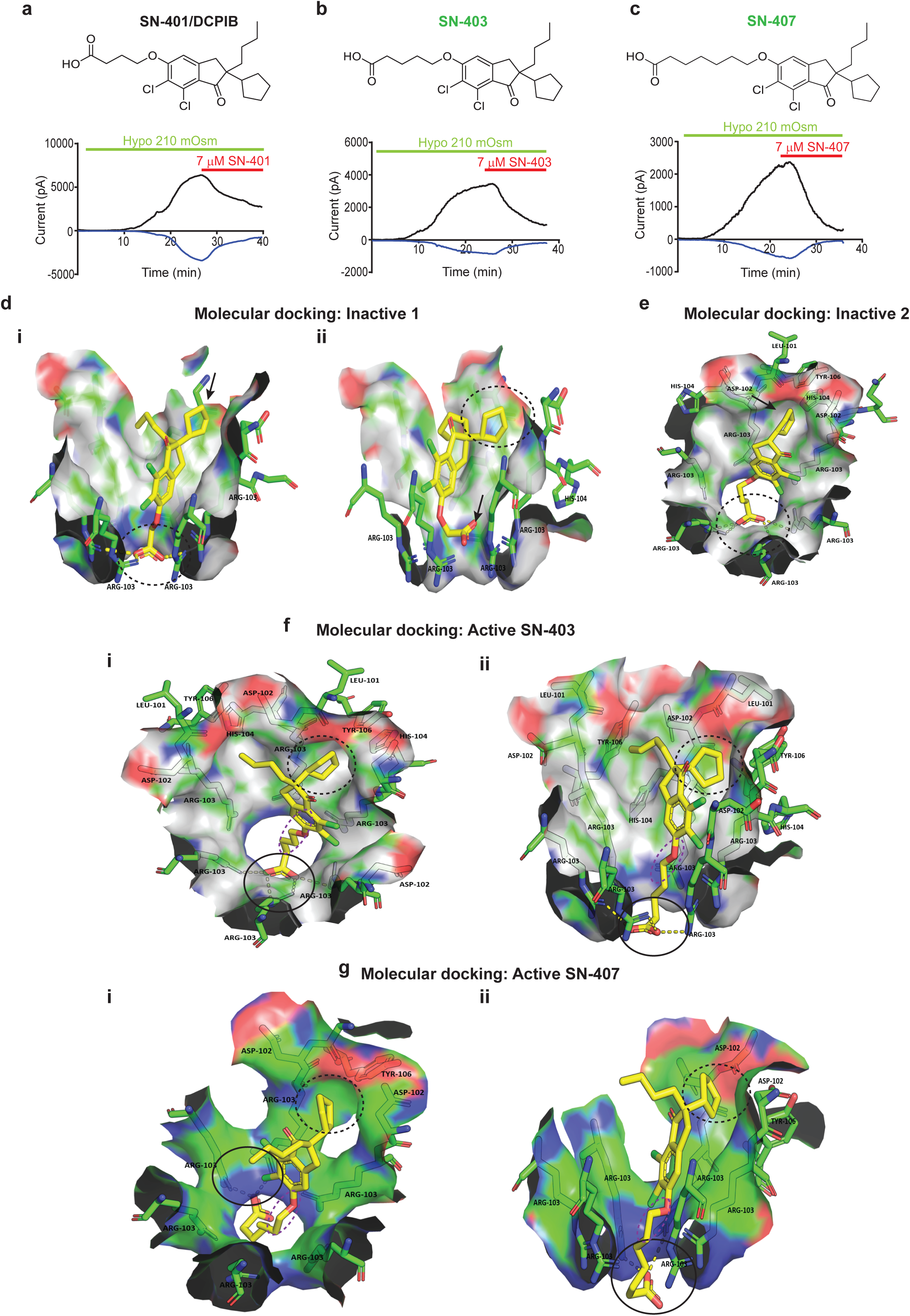
Defining the SN-401-SWELL1 structure-activity relationship by combining chemical synthesis, molecular docking simulations and patch-clamp electrophysiology. Chemical structures (top) of **a**. SN-401/DCPIB, (**b**) SN-403 and (c) SN-407 and I_Cl,SWELL_ inward and outward current over time (bottom) upon hypotonic (210 mOsm) stimulation and subsequent inhibition by 7 μM SN-401, SN-403 and SN-407 in HEK-293 cells. **d.** Binding poses for Inactive 1; (**i**) side view of first binding pose of Inactive 1 showing potential electrostatic interaction with R103 (dotted circle) but unable to reach into and occupy the hydrophobic cleft (black arrow); (ii) side view of second pose for Inactive 1 with the cyclopentyl group occupying the hydrophobic cleft (broken circle) but the carboxylate group unable to reach and interact with R103 (black arrow). **e**. Binding pose for Inactive 2 reveal that the carboxylate group can reach and electrostatically interact with R103 but in the absence of the butyl group cannot orient the cyclopentyl ring to occupy the hydrophobic cleft without introducing excessive structural strain on the carbon connecting the core with the cyclopentyl ring. **f-g**. (i) top and (ii) side view of binding poses of SN-403 (f) and SN-407 (**g**); the carboxylate groups interact with guanidine group of R103 residues (black circle), the cyclopentyl group occupies a shallow hydrophobic cleft at the interface of two monomers formed by D102 and L101 (black broken circle) and the alkyl side chain SN-403 or SN-407 interacts with the alkyl side chain of R103 (purple broken circle).

**Supplementary Fig. S6.**
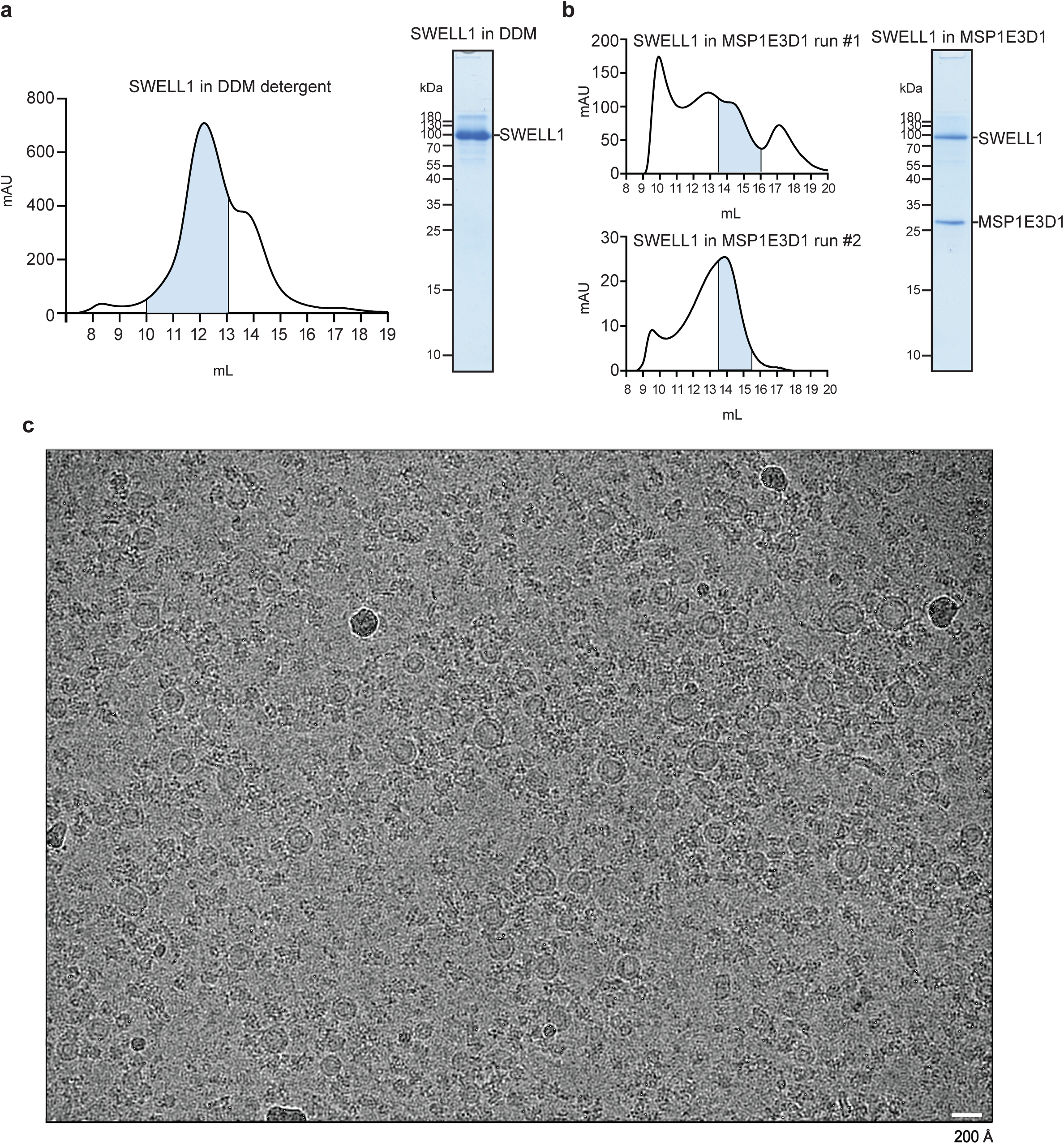
Purification, reconstitution, and cryo-EM imaging of SWELL1. **a.** Size exclusion chromatogram (Superose 6 Increase) of SWELL1 purified into DDM detergent (left). Pooled fractions corresponding to hexameric SWELL1 are highlighted in blue. Coomassie-stained SDS-PAGE of pooled SWELL1 homohexamer-containing fractions (right). **b.** Size exclusion chromatogram of SWELL1 reconstituted into MSP1E3D1 lipid nanodiscs (left-upper). Pooled fractions were then re-run (left-lower) and pooled fractions were concentrated for drug addition, grid freezing, and coomassie-stained SDS-PAGE (right). **c.** Example micrograph from SN-407-SWELL1 in MSP1E3D1 cryo-EM data collection.

**Supplementary Fig. S7.**
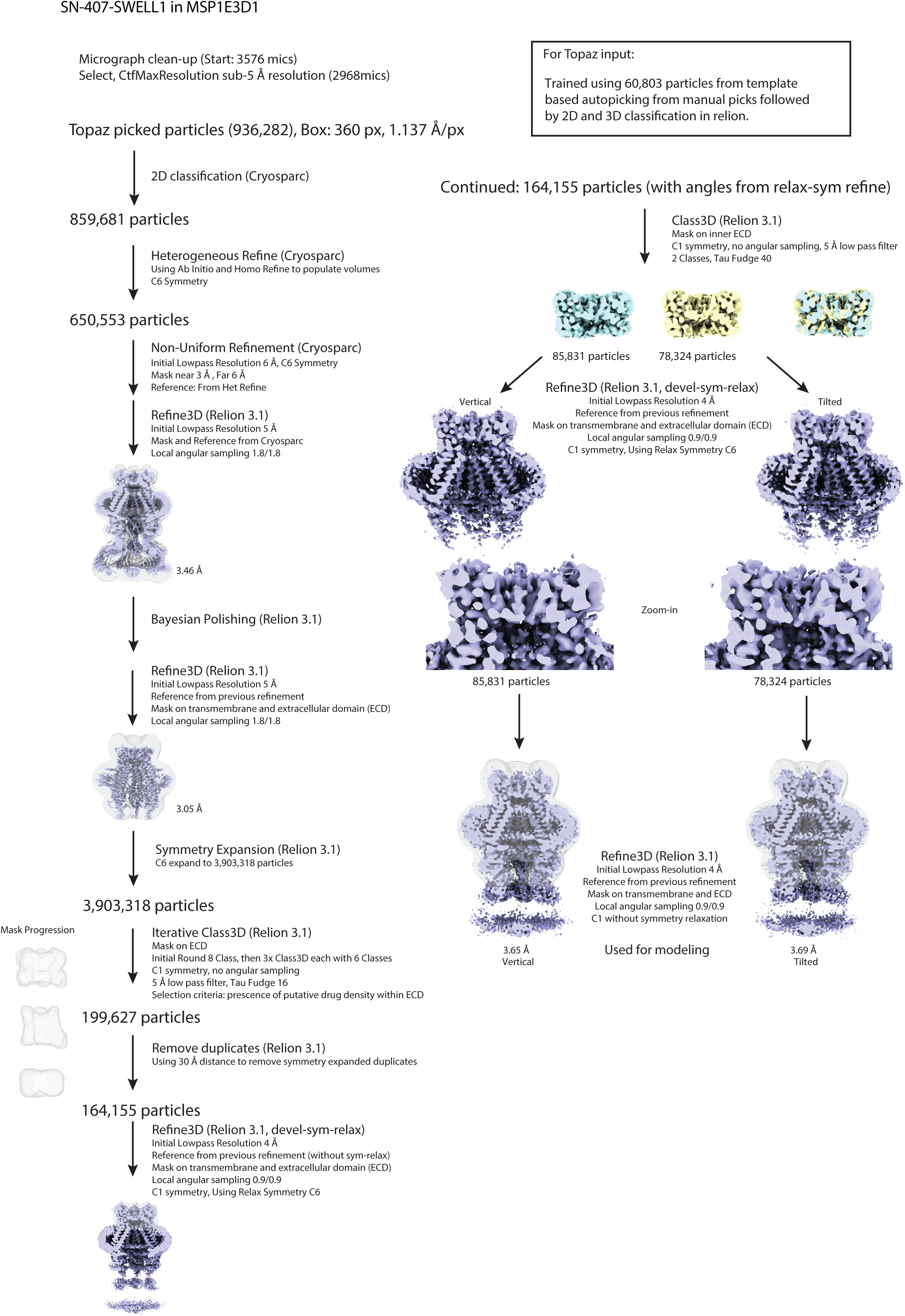
Cryo-EM processing pipeline for SN-407-SWELL1 in MSP1E3D1 lipid nanodiscs. Overview of Cryo-EM data processing pipeline in cryoSPARC and Relion. See Methods for additional details.

**Supplementary Fig. S8.**
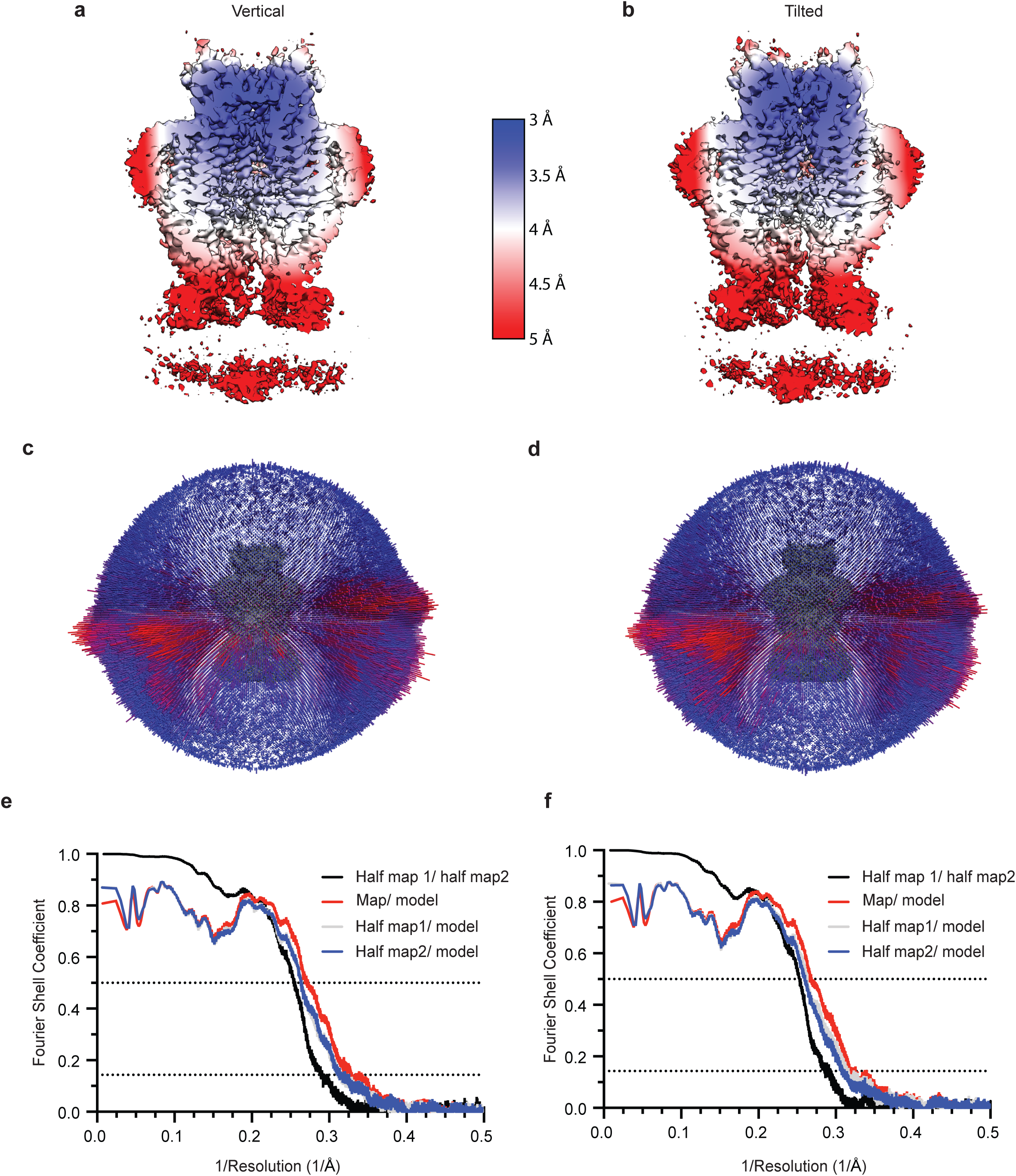
Cryo-EM validation for SN-407-SWELL1 in MSP1E3D1 lipid nanodiscs. **a-b.** Local resolution estimated in Relion colored as indicated on the final map for vertical (left) and tilted (right) drug density classes. **c-d.** Angular distribution of particles used in final refinement with maps for reference. **e-f**. Fourier Shell Correlation (FSC) relationships between (black) the two unfiltered half-maps from refinement and used for calculating overall resolution at 0.143, (red) the final map and model, (gray) half-map one and model, and (blue) half-map and model.

**Supplementary Fig. S9.**
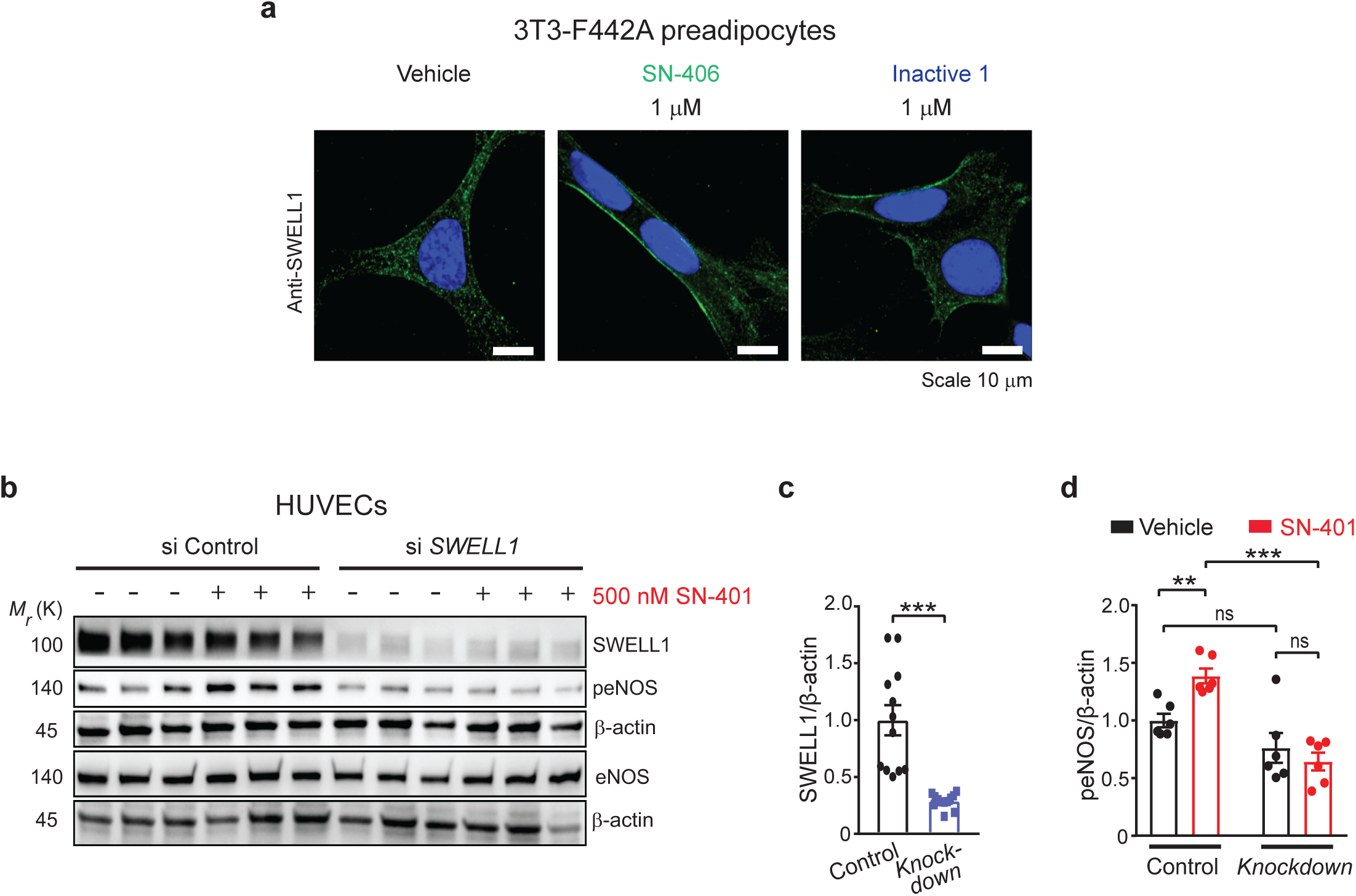
SN-406 increase SWELL1 plasma membrane localization while Inactive 1 does not and SN-401 mediated induction of peNOS activity in HUVECs is SWELL1 dependent. **a.** Representative immunostaining images demonstrating localization of endogenous SWELL1 in WT 3T3-F442A preadipocytes treated with vehicle or SN-406/Inactive 1 at 1 µM for 48h (Scale bar: 10 µm). **b.** Western blots detecting SWELL1, peNOS, eNOS and β-actin in siControl and si*SWELL1* mediated knockdown in HUVEC cells treated with either vehicle or 500 nM SN-401 for 96 hours (n= 6 each). **c-d.** Densitometric ratios of SWELL1/β-actin (n=12 each) (**c**) and peNOS/β-actin (n=6 each) (**d**) combined from small interfering and short hairpin mediated *SWELL1* knockdown in HUVECs treated with either vehicle or 500 nM SN-401 for 96 hours. Data are represented as mean ±SEM. Two-tailed unpaired t-test was used in **c & d**. *, ** and *** represents *p<0.05*, *p<0.01* and *p<0.001* respectively. ‘ns’ indicates the difference was not significant.

**Supplementary Fig. S10.**
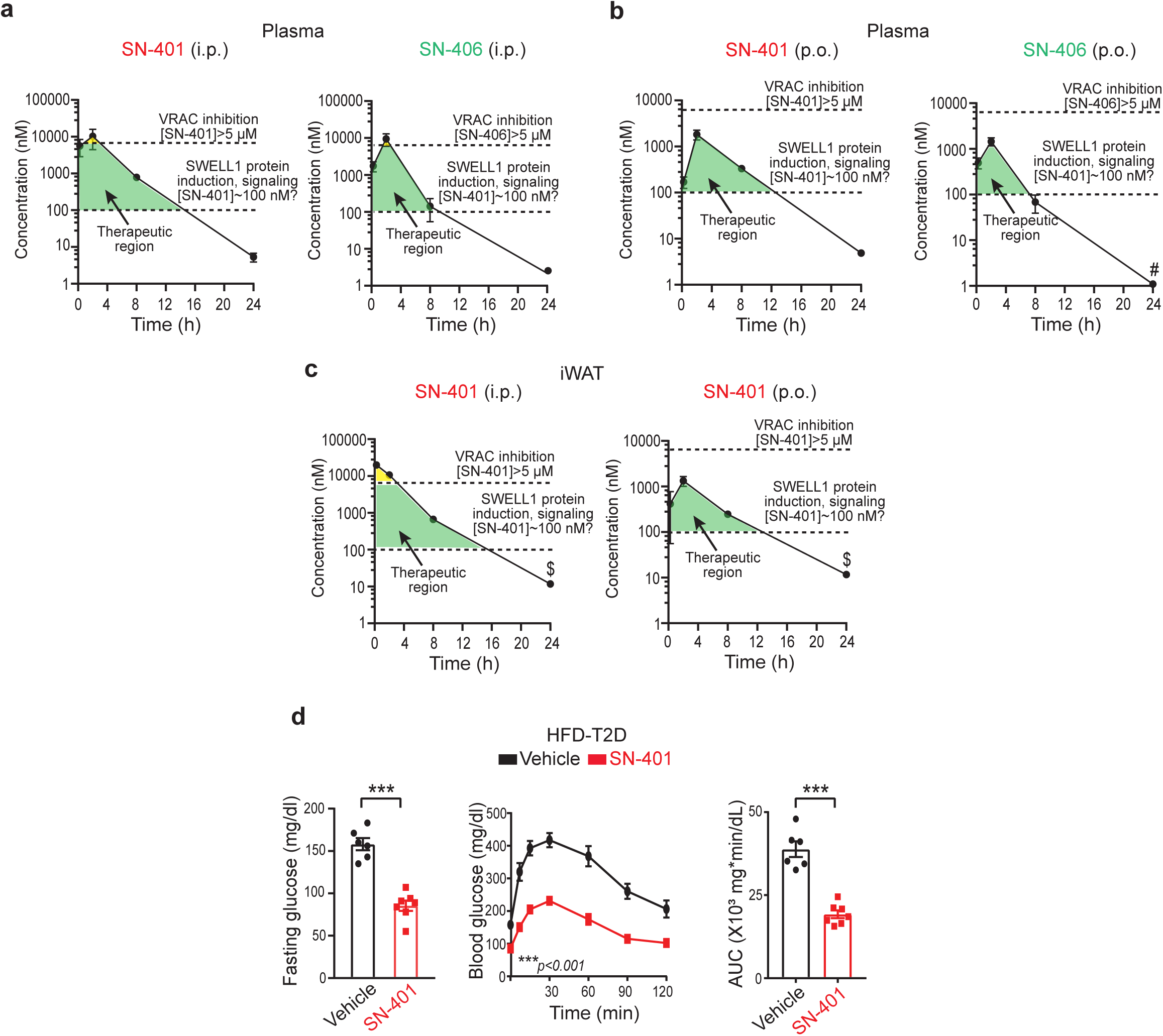
SN-40X compounds *in vivo* pharmacokinetics and oral efficacy. **a-b.** SN-401 and SN-406 *in vivo* pharmacokinetics 5 mg/kg intraperitoneally (**a**) and by oral gavage (**b**) in plasma (n=3 mice for each time point, #-below detection limit of 1 nM). **c.** SN-401 *in vivo* pharmacokinetics 5 mg/kg intraperitoneally (i.p.) and by oral gavage (p.o.) in iWAT (n=3 mice for each time point, $-below detection limit of 11 nM). **d.** Fasting glucose levels, GTT and AUC of HFD-T2D mice (10 weeks HFD) treated with either vehicle (n = 6 males) or SN-401 (5 mg/kg p.o, n = 7 males) for 5 days. Data are represented as mean ±SEM. Two-tailed unpaired t-test was used in **d** for FG and AUC. Two-way ANOVA was used for GTT in d. Statistical significance is denoted by *, ** and *** representing *p<0.05*, *p<0.01* and *p<0.001 respectively*.

## Supplementary Tables

**Supplementary Table. S1.** Characteristics of patients from whom cadaveric non-T2D and T2D islets were obtained for β-cell patch-clamp studies in Figure 1b**&d**. NA, not available.

| Patient | Age (years) | Sex | BMI | Random Glucose (mg/dl) | Estimated Glucose (mg/dl) | HbA1c (%) |
| --- | --- | --- | --- | --- | --- | --- |
| Non-T2D | 44 | F | 26.8 | 151.8 | NA | 6.1 |
|  | 57 | M | 28.7 | 144.3 | NA | 5.3 |
|  | 24 | F | 32.2 | 234 | NA | NA |
| T2D | 46 | F | 35.9 | 262.4 | NA | 6.8 |
|  | 37 | F | 38.1 | 253.8 | NA | 8.2 |
|  | 51 | M | 35.59 | NA | 157 | 7.1 |

**Supplementary Table. S2.** Characteristics of non-T2D and T2D patients from whom cadaveric islets were obtained to measure SWELL1 protein expression and GSIS assay.

| Patient | Age (years) | Sex | BMI | HbA1c (%) | Figure |
| --- | --- | --- | --- | --- | --- |
| Non-T2D | 50 | F | 31.7 | 5.7 | Fig. 1e |
|  | 61 | M | 19.6 | 5.9 | Fig. 1e |
|  | 54 | M | 26.4 | 5.1 | Fig. 1e |
|  | 31 | M | 26.2 | 5.3 | Fig. 7d,e |
|  | 31 | F | 20.3 | 4.8 | Fig. 7d,e |
|  | 66 | M | 25.6 | 4.9 | Fig. 7d,e |
| T2D | 62 | M | 25.9 | 10 | Fig. 1e |
|  | 48 | F | 30.4 | 7.5 | Fig. 1e |
|  | 54 | F | 24.4 | 7.2 | Fig. 1e |

**Supplementary Table. S3.** Characteristics of lean, non-T2D, and T2D bariatric surgery patients from whom primary adipocytes were isolated for patch-clamp studies Figure 1g. ^#^Data from lean and obese non-T2D patients were reported previously in Zhang, Y *et al*. (2017).

| Patient | Age (years) | Sex | BMI | Random Glucose (mg/dl) | Estimated Glucose (mg/dl) | HbA1c (%) |
| --- | --- | --- | --- | --- | --- | --- |
| Lean<sup>#</sup> | 52 | M | 27.56 | 97 | 111 | 5.5 |
|  | 61 | F | 28.36 | 112 | NA | 5.5 |
| Obese non-T2D<sup>#</sup> | 38 | F | 55.10 | 88 | 117 | 5.7 |
|  | 65 | F | 32.02 | 100 | 111 | 5.5 |
|  | 51 | F | 48.80 | 97 | 114 | 5.6 |
| Obese-T2D | 41 | F | 52.31 | 148 | 151 | 6.9 |

**Supplementary Table. S4.** Characteristics of lean, obese non-T2D, and obese T2D patients from whom adipose samples were obtained to measure SWELL1 protein expression levels in Figure 1h.

| Patient | Age (years) | Sex | BMI | Random Glucose (mg/dl) | Estimated Glucose (mg/dl) | HbA1c (%) |
| --- | --- | --- | --- | --- | --- | --- |
| Lean | 47 | F | 24.85 | 97 | 111 | 5.5 |
| Obese non-T2D | 36 | F | 43.05 | 106 | 114 | 5.6 |
| Obese non-T2D | 49 | M | 59.43 | 119 | 126 | 6.0 |
| Obese non-T2D | 49 | F | 41.6 | 101 (fast) | NA | 5.7 |
| Obese non-T2D | 52 | F | 49.8 | 89 (fast) | NA | 5.9 |
| Obese non-T2D | 48 | F | 50.18 | 84 | 97 | 5.0 |
| Obese non-T2D | 50 | F | 38.81 | 100 | 117 | 5.7 |
| Obese non-T2D | 43 | M | 57.65 | 105 | 105 | 5.3 |
| Obese-T2D | 57 | F | 53.69 | 273 | 183 | 8.0 |
| Obese-T2D | 53 | F | 57.21 | 122 (fast) | NA | 6.4 |
| Obese-T2D | 65 | M | 40.53 | 250 | 229 | 9.6 |
| Obese-T2D | 52 | F | 37.70 | 109 | 160 | 7.2 |

**Supplementary Table S5.** Average body weights of mice used for *in vivo* experiments in this study.

| Figure | Group | Body weight | SEM | N | Group | Body weight | SEM | N | Significance |
| --- | --- | --- | --- | --- | --- | --- | --- | --- | --- |
| Fig. 3c (GTT) | Vehicle | 41.2 | 1.59 | 7 | SN-401 | 37 | 1.85 | 7 | ns |
| Fig. 3c (ITT) | Vehicle | 40.9 | 1.51 | 7 | SN-401 | 36.2 | 1.74 | 7 | ns |
| Fig. 3e (KKA<sup>a</sup>) | Pre SN-401 | 32.9 | 1.49 | 3 | SN-401 | 30.9 | 1.96 | 3 | ns |
| Fig. 3e (KKA<sup>y</sup>) | Pre SN-401 | 45.4 | 1.03 | 6 | SN-401 | 42.5 | 1.01 | 6 | ** |
| Fig. 3f (KKA<sup>a</sup>) | Pre SN-401 | 30.7 | 2.61 | 3 | SN-401 | 30.9 | 1.80 | 3 | ns |
| Fig. 3f (KKA<sup>y</sup>) | Pre SN-401 | 42.6 | 0.73 | 6 | SN-401 | 43.4 | 1.13 | 6 | ns |
| Fig. 3h | Vehicle | 23.9 | 0.62 | 6 | SN-401 | 24.3 | 0.88 | 6 | ns |
| Fig. 3j (GTT) | Vehicle | 47.7 | 1.83 | 6 | SN-401 | 46 | 2.28 | 6 | ns |
| **Fig. 3j (ITT)** | Vehicle | 48.2 | 1.86 | 6 | SN-401 | 44.3 | 2.50 | 6 | ns |
| **Fig. 4b** | Vehicle | 38.8 | 1.0 | 7 | SN-401 | 37.5 | 0.9 | 8 | ns |
| **Fig. 4f** | Vehicle | 51.1 | 1.68 | 6 | SN-401 | 43.5 | 2.92 | 6 | * |
| **Fig. 8a** | Inactive 1 | 38 | 0.86 | 5 | SN-403 | 37.2 | 0.86 | 5 | ns |
| **Fig. 8b** | Pre SN-406 | 47.8 | 0.74 | 5 | SN-406 | 45.4 | 1.05 | 5 | * |
| **Fig. 8c** | Inactive 1 | 44.5 | 0.91 | 7 | SN-406 | 44.7 | 0.74 | 7 | ns |
| **Fig. 8f** | Inactive 1 | 38.4 | 0.85 | 5 | SN-407 | 38.7 | 1.11 | 6 | ns |
| **Suppl. Fig. 3b GTT** | Vehicle | 25.7 | 0.33 | 7 | SN-401 | 25.2 | 0.27 | 7 | ns |
| **Suppl. Fig. 3c ITT** | Vehicle | 26 | 0.31 | 7 | SN-401 | 25.4 | 0.31 | 7 | ns |
| **Suppl. Fig. 10d** | Vehicle | 40.3 | 0.77 | 6 | SN-401 | 37.2 | 0.89 | 7 | * |

**Supplementary Table. S6.** Cryo-EM data collection, processing, refinement, and modeling data for SWELL1-SN-407 in MSP1E3D1 nanodiscs for vertical and tilted poses of SN-407.^$^Information will be available upon deposition.

| **Data collection** | **LRRC8A with SN-407 (Vertical)** | **LRRC8A with SN-407 (Tilted)** |
| --- | --- | --- |
| PDB<sup>\$</sup> | XXXX | XXXX |
| EMDB<sup>\$</sup> | XXXX | XXXX |
| EMPIAR<sup>\$</sup> | XXXX | XXXX |
| Total movies # | 3576 |  |
| Selected movies # | 2968 |  |
| Magnification | 36,000 x |  |
| Voltage (KV) | 200 |  |
| Electron exposure (e<sup>-</sup>/Å<sup>2</sup>) | 51.59 |  |
| Frame # | 50 |  |
| Defocus range (um) | -0.7 to -2.2 |  |
| Super resolution pixel size (Å<sup>2</sup>) | 0.5685 |  |
| Binned pixel size (Å<sup>2</sup>) | 1.137 |  |
| **Processing** |  |  |
| Initial particle images (no.) | 936,282 |  |
| Final particle images (no.) | 85,831 | 78,324 |
| Map resolution Masked (Å, FSC = 0.143) | 3.65 (Relion) | 3.69 (Relion) |
| Symmetry imposed | C1 | C1 |
| **Refinement** |  |  |
| Model resolution (Å, FSC = 0.143 / FSC = 0.5) | 3.06/3.72 | 3.15/3.75 |
| Map-sharpening B factor (Å<sup>2</sup>) | 0 | 0 |
| Composition |  |  |
| Number of atoms | 16039 | 16309 |
| Number of protein residues | 1878 | 1878 |
| R.m.s. deviations |  |  |
| Bond lengths (Å) | 0.005 | 0.004 |
| Bond angles (Å) | 0.851 | 0.861 |
| Validation |  |  |
| MolProbity score | 1.41 | 1.43 |
| Clashscore | 3.15 | 3.81 |
| Ramachandran plot |  |  |
| Favored (%) | 95.66 | 96.15 |
| Allowed (%) | 4.34 | 3.85 |
| Disallowed (%) | 0 | 0 |
| Rotamer outliers (%) | 0 | 0.06 |

**Supplementary Table S7.** SN-401 and SN-406 *in vivo* PK parameters.

| PK parameters | SN-401 |  |  | SN-406 |  |  |
| --- | --- | --- | --- | --- | --- | --- |
|  | Oral | Intravenous | Intraperitoneal | Oral | Intravenous | Intraperitoneal |
| AUCinf (ng*h/mL) | 4682 | 5958 | 23030 | 3131 | 6532 | 18180 |
| Oral Bioavailability | 79% | NA | NA | 48% | NA | NA |
| C<sub>max</sub> (ng/mL) | 781 | 5443 | 4367 | 660.7 | 15130 | 4300 |
| T-half (h) | 2.585 | 1.428 | 2.056 | 2.058 | 0.7689 | 1.809 |

